# Loss of Mast cells and histaminergic signaling link diet to platelet-mediated NETosis and mammary cancer recurrence

**DOI:** 10.64898/2026.04.14.718388

**Authors:** Claire P. Schane, Adam T. Nelczyk, Cheng Chen, Jiyoung Seo, Yu Wang, Natalia Krawczynska, Shruti V. Bendre, Erin Weisser, Dhanya Pradeep, Hashni Epa Vidana Gamage, Lara Kockaya, Yifan Fei, Anasuya Das Gupta, Hannah Kim, Madeline Henn Bungert, Rossy Ivette Tejeda, Mei Wang, Jianping Zhao, Amar G. Chittiboyina, Ikhlas A. Khan, Jenny Drnevich, Mohammed Kadiri, Michael T. McHenry, Joy J. Chen, Liqian Ma, Sisi He, Shih-Hsuan Hsiao, Timothy M. Fan, Michael Wendt, Zeynep Madak-Erdogan, Nicki Engeseth, William G. Helferich, Erik R. Nelson

**Author notes:** These authors contributed equally to this work (Claire P. Schane, Adam T. Nelczyk, Cheng Chen). School of Medicine, Hangzhou City University, Zhejiang, China (Cheng Chen). To whom all correspondence should be addressed.

## Abstract

Breast cancer recurrence remains a clinical challenge. The period after the treatment of the primary tumor while cancer cells that evaded initial treatment lay dormant, provides a unique window of opportunity for interventions to prevent recurrence. Specific modifiable factors such as consumption of high fat diets or elevated circulating cholesterol are associated with decreased time to recurrence. Mechanistically, oxidized cholesterol and lipid species have been implicated in the regulation of the tumor microenvironment. This suggests that consumption of food prepared under oxidizing conditions such as pan-frying, may be an underappreciated risk. Using murine models of mammary cancer dormancy, we found that a diet enriched with fat from fried, cured bacon (cfBF) decreased dormancy latency times. Resulting lesions had fewer mast cells (MCs). Loss of MCs alone resulted in reemergence from dormancy. Elevated expression of a MC gene signature in breast tumors was associated with improved progression free and overall survival, highlighting the human relevance of these findings. MCs are a major source of tissue histamine, and lesions from mice fed cfBF had decreased concentrations. Importantly, antagonists of the histamine receptor 2 (H_2_R) sparked reemergence from dormancy. H_2_R antagonists are over-the-counter drugs are taken to alleviate gastroesophageal reflux disease. Chronic treatment of mice with H_2_R-antagonists sensitized platelets towards activation and crosstalk with neutrophils, and subsequent formation of neutrophil extracellular traps (NETs). The loss of platelet or NETosis activity mitigated the H_2_R-antagonist stimulated reemergence from dormancy. Therefore, we establish a novel metastatic axis which links diet to recurrence via MCs, histaminergic signaling and NETosis: Diet -- MC -- H_2_R -- (*decreased*) Platelet Activity -- (*decreased*) Neutrophil-NETosis -- (*decreased*) Reemergence from Dormancy. Our data reveal several potential intervention strategies: lifestyle, MC stabilization, histaminergic signaling, and neutrophil and platelet activity.

## INTRODUCTION

Breast cancer afflicts approximately one in eight women during their lifetime. Similar to other solid tumors, primary intervention involves surgical removal followed by courses of chemotherapy, radiation, and if warranted, targeted therapy. These strategies have dramatically increased survivorship. However, many patients develop recurrent, metastatic disease, which is much more challenging to treat. As such, breast cancer remains the second leading cause of cancer-related mortality in women^1^. Our current understanding is that metastasis involves a series of sequential steps including epithelial to mesenchymal transition, invasion and intravasation, cells in circulation, homing to specific tissues, extravasation, mesenchymal to epithelial transition, and subsequent outgrowth^2,3^. Emerging evidence suggests that the majority of breast cancer survivors continue to harbor cancer cells in distal tissues, that remain in a non-proliferative or dormant state^4–8^. In fact, up to 40% of all newly diagnosed breast cancer patients already have detectable breast cancer cells in their bone marrow^9,10^. Through mechanisms that remain unclear, dormant cells can reemerge as growing metastatic lesions. Thus, one way to have significant impact on survivorship is to develop strategies focused on preventing reemergence from this dormant stage.

Although the definitions and semantics describing dormancy are used rather fluidly in the community, dormancy can be broadly defined through two mechanisms: (1) cancer cell quiescence, where cells are not dividing or dividing slowly (for example, when oncogenic signaling is inhibited/impaired), and (2) immune-mediated dormancy, where cells are dividing but kept under control by immune surveillance. In reality, both mechanisms are likely at play simultaneously^11^. In this paper, we focus on immune-mediated dormancy.

There are several reports of inflammatory states increasing metastasis^12–14^. In seminal studies using preclinical models, the Egeblad group have characterized two ‘triggers’ of reemergence: (1) exposure to inflammatory cues (lipopolysaccharide, LPS, or inhaled cigarette smoke), and (2) chronic stress or glucocorticoid receptor signaling^15–17^. Inflammation has long been suspected to stimulate cancer; most work focuses on chronic inflammation^18^. These observations also reinforce work by the Bissell group, where pigeons infected with the Rous Sarcoma Virus only developed tumors at wounded sites^19^. Mechanistically, LPS and chronic stress were shown to stimulate neutrophils to form neutrophil extracellular traps (NETs, process termed NETosis); the presence of NETs sparking reemergence^15–17^. In related experiments, LPS also promoted recurrence through the induction of ZEB1 in cancer cells and polymorphonuclear neutrophils (PMNs) using the minimally metastatic mammary cancer lines (D2A1-d and 67NR)^20^.

Circulating cholesterol levels have been associated with poor prognosis for breast cancer patients (studies and caveats reviewed in^21,22^). On the other hand, patients taking cholesterol lowering drugs such as statins demonstrate increased time to recurrence^23–26^. Furthermore, being a carrier of a germline gain-of-function variant of *PCSK9* is associated with decreased survival amongst breast cancer patients^27^. The involvement of cholesterol in promoting metastasis was confirmed using murine models^28^. Interestingly however, while the effects of cholesterol are likely multi-faceted, an oxidized primary metabolite, 27-hydroxycholesterol (27HC) appears to be a predominant mediator. Mechanistically, 27HC works through the liver x receptors (LXRs) in myeloid immune cells, shifting them to a highly immune-suppressive phenotype^28–31^, and also stimulates extracellular vesicle biogenesis which rewires cancer cells towards a more stem-like and mesenchymal phenotype^32,33^.

Consuming a high fat diet is also associated with poor prognosis^34–37^. It is now clear that not all fats are the same in terms of risk or mechanism of action, with respect to cancer. For example, differences in saturation (monounsaturated fatty acids vs. polyunsaturated fatty acids) impart differential risks^38^. Oxidized lipids have emerged as species with significant adverse effects in terms of cancer, especially as we better appreciate lipid peroxidation and ferroptosis^39–41^. Oxidized lipids produced from myeloid cells have also been implicated in the regulation of the local tumor microenvironment^42^, as has lipid peroxides in regulatory T cells^43^. Therefore, oxidized lipids, including oxidized cholesterol species, appear to be important mediators of breast cancer progression. However, context-specific relevance, and molecular or cellular mechanisms are not well elucidated.

Here, we demonstrate that consuming a diet prepared in oxidative conditions increases mammary tumor and metastatic growth, and importantly decreases time to reemergence from dormancy. Consumption of this diet decreased mast cells and thus a loss of local histamine. Histamine, signaling through the H_2_R reduces the ability for platelets to activate and thereby induce NETosis from neutrophils, sparking recurrence.

## RESULTS

### Excess dietary cholesterol does not influence reemergence from dormancy

Retrospective studies have shown that elevated circulating cholesterol is associated with breast cancer recurrence, while statins are protective^21^. Preclinically, a high cholesterol diet increases tumor growth and metastases^28,44^. Therefore, to evaluate whether dietary cholesterol had effects on reemergence from dormancy, mice were placed on a 2% cholesterol diet. The control diet was isocaloric and matched for sodium cholate. After 8 weeks on diet, D2.0R cells were grafted. Upon intravenous graft, D2.0R cells form dormant lesions, most commonly in the lungs and bones. Metastatic lesions were monitored through time by bioluminescence imaging. No differences in metastatic burden were observed through day 111, so we administered LPS to “trigger” reemergence. While reemergence was observed (as assessed by a doubling of metastatic burden), there were no differences between mice on a normal or high cholesterol diet (**SFig. 1A**).

### Oxidized cholesterol, 27-hydroxycholesterol (27HC) reduces time to recurrence

Previous work demonstrated that oxidized cholesterol metabolites such as 27HC and 25HC were responsible for mediating many of the effects of cholesterol in terms of tumor growth and metastasis, including direct effects on cancer cells by modulating estrogen receptor activity, and suppressive effects on different immune cell populations^21,28,30–33,45,46^. Therefore, we tested the effects of increased 27HC on the D2.0R model of dormancy. Unlike cholesterol, chronic exposure to exogenous 27HC decreased the time to recurrence, even in the absence of a trigger such as LPS (**SFig. 1B**). We next utilized the Py230 primary tumor model. When grafted orthotopically, these tumors have a prolonged latency period, making them a model of dormancy. 27HC decreased the time to detection of a palpable tumor (**SFig. 1C**). To confirm these results in a system modelling clinical care, Met1 mammary-cells were grafted orthotopically and primary tumors allowed to form (∼200mm^3^), at which point they were surgically removed. Mice were then treated with three rounds of doxorubicin prior to daily treatment with 27HC. 21 days after the last doxorubicin treatment, lungs were excised and imaged for metastatic lesions. In this model of distal recurrence, 27HC increased metastatic tumor burden (**SFig. 1D**). Thus, oxidized lipids such as 27HC may be more important in terms of recurrence than their non-oxidized counterparts.

### Consumption of fat from cured fried bacon increases tumor and metastatic growth, and promotes recurrence from dormancy

Oxidized lipids appear to promote the progression of metastatic breast cancer. This would suggest that the preparation and processing of lipid-rich food may be very important in terms of the potential risk of recurrence for breast cancer survivors. Cured bacon is high in fat and cholesterol, and is commonly consumed in the USA. The average consumption of bacon in the USA is 18 pounds per person per year^47^. In the household-setting, bacon is most often prepared by pan-frying in its own fat. The curing and pan-frying process results in production of various oxidized lipid and cholesterol products, including oxysterols^48–55^. As a cured meat, it is listed by the World Health Organization as a carcinogen, although its impact on breast cancer progression is less clear.

Given that bacon is high in lipid and is commonly cured and then prepared under oxidative conditions (pan-frying), we wanted to know the impact of its consumption on tumor biology. Cured bacon and (uncured) pork belly was obtained from University of Illinois Meat Sciences Laboratory. Bacon was pan-fried and uncured pork belly was rendered in an oven and filtered through cheesecloth prior to cooling. After pan-frying, cured bacon samples were dried for 24 hours, with the dry weight being 81.99% ± 4.51%. After Soxhlet extraction, the total lipid content from dried bacon was 42.19% ± 4.41%, and 34.32% ± 2.59% from wet (undried) bacon (**SFig. 2A**).

The fatty acid composition was determined by GC-MS (standard peaks shown in **SFig. 2B** and peak identification in **S. Table 1**). Representative GC-MS traces for lard or fat from cured fried bacon fat (cfBF) are shown in **SFig. 2C**, and the breakdown of identified species listed in **S. Table 2**. The major fatty acids of fat from cured fried bacon fat (cfBF) were C18:1ω9 oleic acid (43.60% ± 0.20%), C16:0 palmitic acid (23.27% ± 0.01%), C18:2ω6 linoleic acid (11.21% ± 0.02%), and stearic acid C18:0 (10.55% ± 0.23%). Compared with rendered lard, cfBF had decreased total polyunsaturated fatty acids, increased monounsaturated fatty acids, and an increased ω6/ω3 fatty acid ratio. Total trans-fatty acids in fat from the dried bacon post-frying or cfBF and lard were similarly low. Principle component analysis comparing lipid species between lard, bacon fat and cfBF indicated that the main difference in fatty acid composition between lard and cfBF were determined by the concentration of oleic acid (C18:1) and linoleic acid (C18:2) (**SFig. 2D**).

LC-MS/MS analysis found the total cholesterol in cfBF to be 98.5 ± 13.7 mg/100g fat (**S. Table 3**). A panel of the four predominant oxysterols was run and found that the total oxysterol content was 112.3 ± 27.5 µg/100g in cfBF compared to 70.0 ± 6.5 mg/100g fat in lard, with increases in 5 of the 6 oxysterol species measured, and the major measured oxysterol species being 7α/β-HC (77.1 ± 7.4 mg/100g fat in cfBF; **S. Table 3**).

Using AIN-93G purified diet as a base, murine diets were formulated to include 5% fat from bacon or lard and 5% from soybean oil, with a control diet containing 10% fat from soybean oil. These are not considered high fat diets, and mice consuming these diets ad libitum did not show changes in weights between the diets (**SFig. 3A-B**). As expected, given its increased cholesterol content, a lard-enriched diet increased 4T1 mammary tumor growth compared to the control diet (**Fig. 1A**). However, tumors in mice consuming cfBF grew even faster, strongly supporting the contribution of oxidized lipids in tumor progression (**Fig. 1A**). Flow cytometric assessment of resulting tumors indicated a decrease in T cell infiltrate, suggesting that acquired immunity may be playing a role (**SFig. 3C**).

**Figure 1:**
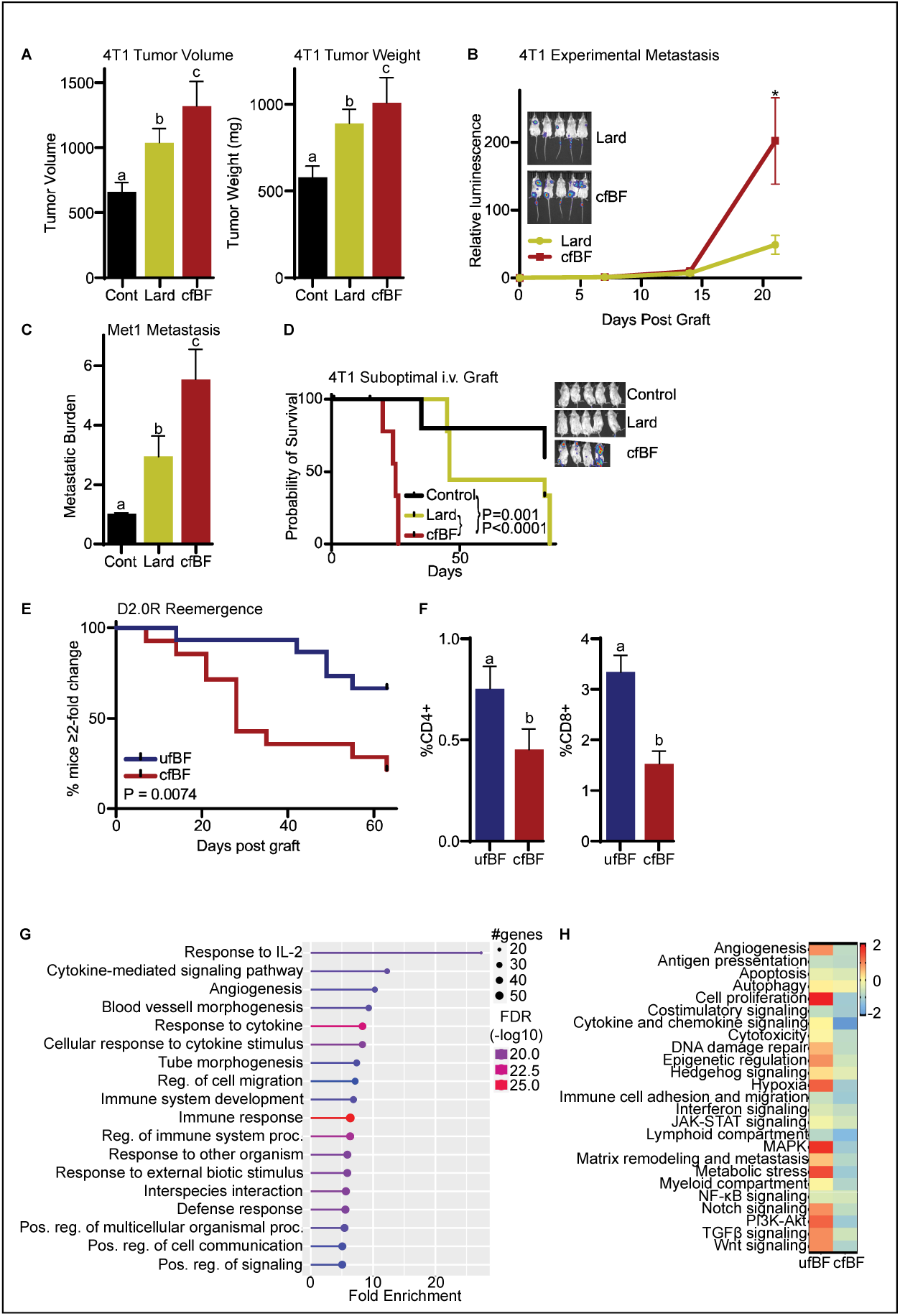
Consumption of fat from cured fried bacon increases tumor and metastatic growth, and promotes recurrence from dormancy. **(A)** 4T1 primary tumors grow at an accelerated rate in mice fed a diet enriched with pork lard or fat from cured, fried bacon (cfBF). Tumor volume on day 19 post-graft to the left of tumor weight at necropsy. Different letters denote statistically significant differences between geometric means (P<0.05, 1-Way ANOVA followed by multiple comparison test with Šidák’s correction, N=39). **(B)** 4T1 experimental metastases grew at an accelerated rate in mice fed a cfBF diet compared to a lard diet. Bioluminescence through time shown here (P<0.05, 1-Way ANOVA followed by multiple comparison test with Šidák’s correction, N=18). **(C)** Met1 experimental metastases grew at an accelerated rate in cfBF fed mice compared to lard or control diets. *Ex vivo* analyses of metastatic lungs upon necropsy. Different letters denote statistically significant differences (P<0.05, 1-Way ANOVA followed by multiple comparison test with Šidák’s correction, N=32). **(D)** To model dormancy, a suboptimal number of 4T1 cells were grafted intravenously. Mice fed a diet enriched with cfBF had significantly decreased survival times compared to those fed lard-enriched or control diets. Indicated P values are from a Logrank (Mantel-Cox) test (N=25). **(E)** Mice consuming a cfBF-enriched diet had significantly decreased time to recurrence compared to those consuming a diet enriched with uncured, fried bacon fat (ufBF), in the D2.0R model of dormancy (P value from Logrank test, N=29). **(F)** Lungs from tumor-naïve mice fed a cfBF have decreased CD4+ and CD8+ T cell infiltrate compared to ufBF. **(G)** NanoString analyses (nCounter PanCancer IO360 panel & Myeloid Innate Immunity panel) indicate consumption of a cfBF diet substantially alters the metastatic lung immune transcriptome. Gene ontology enrichment analysis of differentially expressed genes (p<0.05) was performed with ShinyGO (v0.741). Fold enrichment represents overrepresentation of genes in a pathway, FDR indicates likelihood that enrichment is by chance. **(H)** Heatmap illustrating nCounter Advanced Analysis pathway scores using normalized gene expression values (scaled using Z-transformation).

Using an experimental model of metastatic colonization and outgrowth, we found that metastatic 4T1 lesions were significantly larger in mice consuming a cfBF diet (**Fig. 1B**, **SFig. 3D**). We confirmed these results in a second model of experimental metastasis, grown in a different strain of mice (Met-1 model of triple negative breast cancer (TNBC), FVB strain compared to BALB/c for 4T1 cells); cfBF led to increased metastatic burden as assessed *ex vivo* imaging of the lungs after necropsy (**Fig. 1C**, **SFig. 3E**). Similar to 4T1 primary tumors, we observed a decrease in T cell infiltrate within the Met1-metastatic lungs of cfBF-fed mice (**SFig. 3F**).

We next adopted the 4T1 metastasis model whereby we introduced a very small number of cells intravenously. These lesions exhibit a prolonged latency period, modeling dormancy. In this model, median survival was significantly shorter in mice consuming a cfBF diet compared to control diet or lard-enriched diet (**Fig. 1D**, **SFig 4**). Thus, similar to its effects on primary tumor growth and metastasis, cfBF also stimulated reemergence from dormancy.

Whether the curing process was required for the effects of cfBF was of interest given the risk of cancer onset associated with consumption of cured meats, and curative agents acting as potential catalysts. As commercial sources of bacon are subjected to variable, and often proprietary curing techniques, performing this process “in-house” provided us with a consistent input for diet formulation. Briefly, the approach used for curing the bacon involved a combination of injecting a “cure solution” consisting of several sodium conjugates, sugar, spices and nitrites, followed by cooking and smoking (see methods). Additionally, to increase confidence that observed effect(s) were due to the curing process rather than variability of input material, bacon sourced from the same hog was utilized to create both uncured fried (uf) and cured fried (cf) bacon fat diets. Using the well-established D2.0R model of dormancy, we found that cfBF resulted in significantly shorter dormancy periods compared to ufBF (**Fig. 1E**, **SFig. 5A**). Increases were observed in total, bone-specific and lung/liver-specific metastatic burden, although these studies were underpowered to confidently assess bone (**SFig. 5B**).

To remove metastatic burden as a variable in our subsequent analyses, in a separate study the lungs of tumor-naïve mice were assessed after being on diet for 8 weeks. Under these experimental conditions, we observed a decrease in CD4+ and CD8+ T cells (**Fig. 1F**) with no significant changes in dendritic or NK cells (**SFig. 6**). Thus, cfBF results in an altered immune landscape in a tissue predisposed to metastatic spread of breast cancer.

In order to gain a better mechanistic picture of how cfBF changed the tumor microenvironment and thus stimulates recurrence, we performed Nanostring transcriptomics of the lungs (nCounter PanCancer IO360 & Myeloid Innate Immunity Panels). This revealed changes in genes associated with several different cellular pathways and signaling axes including immune response and cytokine signaling (**Fig. 1G-H**).

### Loss of mast cells is associated with recurrence of several types of cancers, including breast

Interestingly, transcriptomic (Nanostring) analysis predicted decreased mast cells (MCs) in lungs of mice fed a cfBF diet compared to ufBF, along with modest changes in other immune populations (**Fig. 2A**, **SFig. 7**). Histological examination by a veterinary pathologist who was blind to the experimental design confirmed that lungs from cfBF fed mice had fewer MCs (**Fig. 2B**). LPS is also known to induce recurrence^15^. Intriguingly, when we treated D2.0R-bearing mice with low doses of LPS sufficient to induce reemergence, we also observed a corresponding decrease in MCs (**Fig. 2C**). Thus, loss of MCs may be an upstream, generalized mechanism of recurrence from dormancy.

**Figure 2:**
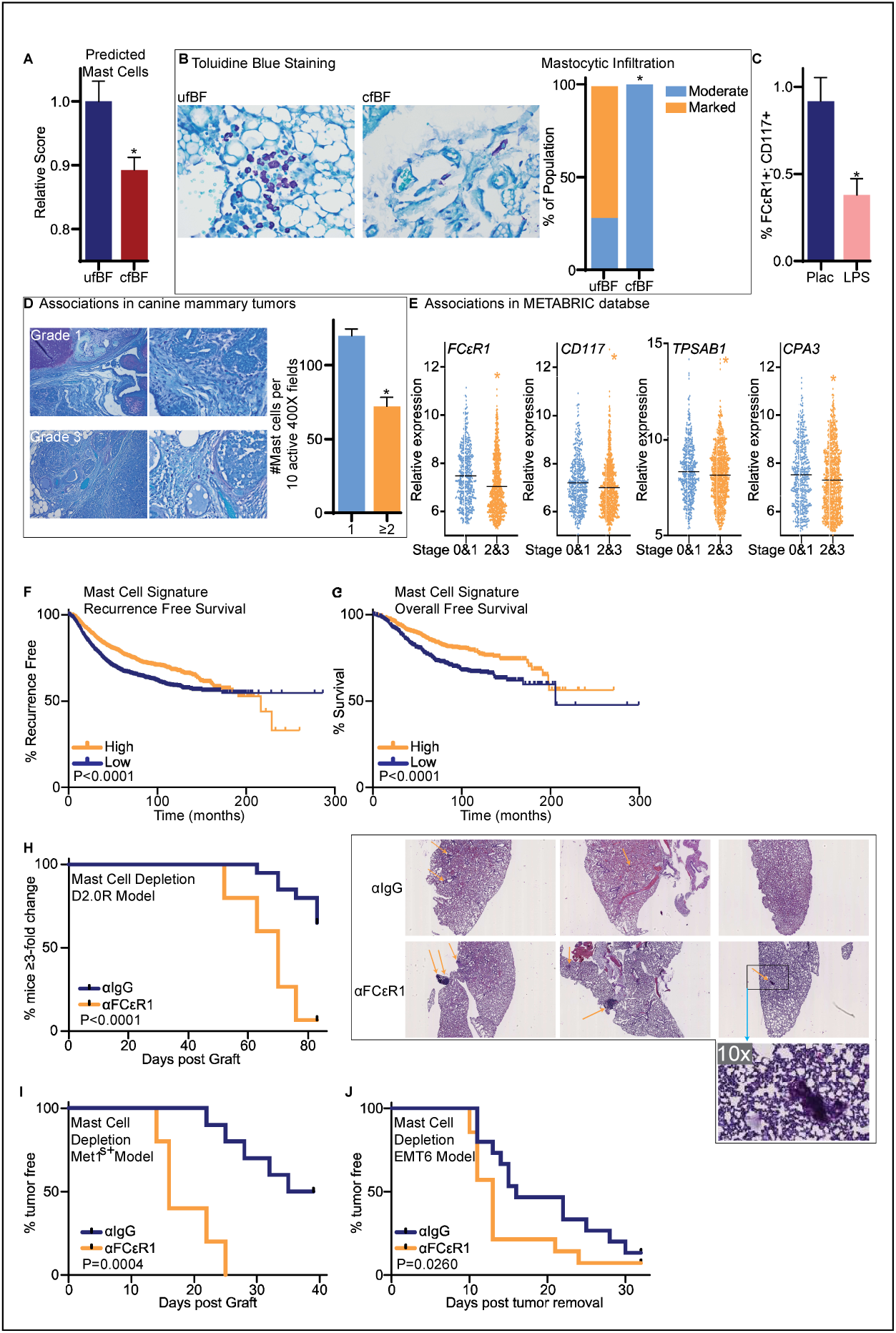
Loss of mast cells (MCs) results in recurrence from mammary cancer dormancy. **(A)** NanoString analysis of D2.0R metastases bearing mice predict a decrease in MCs in mice consuming cfBF compared to ufBF (unpaired t test, N=12). **(B)** Toluidine blue staining of lung sections from D2.0R metastases bearing mice indicate decreased MCs in cfBF fed mice. Representative micrographs to the left of data quantified by a blinded veterinary pathologist (Fisher’s exact test, N=14). **(C)** D2.0R lesion-bearing mice treated with LPS had decreased MCs as determined by flow cytometry of the lungs for FCεR1/CD117 double positive cells. **(D)** MC abundance in canine mammary tumors is decreased in higher grade tumors. Mammary tumor sections were stained with toluidine blue and assessed by a veterinary pathologist blinded to the study. Representative micrographs to the left of quantified data (unpaired t test, N=16). **(E)** Markers of MCs are decreased in human breast tumors with higher stage compared to lower. Data from the METABRIC archive, binned into tumors of stage 0&1, or 2&3 (unpaired t test, N=1358). **(F-G)** Elevated expression of a ‘MC-gene signature’ in human breast tumors is associated with increased recurrence free survival (RFS) and overall survival (OS). All breast cancer subtypes considered here. Data accessed from the Kaplan-Meier Plotter webtool based on data from GEO, EGA, and TCGA (P values indicated from Logrank test, RFS N=4890, OS N=1877). **(H)** Depletion of MCs using an antibody against FCεR1 (αFCεR1) results in decreased time to recurrence in the D2.0R model of dormancy. Survival curve to the left is based on bioluminescence (P value from Logrank test, N=35). To the right are representative micrographs of lungs with arrows denoting suspected metastatic lesions. Experimental overview and corresponding data in **SFig. 9**. **(I)** Depletion of MCs using αFCεR1 results in decreased latency using the immunogenic Met1-iRFP^S+^ line. Met1-iRFP^S+^ cells were grafted orthotopically. Mice were treated with αIgG control or αFCεR1, 3d, 4d, 6d post-graft, and then weekly. Time to detection of palpable tumor is plotted with P value from Logrank test (N=15). **(J)** The time to local recurrence of a surgically removed EMT6 mammary tumor was decreased in mice immune-depleted of MCs (P value from the Gehan Breslow Wilcoxon test, N=29).

Although we noted that DCs were also decreased in the lungs of cfBF-fed mice (**SFig. 7**), several lines of evidence suggested that this may be downstream of MCs. First, MCs have been shown to regulate DC trafficking and modulate their control of the Th1/Th2 balance^56–61^. MCs also condition DCs towards allograft tolerance^62^, and can form synapses for direct antigen transfer^63^. Finally, a major source of histamine is from MCs, and histamine has been shown to regulate DCs in many different ways, including their maturation^64–66^.

MCs are innate immune cells that reside in tissues and play critical roles in the inflammatory response and tissue homeostasis^67^. They contain granules with several mediators of the inflammatory response including histamine, which are released upon degranulation. Increased presence of MCs has been associated with both good or poor outcomes, depending on the tumor type and stage^67^. Mechanistically, MCs are largely understudied in tumor pathophysiology, although they are starting to be recognized as potential targets^68^.

Dogs (canines) develop mammary cancer. Naturally occurring canine tumors offer advantages as a model since they develop spontaneous tumors that metastasize, exhibit heterogeneity similar to human tumors, and often share an environment with their human companions^69^. Interestingly, when assessing toluidine-stained sections from archival canine mammary tumors, we found that higher grade tumors tended to have decreased MCs (**Fig. 2D**). MCs can be identified based on their high expression of FCεR1, CD117 (c-KIT), TPSAB1 (tryptase-2) and CPA3. Utilizing the METABRIC dataset, we found that decreased mRNA expression of MC-markers was a characteristic of higher stage human breast tumors (**Fig. 2E**). Likewise, *FCεR1A* was lower in higher grade tumors when assessing an independent dataset (TCGA Invasive Breast Cancer dataset, Firehose Legacy, **SFig. 8A**). Therefore, the inverse correlation of MCs to tumor stage appears to be conserved across mammalian species and suggests that MCs are protective.

We created an equally-weighted “MC-gene signature” including *FCεR1A*, *CD117*, *CPA3*, *CMA1* and *TPSAB1*. Elevated expression of this signature in breast tumors was associated with increased recurrence free- and overall- survival (**Fig. 2F-G**). This association was apparent when only *FCεR1A* was considered (**SFig. 8B**). Interestingly, the MC-gene signature was also correlated to improved progression free survival in patients taking immune checkpoint blockade (ICB, all tumor types considered, which has very few breast tumors if any in the database, **SFig. 8C**). Furthermore, elevated expression of the MC-gene signature was a good prognostic for progression free survival and overall survival in lung cancer, acute myeloid leukemia (AML) and myeloma (**SFig. 8D-I**). Thus, MCs are associated with decreased recurrence in several different cancers, both solid and blood, suggesting that they are a protective cell type with common functionality across different tumor types.

### MCs are protective against mammary cancer recurrence

Our data suggest that loss in MCs results in reemergence from dormancy. To test this relationship more specifically, we treated mice bearing dormant D2.0R lesions with an antibody against FCεR1 to immune-deplete MCs (αFCεR1, experimental overview in **SFig. 9A**). An acute exposure to this antibody resulted in significantly reduced peritoneal MCs (**SFig. 9B**). MCs within the lungs of D2.0R-bearing mice were also decreased upon necropsy at the end of the study (**SFig. 9C**). Bioluminescent imaging revealed that immune-depletion of MCs robustly decreased the time to recurrence in the D2.0R model, which was confirmed with histology (**Fig. 2H**). Flow cytometric analysis of the D2.0R bearing lungs found increases in cells with markers of myeloid derived suppressor cells: neutrophil-like CD11B+;Ly6G+ cells and monocytic-like CD11B+;Ly6C+ (**SFig. 9D-F**). Dendritic cells, NK cells, and T cells were all decreased in αFCεR1-treated mice (**SFig. 9G-J**).

While the D2.0R model is considered a good model of dormancy, the effects may be specific to this cell line. Therefore, we made use of a model of local recurrence; local recurrence being especially important for TNBC. Infrared fluorescent protein (iRFP) was overexpressed in Met1 cells, and cells super-expressing this mildly immunogenic protein were selected through multiple rounds of FACS. When grafted into the mammary fat pad, these Met1^s+^ cells exhibit a prolonged latency compared to the parental cells (median time to palpable tumor of ∼37d vs. 11d), making them a model of dormancy. MC depletion with αFcεR1 resulted in reduced latency in the Met1^s+^ model (**Fig. 2I**). In a different model of local recurrence, TNBC EMT6 tumors were surgically resected and time to local recurrence (palpable tumor) was measured. Here too, αFcεR1 resulted in reduced time to recurrence (**Fig. 2J**). Therefore, loss of MCs results in mammary cancer recurrence and by inference, MCs are protective. Given that depleting MCs alone was sufficient to alter cell abundance of other immune cells (**SFig. 9**, neutrophils, dendritic cells, natural killer cells (NKs) and T cells), it is likely that loss of MCs is an upstream event that occurs before other changes in the microenvironmental immune landscape.

### Histamine receptor H_2_ antagonists promote reemergence from dormancy

Since histamine is a major component of granules within MCs and a primary mediator of MC function, we hypothesized that reduced histamine signaling might contribute to reemergence from dormancy. Indeed, we observed decreased histamine concentrations within the metastatic lungs after exposure to all our known triggers of reemergence: mice (**1**) fed a cfBF, (**2**) whose MCs were depleted, (**3**) treated with LPS, or (**4**) treated with dexamethasone (**Fig. 3A-C**). This strongly suggests that histamine signaling may contribute to reemergence from dormancy.

**Figure 3:**
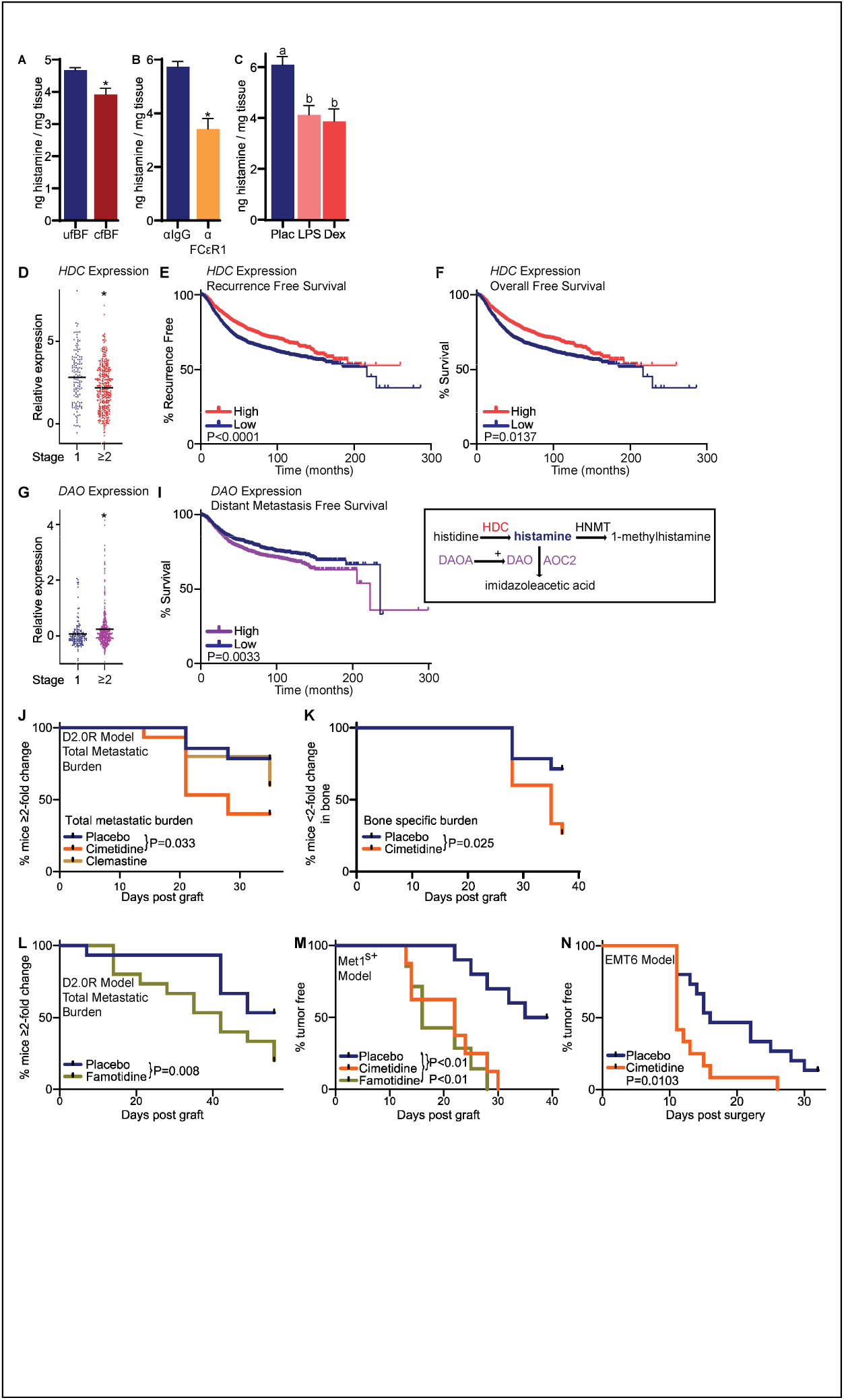
Antagonists of the histamine receptor type 2 (H_2_R) stimulate recurrence. **(A-C)** Histamine concentrations are decreased in lungs treated with “triggers” of recurrence compared to control: cfBF, immune depletion of MCs, LPS or a glucocorticoid agonist (dexamethasone) (A: unpaired T test, N=29, B: unpaired T test, N=15, C: 1-way ANOVA followed by multiple comparison test with Sidák correction, N=18). **(D)** Expression of *HDC* is decreased in higher stage breast tumors (Data from the TCGA Invasive Breast Cancer Firehose Legacy archive, binned into tumors of stage 1, or ≥ 2 (Mann Whitney test, N=498). **(E-F)** Elevated expression of *HDC* in human breast tumors is associated with increased time to recurrence and overall survival. Data accessed from the Kaplan-Meier Plotter webtool based on data from GEO, EGA, and TCGA (P values indicated from Logrank test, RFS N=4890, OS N=1877). **(G)** Expression of *DAO* is increased in higher stage breast tumors Data from the TCGA Invasive Breast Cancer Firehose Legacy archive, binned into tumors of stage 1, or ≥ 2 (Mann Whitney test, N=498). **(I)** Elevated expression of *DAO* in human breast tumors is associated with decreased time to diagnosis with distant metastasis. Data accessed from the Kaplan-Meier Plotter webtool based on data from GEO, EGA, and TCGA (P values indicated from Logrank test, N=2749). **Inset**: schematic demonstrating histamine metabolism; see text for details. **(J)** An antagonist to H_2_R (cimetidine) but not H_1_R (clemastine) stimulated reemergence from dormancy in the D2.0R model. D2.0R cells were grafted and allowed to establish dormant lesions for 6d before treatment with indicated ligands (5d/week treatment). Data shown here is based on total metastatic burden (P values from Logrank test, N=44). **(K)** The reemergence from dormancy stimulated by cimetidine was evident when only bone lesions were considered (P values from Logrank test, N=29). **(L)** A different anti-H_2_R drug, famotidine, also decreased the time to reemergence in the D2.0R model (P values from Logrank test, N=30). **(M)** Two different antagonists of H_2_R resulted in decreased latency using the immunogenic Met1-iRFP^S+^ line, a model of local recurrence (P values from Logrank test, N=15). This was part of a larger experiment, and the placebo data is the same as for Fig. 2I. **(N)** The time to local recurrence of a surgically removed EMT6 mammary tumor was decreased in mice immune-depleted of MCs (P value from Logrank test, N=27). This was part of a larger experiment, and the placebo data is the same as for Fig. 2J.

The rate limiting step of histamine synthesis is the enzyme histidine decarboxylase (HDC). *HDC* expression is decreased in higher stage breast tumors compared to lower stage ones (**Fig. 3D**). Furthermore, elevated expression of *HDC* in breast tumors was found to be associated with an increased recurrence free- and overall-survival (**Fig. 3E-F**). Correlation between *HDC* and survival were also found in lung, AML and myeloma (**SFig. 10A-C**). Notably, tumor expression of *HDC* was also associated with improved response to ICB (**SFig. 10D**).

Histamine is catabolized through two different pathways; histamine N-methyl transferase (HNMT) generates 1-methylhistamine while diamine oxidase (DAO; AOC1) generates imidazoleacetic acid. Monoamine oxidase uses 1-methylhistamine as a substrate to form methyl imidazoleacetic acid. D-amino acid oxidase activator gene (DAOA) regulates DAO. Amine oxidase, copper-containing 2 and 3 (AOC2 and AOC3) have also been reported to catabolize histamine. We found that *DAO* expression is increased in higher grade tumors (**Fig. 3G**). In support of histamine being protective, higher expression of *DAO* in breast tumors was associated with decreased time to distal metastases (**Fig. 3I**). *DAO* expression was also correlated with poor survival in lung tumors and myeloma, but not AML (**SFig. 11**). Likewise, the activator of DAO, *DAOA* expression was associated with worse survival in breast and lung cancer (P=0.0521 and 0.0664 respectively) (**SFig. 11**). There were insufficient samples to make firm conclusions regarding a correlation between *DAOA* and survival in AML or myeloma (**SFig. 11**). AOC2 expression was associated with worse prognosis in breast and lung tumors, but not AML or myeloma (**S. Fig. 11**). Collectively, these results strongly suggest that histamine abundance is associated with reduced risk of recurrence.

There are four different histamine receptors, H_1_-H_4_R^70^. H_1_R is well known for its role in promoting the allergic response. H_2_R mediates the histamine increase in gastric acid secretion. H_3_R and H_4_R are high affinity receptors expressed predominantly in the brain and immune cells respectively^70,71^. Similar to MCs, histamine and its receptors are implicated as both pro- or anti-cancer, likely highly context dependent^72,73^.

In order to directly test if loss of histamine signaling promotes reemergence from dormancy, and to glean insight into receptor-based mechanisms, we treated D2.0R grafted mice with two common antihistamines, an H_1_R antagonist (clemastine), or an H_2_R antagonist (cimetidine). D2.0R cells were grafted 6 days prior to the start of daily treatment with these ligands. Intriguingly, the H_2_R antagonist, but not an H_1_R antagonist broke dormancy (**Fig. 3J**). Interestingly, we found that the effects of cimetidine were also apparent when bone lesions were considered independently (**Fig. 3K**). This is important given the predominance of bone metastasis in breast cancer patients. An independent experiment evaluating the effects of a second H_2_R antagonist (famotidine) found similar results (**Fig. 3L**). Both cimetidine and famotidine also reduced latency time in the immunogenic Met1^s+^ model (**Fig. 3M**). Finally, mice treated with cimetidine developed a higher metastatic burden after surgical resection of EMT6 tumors (**Fig. 3N**), further suggesting that blocking H_2_R results in reemergence from dormancy and is a conserved mechanism across models and tissues.

We found that both H_1_R and H_2_R expression is decreased through time in metastatic 4T1 lesions, while H_3_R and H_4_R expression did not significantly change (**SFig. 12**). This suggests that H_2_R signaling is selected against as tumors progress. Collectively, correlational data from human tumors and our preclinical evidence strongly implicate H_2_R activity as being protective in that it maintains dormancy and/or prolongs latency.

### T cells are required for recurrence-promoting effects of anti-H_2_R drugs in an indirect manner

To start elucidating the mechanism by which H_2_R blockade stimulates recurrence, we first ruled out direct effects of anti-H_2_R ligands on the proliferation of cancer cells themselves (**SFig. 13**). Thus, blockade of H_2_R in cells extrinsic to cancer cells was likely what drives recurrence. RNAseq analysis revealed 1693 differentially regulated genes (DEGs) in D2.0R metastatic lungs from cimetidine treated mice compared to control (**Fig. 4A**). Enrichment of cancer-associated pathway genes was noted in cimetidine-treated lungs, including PI3K-AKT, MAPK, RAS, and chemical carcinogenesis pathways (**Fig. 4B**). Enrichment of these pathways would be expected as cimetidine increased metastatic burden/progression. Downregulated pathways included several involving immune-mediated pathways, including Th17, Th1 and Th2 differentiation, T cell receptor signaling, B cell receptor signaling, PDL1 and PD1 checkpoint pathway, and the NFκB pathway (**Fig. 4C**). Similarly, gene set enrichment analyses (GSEA) revealed several immune-pathways to be suppressed (**Fig. 4D**)

**Figure 4:**
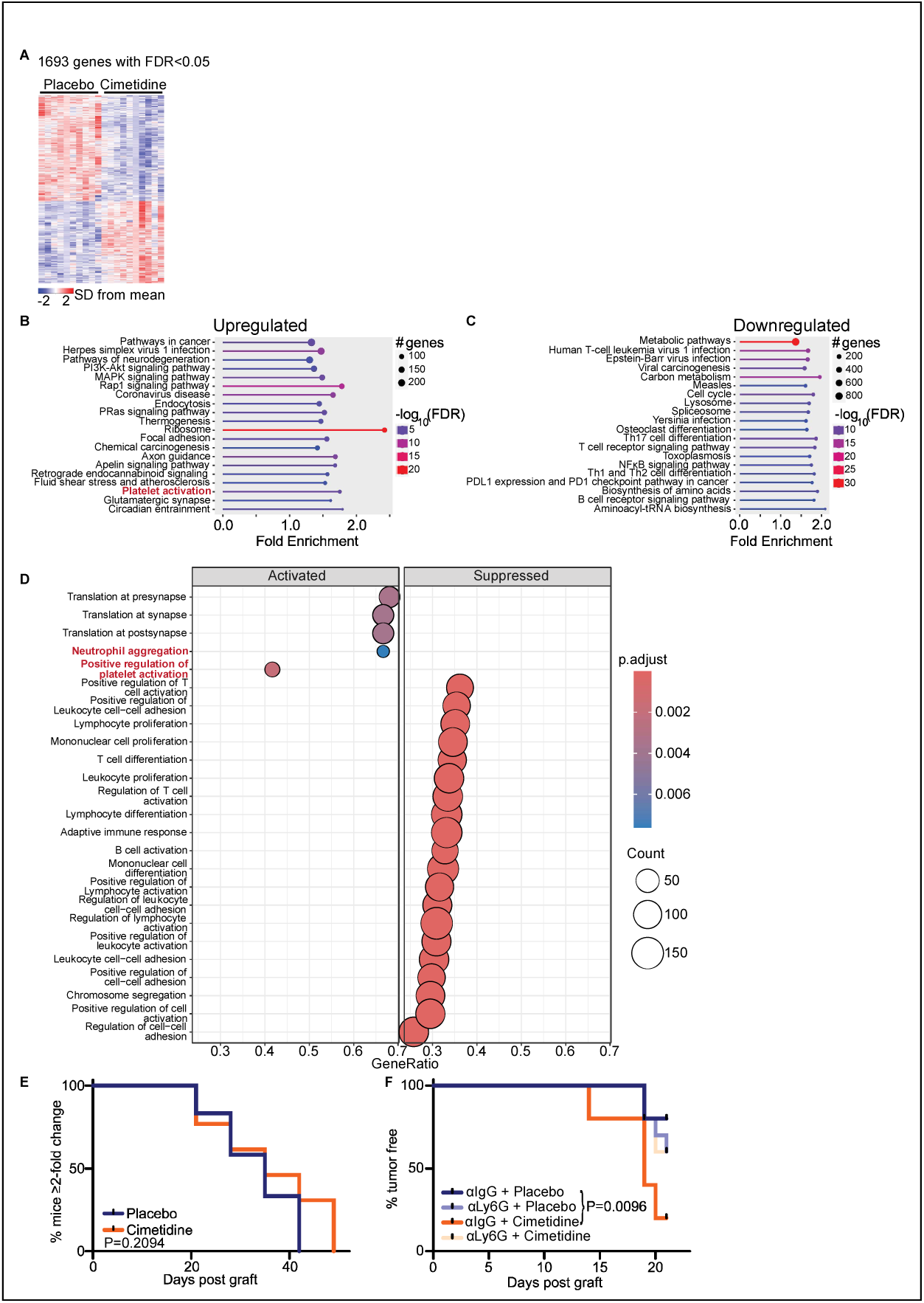
T cells and neutrophils are required for the reemergence of dormancy stimulated by H_2_R antagonists. **(A)** RNA-seq was performed on D2.0R-bearing lungs from mice treated with placebo or cimetidine (samples from Fig. 3J). Heatmap of differentially expressed genes (DEGs) shown here. Each column represents a single sample, from an individual mouse (ie: biological replicate). **(B)** Gene ontology (GO) analysis for enrichment of upregulated genes (FDR of P≤0.01). **(C)** GO analysis for enrichment of downregulated genes (FDR of P≤0.01). **(D)** Gene set enrichment analyses (GSEA) was performed and the top enriched pathways are indicated here. **(E)** Anti-H_2_R ligand, cimetidine, does not influence recurrence from dormancy in mice lacking mature T cells (nu/nu mice). Nu/nu mice were grafted with D2.0R cells prior to daily treatment with placebo or cimetidine. **(F)** Anti-H_2_R ligand, cimetidine, does not influence recurrence from dormancy in mice depleted of neutrophils with an antibody against Ly6G (αLy6G). Mice were grafted with Met1^s+^ cells prior to treatment initiation as indicated.

Up to this point, our data suggested three different downstream cellular pathways may be mediating the stimulus for recurrence: (A) decreased NK cell signature in metastatic lungs of cfBF-fed mice, (B) decreased DCs in metastatic lungs of cfBF-fed or anti-H_2_R treated mice, and (C) decreased T cells in cfBF-fed or anti-H_2_R treated mice as evidenced by RNAseq and flow cytometry (**Fig. 1G, 4B-C, SFig. 7** and **SFig. 9**). We subsequently assessed the effects of anti-H_2_R ligands on these cell types.

Single cell RNA-seq (scRNAseq) of circulating immune cells indicated that H_2_R expression is relatively higher in myeloid subtypes compared to lymphocytes and NKs (**SFig. 14A-C**; data mined from the Human Protein Atlas^74^). Likewise, scRNAseq analysis of human or mouse mammary tumors found relatively higher expression of H_2_R in cells of the myeloid immune lineage compared to lymphocytes or other cell types, with very little expression in the cancer cells themselves (**SFig. 14D-G**). It should be noted that the tissue processing and cell-preparation for scRNAseq often leads to the loss of neutrophils, basophils, MCs and platelets, resulting in their underrepresentation. qPCR analysis of cultured murine immune cells found H_2_R was expressed across all cells assessed, with relatively higher concentration in neutrophils, DCs, MCs and platelets, compared to T cells or macrophages (**SFig. 14H**).

### NK cells are not directly involved in mediating anti-H2R response

Treatment of murine NK cells with histamine, cimetidine or famotidine did not significantly or consistently alter viability of NK cells (**SFig. 15A**). Likewise, markers of NK-activation were not altered by treatment with anti-H_2_R ligands (**SFig. 15B**).

### T cells are not directly involved in mediating anti-H2R response

T cell viability was not significantly altered when directly treated with different histamine receptor ligands (if anything, increased by anti-H_2_R ligands; **SFig. 16A**). Expansion of T cells treated with anti-H_2_R ligands was not altered after activation with antibodies against CD3 and CD28, regardless of T cell type (pan, CD4+, Treg, or CD8+; **SFig. 16B**). Given their highly immune-suppressive role and associations with poor prognosis, this experiment was repeated under culturing conditions to promote Treg differentiation; cimetidine nor famotidine altered T cell expansion under these conditions (**SFig. 16C**). Thus, it was unlikely that effector cells such as NK or T cells were the primary target for anti-H_2_R drugs, at least in terms of recurrence.

### Myeloid immune cells are not directly involved in mediating anti-H2R response

Bone marrow derived macrophages did not show altered viability or expression of genes associated with activation, with the exception of *Selplg*, which showed upregulation after treatment with anti-H_2_Rs, an anti-H_1_R or histamine itself **(SFig 17A-B)**. Macrophages are known to support T cell expansion. To assess this, we pre-treated macrophages and then co-cultured them with activated T cells. We observed no changes in T cell expansion, regardless of whether we considered pan T cells, or just CD4+, or CD8+ cells (**SFig. 17C**). DCs have relatively high H_2_R expression (**SFig. 14**). However, treatment of DCs with antagonists to H_2_R did not result in altered viability or markers of activity (**SFig. 18A-B**). No changes were observed in the expansion of different T cell subtypes after co-culture with DCs pretreated with anti-H_2_R ligands (**SFig. 18C**). The resulting expanded T cells also showed no differences in IL4, IL17 or IFNγ (**SFig. 18D**).

MCs themselves express H_2_R. However, treating MCs with cimetidine or famotidine had no major effects on MC viability or expression of genes associated with their activation (**SFig 19A-B**). Likewise, H_2_R ligands had no significant effects on MC degranulation, either basal or in the presence of a degranulating activator C48/80 (**SFig. 19C**). Although MCs are not traditionally thought of as supporting T cell activity, they do have the capacity to present antigen as well as secrete various cytokines^75,76^. However, anti-H_2_R ligands had no effect on the ability of MCs to present an ovalbumin antigen and activate T cells engineered to recognize ovalbumin peptides (OTI and OTII system, **SFig. 19D-E**). It was also possible that MCs were acting on T cells by modulating DC activity. However, when MCs and DCs were co-cultured in the presence of different H_2_R ligands, and then cultured with activated T cells, there were no differences observed in pan, CD4+ or CD8+ T cell expansion (**SFig. 19F**).

Neutrophils also express H_2_R, and we did find increased Ly6G^+^ granulocytes in metastatic lungs depleted of MCs (**SFig. 9D-F**). However, treatment of neutrophils with cimetidine or famotidine resulted in similar viability compared to vehicle, and did not significantly alter the expression of genes associated with activity (**SFig. 20**).

Although we found no direct effects of H_2_R antagonists on T cell expansion, nor their expansion when co-cultured with macrophages, DCs or MCs pretreated with these drugs (**SFigs. 16-19**), transcriptomic evidence still suggested their involvement (**Figs. 1, 4, SFig. 7**). To test this more directly, we evaluated the effect of cimetidine on D2.0R recurrence in athymic (nu/nu) mice which lack functional T cells. When these mice were treated with cimetidine, there were no differences in recurrence, confirming that T cells are required for its pro-recurrence effects (**Fig. 4E**). Therefore, H_2_R antagonists break dormancy in a manner that requires T cells, but in an indirect manner.

### Neutrophils required for recurrence stimulated by antagonists to H_2_R

Neutrophils have been implicated in the process of ‘breaking’ cancer dormancy. There are several reports of inflammatory states increasing metastasis^12–14^. Two ‘triggers’ of reemergence from immune-mediated dormancy have been previously described: (1) exposure to inflammatory cues (LPS or inhaled cigarette smoke), and (2) chronic stress or glucocorticoid receptor signaling^15–17^. These triggers stimulate neutrophils to form neutrophil extracellular traps (NETs, process termed NETosis); the presence of NETs sparking reemergence. LPS also promoted recurrence through the induction of ZEB1 in cancer cells and neutrophils using the minimally metastatic mammary cancer lines (D2A1-d and 67NR)^20^.

Although anti-H_2_R drugs had no effects on neutrophil viability or markers of activation (**SFig. 19**), we had observed increased presence when MCs were immune-depleted (**SFig. 9D**). Furthermore, neutrophil aggregation was an activated pathway, as identified by GSEA (**Fig. 4D**). Thus, in order to test whether neutrophils were required for the breaking of recurrence observed upon anti-H_2_R treatment, we immune depleted these cells using an antibody against Ly6G^28^. As expected, cimetidine treatment of Met1^s+^ grafted mice resulted in decreased latency time (**Fig. 4F**). However, cimetidine had no effects when mice were depleted of neutrophils (**Fig. 4F**). Therefore, although cimetidine had no obvious effects on the expression of genes associated with neutrophil function, they are required for its effects on latency/recurrence.

### H_2_R antagonists stimulate mammary cancer recurrence by sensitizing platelets to activation, which then induce NETs from neutrophils

Given that we did not find changes in the expression of genes associated with other cancer-related functions of neutrophils (**SFig. 20**), we next assessed whether anti-H_2_Rs influenced NETosis, a process that would not necessarily result in altered gene expression. However, direct treatment of murine, bone marrow derived neutrophils with cimetidine or famotidine did not alter NETosis (**Fig. 5A**). These data suggested that H_2_R antagonists might not have direct effects on neutrophils but did not rule out the possibility of an intermediate cell type with the capacity to induce neutrophil NETosis.

**Figure 5:**
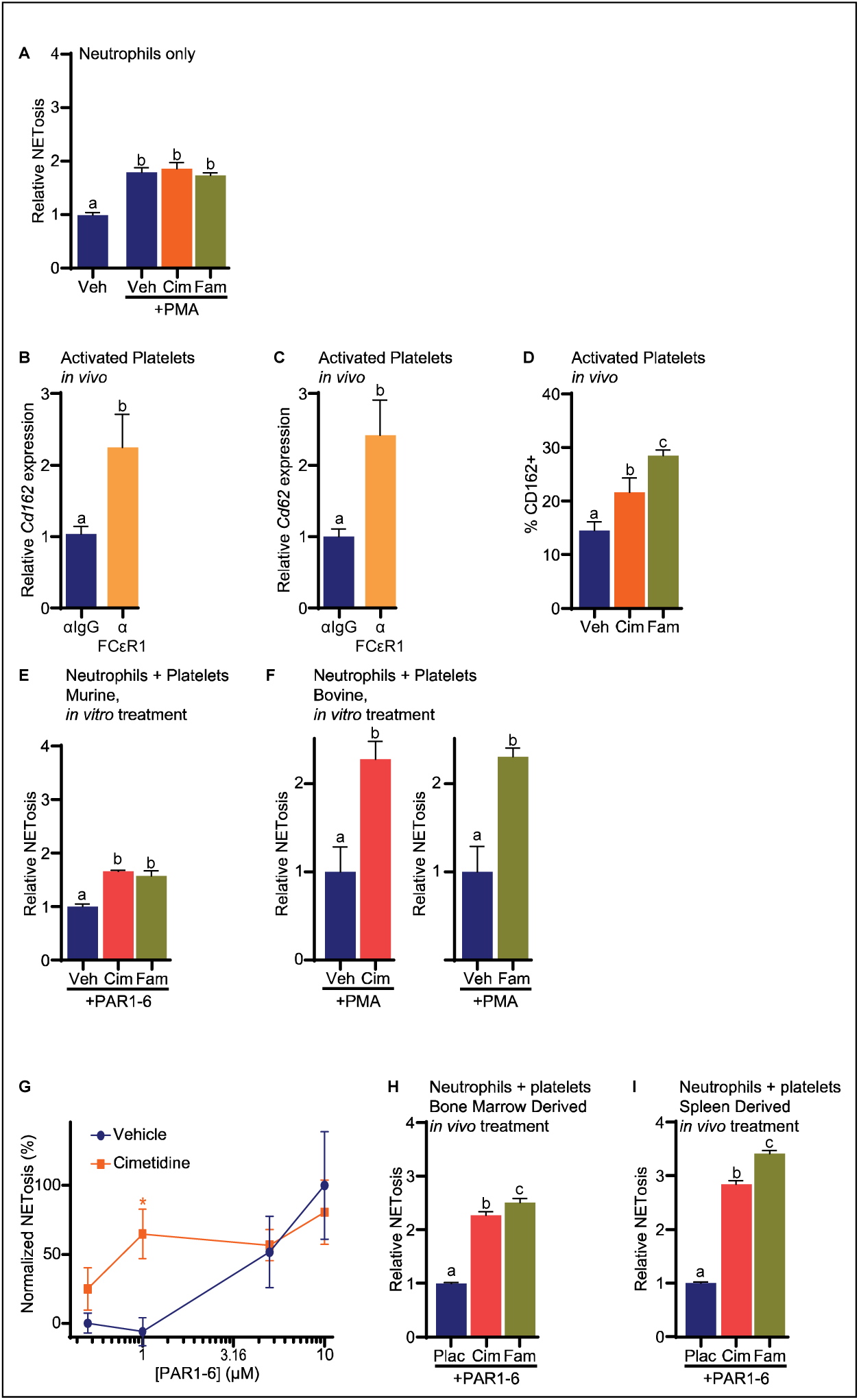
H_2_R antagonists sensitize platelets towards activation, which then stimulate NETosis and subsequent recurrence. **(A)** Neutrophils co-treated with PMA and either cimetidine or famotidine and stained with SYTOX Green continue to undergo NETosis at the same rate compared to control, when induced by PMA. Relative NETosis was analyzed by Flow Cytometry by number of positive SYTOX Green cells.N=15-21. **(B)** D2.0R-bearing lungs of mice depleted of MCs (αFCεR1) show increased *Cd162* (*Selplg*) and **(C)** Cd62 (*Selp*) expression (unpaired T test, N=35). **(D)** Lungs from tumor-naïve mice that were chronically treated with cimetidine or famotidine had increased presence of activated platelets (CD162+ events), as assessed by flow cytometry (1-Way ANOVA followed by multiple comparison test with Šidák’s correction, N=43). **(E)** NETosis after co-culture of murine neutrophils and platelets, treated with vehicle, cimetidine or famotidine. Platelets were activated with a sub-optimal dose of PAR1-6 (thrombin receptor 1 activator peptide 6) (1-Way ANOVA followed by multiple comparison test with Šidák’s correction, N=12). **(F)** NETosis after co-culture of bovine neutrophils and platelets, treated with vehicle, cimetidine or famotidine. Platelets were activated with a sub-optimal dose of either PMA (unpaired T test, Left panel N=10, Right panel N=9). **(G)** Cimetidine sensitizes platelets towards activating NETosis. Platelets and neutrophils were co-cultured in increasing doses of PAR1-6 the presence of either vehicle or cimetidine, and resulting NETosis was measured. NETosis was normalized to vehicle means at 0.5µM and 10µM. Asterisks indicate statistical difference (2-Way ANOVA followed by multiple comparison test with Šidák’s correction, N=119). **(H)** NETosis in a neutrophil-platelet co-culture after chronic in vivo treatment of mice with placebo, cimetidine or famotidine. Mice were treated with indicated ligands for 49 days before neutrophils and platelets were harvested from the bone marrow, or **(I)** spleen (1-Way ANOVA followed by multiple comparison test with Šidák’s correction, N=33).

In this regard, gene ontology analysis of RNAseq from lungs of D2.0R mice indicated that cimetidine resulted in platelet activation (**Fig. 4B-C**). Likewise, GSEA identified neutrophil aggregation and positive regulation of platelet activation as enriched (**Fig. 4D**). In support of this, we observed an increase in markers of platelet activation including *Selp* (p-selectin/CD62P) and its ligand *Selplg* (CD162) after MC depletion (**Fig. 5B-C**). In a separate experiment tumor-naïve mice were chronically treated with cimetidine or famotidine. Flow cytometry of lungs from these mice treated with either anti-H_2_R ligand, had increased CD162^+^ events (ie: activated platelets) (**Fig. 5D**). Events positive for other platelet-associated markers, CD41a and CD61, were also increased, especially in famotidine-treated mice (**Supplementary Fig. 21**).

Platelets have long been implicated in cancer metastasis including breast cancer^77^. They have been associated with epithelial to mesenchymal transition (EMT), protecting cancer cells in circulation, increasing adhesion of cancer cells to endothelial cells and promoting extravasation, and neo-angiogenesis^78–85^. Interestingly, recent evidence suggests an interplay between platelets and NETosis, whereby activated platelets can induce neutrophil-NETosis^86^. Extruded NETs can trap and block other immune cells such as NK and T cells from gaining access to tumors^87^.

Therefore, we tested the hypothesis that H_2_R antagonists worked through platelets to induce NETosis. When murine platelets were co-cultured with neutrophils, we observed increased NETs when treated with either cimetidine or famotidine in the presence of the thrombin mimetic PAR1-6 (**Fig. 5E**). To confirm that this effect was not specific to mice, we tested the effects of cimetidine and famotidine on bovine cells. As expected, both of these anti-H_2_R ligands resulted in increased NETosis when included in the co-culture of bovine platelets and neutrophils (**Fig. 5F**). Thus, the effects of H_2_R antagonists on platelet-neutrophil co-cultures is a mechanism conserved across species.

To better evaluate whether H_2_R-antagonists sensitize platelets towards activating neutrophil NETosis, co-cultures were treated with either vehicle or cimetidine with increasing concentrations of PAR1-6. Intriguingly, cimetidine robustly increased the ability of low doses of PAR1-6 to stimulate NETosis, whereas PAR1-6 alone had no effect at these doses (**Fig. 5G**).

Finally, to establish the physiological relevance of these *in vitro* findings, we treated mice chronically with either cimetidine or famotidine for 49 days. When platelets and neutrophils were harvested and co-cultured, NETosis was increased upon treatment with H_2_R antagonists, regardless of whether we utilized cells isolated from the bone marrow or spleen, or whether we assessed NETosis via extruded DNA or MPO (**Fig. 5H-I**, **SFig. 22**). In fact, the NETosis observed after chronic *in vivo* treatment was even greater than when naïve cells were co-cultured (**Fig. 5E** vs. **H&I**). Taken together, these data support a model whereby H_2_R antagonists sensitize platelets towards activation, even under conditions with very low levels of traditional activators (ie: thrombin).

### Platelets and NETs required for effects of H_2_R antagonist on reemergence from dormancy

Collectively, our data support a model whereby anti-H_2_R drugs sensitize platelets towards activation, which then stimulate NETosis from neutrophils, thus inducing recurrence. To test this model, we treated D2.0R bearing mice with cimetidine in combination with either heparin to inhibit thrombin-activation of platelets or DNAse to digest NETs. Cimetidine alone stimulated recurrence, as we expected (**Fig.6**). However, co-administration of either heparin or DNAse attenuated this response (**Fig. 6**). Therefore, H_2_R antagonists likely induce recurrence through their sensitization of platelets to activation and subsequent NETosis from neutrophils. Since T cells are also required for the effects of H_2_R antagonists (**Fig. 4E**), it is likely that NETs act to exclude access of T cells to dormant lesions, allowing the cancer cells to escape immune control (**Fig. 7**).

**Figure 6:**
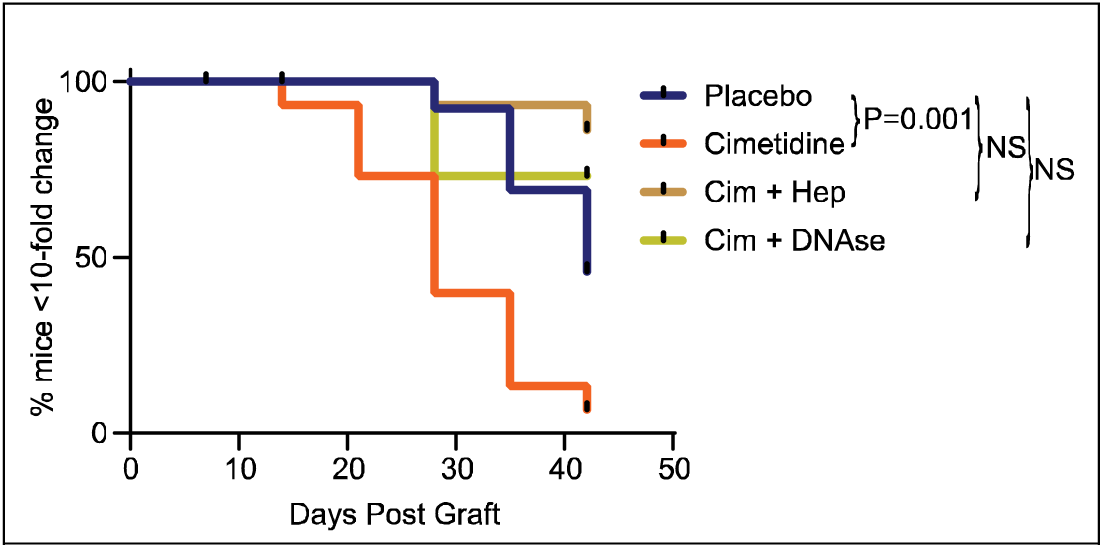
Effects of H_2_R antagonists on reemergence from dormancy requires activation of platelets and NETs. D2.0R lesion bearing mice were treated with placebo or cimetidine alone or in combination with either heparin to prevent platelet activation, or DNAse to reduce NETs. Either heparin or DNAse prolonged cimetidine-mediated recurrence (P value from Logrank test, N=60).

**Figure 7:**
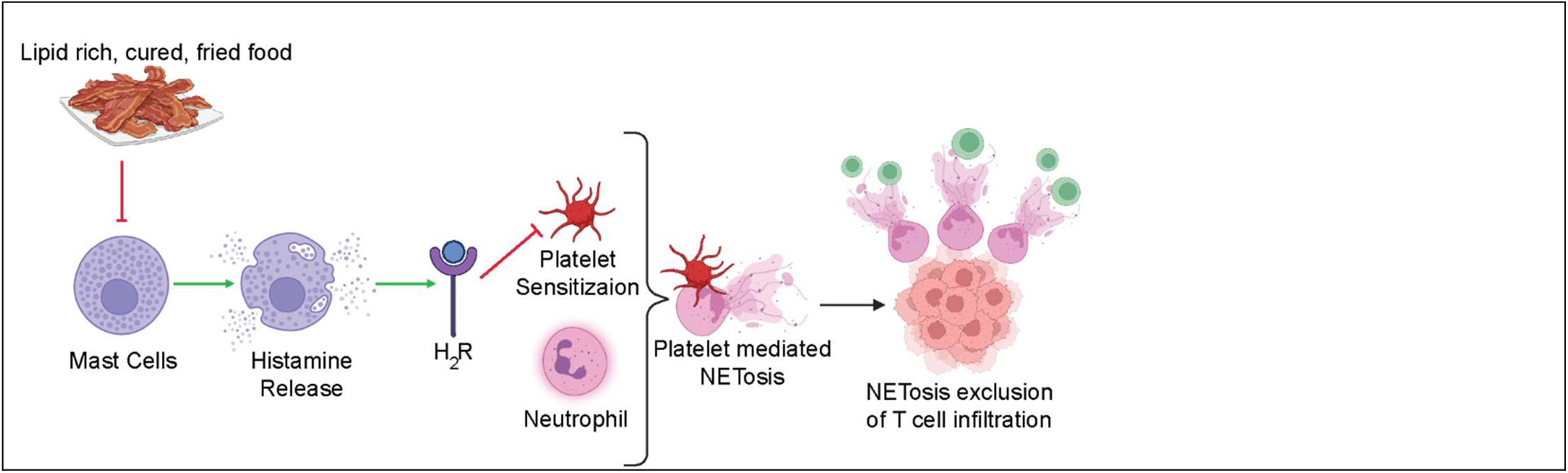
Schematic overview of proposed model. Consumption of cured, fried bacon fat results in the loss of mast cells and thus, local histamine. Histamine, normally signaling through the H_2_R in platelets reduces their likelihood of activation; anti-H_2_R ligands prime platelets towards activation. Activated platelets then stimulate NETosis from neutrophils. The platelet/NETs exclude immune cell infiltration (DCs, T cells, NK cells), allowing lesions to escape immune surveillance and recur.

## DISCUSSION

The time from the cessation of primary intervention to recurrence of breast cancer provides a period uniquely positioned for preventative intervention strategies. Although there are many factors associated with recurrence, such as obesity and elevated cholesterol, cellular and molecular mechanisms remain diffuse and/or poorly understood. Dormancy itself likely has two broad mechanisms: (1) cancer cell quiescence or slowly dividing cancer stem-like cells, and (2) tumor mass dormancy, where a slowly dividing population of cells are kept at low levels by active immune surveillance^88^.

Neutrophils have been implicated in breast cancer progression and metastatic recurrence. Some positive attributes of neutrophils have been described, although the majority of studies suggest a pro-tumor role. These different reports are likely due to the various functional states neutrophils can adopt; similar to other myeloid immune cells such as macrophages, neutrophils can be polarized into anti-pathogen N1 and wound healing N2 polarities. In preclinical models of mammary cancer metastasis, γδ T cells secrete IL17 resulting in a systemic, G-CSF-dependent expansion of neutrophils that facilitate the establishment of a favorable metastatic niche^89^. Interestingly, 27HC promotes metastasis by increasing neutrophil and γδ T cell biomass at the lung metastatic site, shifting neutrophils towards ‘N2’, and generally impairing the ability of these myeloid cells to support T cell expansion^28,31^. Through a ROS-dependent mechanism, 27HC also increases neutrophil secretion of extracellular vesicles, which act directly on cancer cells, shifting them towards a more stem-like and mesenchymal-like phenotype^32,33,45^. The resulting cancer cells have increased WNT signaling, which allows them to survive in an anchorage-independent way, have a lower proliferation rate, are more migratory and are resistant to cytotoxic chemotherapy ^32,33,45^. Cholesterol itself can activate Dvl2, although whether oxysterols can do this, or whether this occurs in neutrophils is not yet known^90^.

Neutrophil NETosis and subsequent NETs are known to assist in tumor growth and ‘spark’ metastatic recurrence. Interestingly, activated platelets have recently been shown to stimulate NETosis^86^. Furthermore, by promoting accumulation of iron, platelets can serve as a stimulus for lipid peroxidation, ultimately resulting in NETosis^91^. Similarly, heme-induced platelet activation mediates platelet-neutrophil aggregates and NETosis^92^. Platelet priming by Willebrand factor induces integrin αIIbβ3 activation enhancing neutrophil binding, ultimately inducing NETs through mechanosensitive-dependent mechanisms^93^. This results in NETs which shelter cancer cells from immune surveillance, allowing them to establish independent immune-evasion tactics, and ultimately developing into overt metastatic lesions^87^. In preclinical models, anticoagulants appear to increase the efficacy of ICB, although retrospective analyses of clinical data yields a more complex picture^94^.

We have demonstrated that consumption of a diet enriched with oxidized lipids depleted MCs resulting in a loss of histamine. Reduced H_2_R signaling results in sensitization of platelets towards activation, subsequent neutrophil NETosis and ultimately, mammary cancer recurrence (**Fig. 7**). This highlights an elegant pathway that links diet to both the innate and acquired immune systems, implicating MCs as an upstream mediator.

The role of MCs in tumor progression is controversial, likely stemming from our poor understanding of their biology and functional states^95^. In fact, MCs in general have long proven controversial, especially around early evidence that they are *bona fide* antigen presenting cells; MC antigen presentation described in 1989^75^ with other papers contesting this^96^, but then redescribed for presenting antigen to certain T cell subpopulations or under specific conditions different times^97–99^, and finally, a phase II trial showing that the stabilization of MCs increases their ability to present antigen, successfully improving the efficacy of αPD1 in patients with ICB-refractory TNBC^100^.

Studies investigating MCs and breast cancer are few, with those that are published indicating a complex ‘dual-role’ – as in, MCs being reported as being both a favorable and poor prognostic^95,101,102^. Intratumoral MC density was not associated with survival in a small study of individuals with invasive breast carcinoma^103^. However, in a larger study of 4,444 invasive breast tumors, stromal MCs were associated with increased survival^104^, a similar survival benefit now described in several other reports^105,106^. One limitation of these association studies is that they generally fail to account for different functional states of MCs. At least two major states of MCs have been described for humans: MC_T_ and MC_TC_, both of which are likely to be polarized across a spectrum, similar to other myeloid immune cells^107–109^. Regardless, our analysis of genes selectively expressed in MCs suggests a strong association between these cells and increased survival time, reinforced by findings that MC infiltration is inversely associated with tumor grade/stage in both humans and dogs (**Fig. 2** and **SFig. 8**). The survival benefit was observed across several cancer types. Interestingly, MC infiltration in human breast tumors was also associated with T cell density^110^, perhaps due to a decrease in NETs, as our data would suggest. MCs have also been implicated in regulating angiogenesis^101^, supported by clinical correlations between MCs and microvascular density^111^.

In our preclinical models, depletion of MCs resulted in quicker recurrence (**Fig. 2**), supporting the protective nature of MCs. However, a recent study using humanized mice and a melanoma model, has found that infiltrating MCs are increased after treatment with ICB, whereas treatment with sunitinib or imatinib, to inhibit CD117, improved the efficacy of αPD1^112^. The discrepancy between our findings and these could be because (A) neither sunitinib nor imatinib are specific to CD117, and likely have direct effects on cancer cells^113,114^ or other immune cells, or (B) while CD117 is predominantly expressed on MCs and basophils, it is also expressed in other cell types.

Our data suggests that histamine released from MCs is protective by maintaining the threshold for platelet activation through H_2_R signaling, and thus neutrophil NETosis. Similar to MCs, the role of histamine in breast cancer appears complex, likely due to the four distinct receptors as well as differential roles of histamine in different cell types^115,116^. Mice that cannot synthesize histamine experience high rates of skin and colon cancer, while supplementation with histamine is protective^117^. The reported mechanism was due to reduced myeloid maturation and accumulation of what was likely a myeloid derived suppressor cell population (CD11b^+^;Ly6G^+^)^117^. Intriguingly, large meta-analyses have found that people with allergies (and thus increased histamine) show a decreased overall risk for various forms of cancer, including breast^118,119^. Likewise, people with asthma had reduced cancer risk^120^, although a large retrospective analysis indicated increased risk for several cancers, excluding breast^121^. It is possible that mild allergies and/or asthma results in sufficient histamine signaling via the H_2_R, which would be unaffected by most allergy medication which targets the H_1_R. However, severe allergies/asthma cause inflammation which facilitates cancer growth/progression. However, another study found that use of H_1_R-antagonists was associated with improved survival in patients receiving ICB^122^. The few studies that have assessed use of H_2_R-antagonists such as cimetidine and risk of breast cancer have not demonstrated a link ^123–125^. However, based on our preclinical data, these drugs, which are taken for gastroesophageal reflux disease, are more likely to have an effect on recurrence. Thus, our data provides strong rationale for retrospective analyses or prospective trials evaluating use of H_2_R antagonists and recurrence free survival times. In this regard, it is interesting that the H_2_R-antagonist, ranitidine resulted in decreased B and CD25+ T cells in a single arm prospective trial^126^.

The preclinical reports of histamine and cancer are too many to review properly here, but we would like to highlight a few recent articles to exemplify the paradoxical roles for histamine. First, an H_4_R agonist and antagonist both appeared to reduce tumor growth of the 4T1 mouse model, although the study was underpowered to draw any firm conclusions, other than to highlight the complexity of this axis^127^. It was recently reported that H_1_R activation in tumor associated macrophages results in suppression of cytotoxic CD8^+^ T cells, and that H_1_R-antagonists could counteract this immune-suppression^122^. An informatic screening tool identified an H_2_R antagonist for the prevention of hepatocellular carcinoma^128^. Preclinical models confirmed that this H_2_R antagonist could reduce the incidence of hepatocellular carcinoma in mice administered Diethylnitrosamine^128^.

Collectively, our data support a model whereby preparation of a lipid-rich food under oxidative conditions results in the depletion of MCs and subsequently local histamine concentrations (**Fig. 7**). Loss of histamine or blocking of the H_2_Rs sensitizes platelets, and their activation stimulates NETosis. NETs then impair access of NK and T cells, resulting in a privileged microenvironment, ripe for malignant growth. Given that anti-H_2_R ligands are common, over-the-counter drugs used to treat gastric esophageal reflux disease, these results provide rationale for studies evaluating their use in breast cancer survivors. The involvement of platelets also suggests that drugs limiting platelet activation (non-steroidal anti-inflammatory drugs, heparin etc.) may have use in individuals at high risk of recurrence. Finally, agonists of H_2_R may also have a place in the prevention of recurrence. However, the timing of administration and targeting of these drugs to organs likely to harbor dormant/recurrent lesions would be important considerations for translation.

## METHODS

### Reagents

Histamine (Sigma Aldrich CAT#H7250), cimetidine (Sigma Aldrich CAT#C4522), famotidine (Sigma Aldrich CAT#F6889) and clemastine Fumarate (H_1_R antagonist, 25mg/kg) (Sigma Aldrich CAT#SML0445) were dissolved in DMSO (Sigma Aldrich CAT#D8418) and 0.9% sterile saline.

### Clinical Data

Survival analyses were performed utilizing the Kaplan-Meier Plotter (https://kmplot.com/analysis/index.php?p=service&cancer=breast) and GEPIA 2 (http://gepia2.cancer-pku.cn/#general). The Kaplan-Meier Plotter webtool utilizes data from TCGA, GEO, EGA, Metabric, Impact, and PubMed ^129^. GEPIA 2 uses TCGA/GTEx data sets. Gene signature quantification uses UCSC Xena to allow for greater precision when analyzing coding and non-coding transcripts ^130^.

Archival canine mammary gland sections were retrieved from the College of Veterinary Medicine, selecting 5 grade I, 5 grade II and 5 grade III cases. They were stained with toluidine blue and assessed by a. They were assessed by a veterinary pathologist.

### Flow Cytometry

Flow Cytometry was performed on a Thermo Attune or BD Accuri as follows unless otherwise indicated. FACS Buffer used: DPBS (Corning CAT# MT21031CV) supplemented with 2% FBS (GIBCO Cat# A5670801), 1% penicillin-streptomycin (Corning CAT#30-002-CI). Tissue samples were dissociated in collagenase (Thermo Fisher Scientific CAT#17101015) at 37°C and shaking at 180rpm for 1 hour, and then strained through a 70µM filter (Dot Scientific CAT#15-1070-1). Antibodies were used at 1 to 100 dilution. For compensation ultracomp (Fisher Scientific CAT#501129040) beads were used.

### In Vitro Assays

#### Cancer Cell Line cell culture (D2.0R, 4T1, MET1 etc.)

Murine cancer cells were cultured in DMEM supplemented with 10% FBS (GIBCO Cat# A5670801), 1% penicillin-streptomycin (Corning CAT#30-002-CI), 1% nonessential amino acids (NEAA) (Corning CAT#25-025-CI), 1% sodium pyruvate (NAPY) (Corning#25-000-CI). To passage, cells were washed with DPBS (Corning CAT# MT21031CV) and detached with Trypsin+ EDTA (Corning CAT#MT25053CI). All cells tested negative for mycoplasma.

#### Cellular Proliferation (Cancer Cells)

Cancer cells were resuspended and seeded in a 96-well plate and incubated overnight prior to first treatment. Proliferation was assessed by total DNA using Hoechst 3342 staining, as previously described ^131,132^.

#### Peritoneal Mast Cell (PMC) Isolation

We adapted a protocol from^133^. PMCs were isolated by flushing the peritoneal cavity of mice with 3mL DPBS (Corning CAT# MT21031CV). Cells were centrifuged for 5min at 1,200rpm at 4°C. If blood contamination was evident, cells were washed with Gey’s solution (Thermo Fisher Scientific CAT #J67569.AP) and centrifuged for 5min at 1,200rpm at 4°C. Cells were resuspended in PMC Medium [RPMI-1640 (Corning CAT#17-105-CV) supplemented with 15% FBS(GIBCO Cat# A5670801), 1% penicillin-streptomycin (Corning CAT#30-002-CI), 2mM Glutamax (Fisher Scientific CAT# 35-050-079), 0.07% B-Mercaptoethanol (Thermo Fisher Scientific CAT#21985023), 10mM HEPES (Fisher Scientific CAT# 25060CI), 30ng/mL IL-3 mouse (StemCell Technologies CAT#78042), and 20ng/mL Stem Cell Factor mouse (StemCell Technologies CAT#78064)] and cultured in a 6-well plate. Fresh media is added to the cells every 3^rd^ day. If cells reached confluency the cells are passaged. Enrichment of PMCs was confirmed by FACS analysis [double positive for FcεR1alpha (BD Biosciences CAT#566994) and CD117(BD Biosciences CAT#553356)].

#### Murine Platelet Isolation

Murine blood was collected in a Dipotassium EDTA tube (Fisher Scientific CAT#02-669-33) at room temperature, and plasma isolated and further centrifuged at 18,000 x g for 35min at 22̇C. Pellet was resuspended in DPBS (Corning CAT# MT21031CV) and centrifuged at 300 x g for 10min at 22̇C.

#### Neutrophil Isolation

Neutrophils were cultured in Polymorphonuclear Neutrophil (PMN) Media: RPMI-1640 supplemented with 0.1mg/mL BSA (Sigma Aldrich CAT# A2058-100G) and 1% penicillin streptomycin. Bone marrow was isolated from mouse femur and tibia. Cells were passed through a 70uM strainer (Dot Scientific CAT#15-1070-1), red blood cells were lysed with ACK lysis buffer (Fisher Scientific CAT# A1049201). Ly6G positive cells were isolated utilizing anti-Ly6G UltraPure MicroBeads, mouse (CAT#130-120-337) from Miltenyi Biotec. LS Columns (CAT #130-042-401) and 30uM Pre-Separation Filters (CAT #130-041-407) filters for the magnetic isolation Beads were used in accordance with kit protocol. Isolated cells were cultured in PMN media.

#### NETosis Assay

Isolated neutrophils were counted and resuspended in PMN media and stained with SYTOX Green (Thermo Fisher Scientific CAT# S7020). NETosis was induced with 30nM PMA (Fisher Scientific CAT#52-440-05MG) or 1uM PAR1-6 (Cayman Chemical Company CAT# 27111), in combination with other treatments indicated in the figures. After treatment and staining, cells were incubated for 2 hours prior to flow cytometry analyses.

#### T-cell Isolation

Isolated cells were cultured in T-cell media: RPMI-1640 supplemented with 10% heat-inactivated FBS, 1% penicillin-streptomycin, 1% NEAA and 1% sodium pyruvate. Murine spleen was isolated, cells dispersed and passed through a 70uM strainer, red blood cells were lysed with ACK lysis buffer. The isolation of untouched T-cells utilizes the Pan T Cell Isolation Kit II, mouse (CAT# 130-095-130) from Miltenyi Biotec. LS Columns and 30uM Pre-Separation Filters (CAT #130-041-407) filters for the magnetic isolation Beads were used in accordance to kit protocol. Isolated cells were cultured in T-cell media.

#### T-cell Expansion Assay

T-cells were activated with anti-CD3 (BD Biosciences CAT#553057)/anti-CD28 (BD Biosciences CAT#553294) or for regulatory T cells, anti-CD3, IL-2(Biolegend CAT#575402), TGFß(Biolegend CAT#580702), and B-Mercaptoethanol (Thermo Fisher Scientific CAT#21985023), as previously described ^31,134,135^. T-cells were stained with cFSE (Biolegend CAT#423801) or CellTrace Violet (Thermo Fisher CAT#C34557), and assessed by flow cytometry.

#### Dendritic Cell Isolation

Murine spleen was isolated, cells manually dispersed and passed through a 70uM strainer, red blood cells were lysed with ACK lysis buffer. The isolation of CD11c+ cells utilize the CD11c+ MicroBeads UltraPure, mouse (CAT#130-125-835) from Miltenyi Biotec. LS Columns and 30uM Pre-Separation Filters for the magnetic isolation Beads were used in accordance to kit protocol. Isolated cells were cultured in T-cell media.

#### Bone Marrow Derived Macrophage Isolation

Bone marrow was isolated from mouse femur and tibia. Cells were passed through a 70uM strainer and centrifuged at 400g for 8minutes at 4̇C. Supernatant was aspirated and cells were cultured in DMEM supplemented with 10% FBS, 1% penicillin-streptomycin, 1% NEAA, 1% sodium and 20ng/mL murine mCSF (Peptrotech 315-02-250UG). On day 2 post isolation there was a 50% increase in media and on day 6 post isolation cells were washed with DPBS and provided fresh media. Macrophages were used after least 10d of differentiation.

#### Natural Killer (NK) Cell Isolation

Murine splenic cells were manually dispersed, passed through a 70uM strainer and red blood cells were lysed with ACK lysis buffer. The NK Cell Isolation Kit Miltenyi Biotec (CAT#130-115818), LS Columns and 30uM Pre-Separation Filters for magnetic isolation were used in accordance to kit protocol.

#### Cell Viability

Cells were seeded and treated for 24hours at 37̇C and 5% CO2. Cells were then incubated with 20µL of 0.15mg/ml resazurin for 2-4hours covered in foil at 37̇C and 5% CO2. Fluorescence was recorded by plate reader (excitation 560/emission 590).

#### Mast Cell Degranulation

Protocol adapted from^136^. Briefly matured peritoneal MCs were treated with ligands and incubated for 24hours at 37̇C and 5% CO2. After 24h MCs were washed and seeded. Degranulation was induced with 100uM Compound 48/80 (Cayman Chemical Company CAT# 22173) for 10 minutes at 37̇C ^137–139^. Supernatant was collected and added to plate coated with B-hexosaminidase substrate (Cayman Chemical Company CAT# 28954). Cells were incubated covered in foil for 45minutes at 37̇C. Reaction was quenched with 1M NaOH and optical density was read at 405nM.

### Real-time quantitative polymerase chain reaction (qPCR)

RNA isolation from cell lysate was extracted with GeneJet RNA Purification kit (Thermo Fisher Scientific Cat#K0731). RNA isolation from mouse tissue was isolated from Trizol (Fisher Scientific CAT# 15-596-018), cDNA synthesis with iScript™ Reverse Transcription Supermix (Biorad CAT#1708841BUN), and qPCR performed as previously described^29,140,141^. Cyclophilin or TPB were used as housekeeping genes.

### Mast Cell Histopathology and Analysis

Histopathology analysis was performed at the University of Illinois Urbana-Champaign College of Veterinary Medicine Diagnostic Laboratory. A board-certified veterinary pathologist, blinded to experimental design, analyzed the lung and tumor samples.

### Tumor Histology and Analysis

Lung samples from mice grafted with D2.0R were submitted to the Tumor Engineering and Phenotyping core facility at the Cancer Center at Illinois at Beckman Institute at University of Illinois at Urbana Champaign. Samples were sectioned and stained with Hematoxylin and Eosin. Analysis was done with QuPath.

### nCounter Gene Profiling

RNA isolated from metastatic lung tissue was analyzed on the Nanostring nCounter SPRINT Profiler using the PanCancer IO360 and Myeloid Innate Immunity Panels^142^. Alterations in gene expression were analyzed using nSolver (Ver 4.0) and nCounter Advanced Analysis (Ver2.0)

### Bacon preparation

Our goal was to mimic common home-cooking practice. Bacon was sourced from the University of Illinois Meat Science Laboratory; uncured and cured bacon is from the same animal (different side), controlling for inter-animal differences. Commercially “uncured” can have different meanings. In our case, this is unprocessed and uncured (as in no natural curing). The curing process is carried out by the Meat Science Laboratory under a strict and controlled protocol. Specifically, a cure solution, formulated to deliver 16.75% Sodium Chloride, 3.71% sodium tripolyphosphate, 1.24% Sugar, 0.56% sodium erythorbate, 0.41% spices, and 0.15% Nitrite in a 10% pump. Green weight of the pork bellies was determined and then the bellies were injected using a Wolf-Tec Inc., Kingston, NY injector, at a pressure of 2.2 Bar and 51 rpm using 3 mm needles. Bellies are then weighed again to determine pump uptake of 110% of green weight. Bellies were stored in a metal combo for approximately 12 hours for equilibration. Cured bellies were then placed in an Alkar smokehouse (Lodi, WI) and smoked until they reach an internal temperature of 140 degrees F. After thermal processing, bellies are showered with cold water and placed in a cooler and chilled at 36 degrees F until internal temperature reaches 40 degrees F in accordance with Appendix B of FSIS/USDA regulations^143^. Bacon fat from uncured (ufBF) or cured bacon (cfBF) was generated in the Food Sciences Laboratory, by pan-frying 100-150g batches at 180 +/- 5°C in a cast iron skillet (∼1000 cm^2^ surface area). The lipid temperature was monitored throughout the process. The pan was preheated before the fat sample was placed in the pan to decrease the come-up time for the ufBF/cfBF. Bacon was cooked for 3 min on each side. After frying the bacon fat and fat from frying the uncured pork belly was collected, filtered through three layers of cheese cloth, allowed to cool, and stored frozen under argon. Lard was produced by rendering pig fat in an oven at 350 degrees F and filtered through three layers of cheese cloth and stored frozen under argon.

### Mouse Diets

Diets were modified from the standard AIN-93G diet which is a semi-purified diet designed to meet all the nutritional needs of mice^144^. This allowed for specific manipulation of constituent components, while maintaining the same dietary background between groups. Lipid content was 10% with soybean oil (5%), and ufBF or cfBF (5%), similar to our previous work with thermally abused frying oils^145^. It is important to note, that a diet composed of 10% fat (22% of the calories from fat) is not considered “high-fat”, and allows for the examination of dietary components absent the confounding factor of obesity that could be induced from consuming a high-fat diet^35,146^.

### Animal Studies

All studies involving animals were approved by the Institutional Animal Care and Use Committee (IACUC) at the University of Illinois Urbana-Champaign. All mouse experiments will use female mice as breast cancer primarily affects females.

#### In Vivo Metastatic Colonization: NCD vs HCD

BALB/c mice were fed normal or high cholesterol diets. Normal cholesterol diet (NCD) consisted of 0.025% cholesterol (251ppm) and 0.5% sodium cholate. High cholesterol diet (HCD) contained 1.25% cholesterol (12,748 ppm) and 0.5% sodium cholate. Diets were purchased from TestDiet. Mice were put on diet and grafted with D2.0R cells intravenously, Weekly imaging using In Vivo Live Imaging Systems (IVIS). At day 59 post-graft, mice received weekly treatment of 0.02mg/kg lipopolysaccharide (LPS) by intraperitoneal (IP) injection. Mice were euthanized 94 days post-graft.

#### In Vivo Metastatic Colonization: Uncured vs Cured Bacon Fat

BALB/c mice were placed on diet for 1 month prior to grafting. Mice were grafted intravenously with 10^6^ D2.0R-ff-Luc cells. Weekly imaging using IVIS. Consumption of food was analyzed once per week throughout the study. Mice were euthanized 65 days post-graft.

#### In Vivo Metastatic Colonization Histamine & Anti-histamines

BALB/c mice were grafted intravenously with 10^6^ D2.0R-ff-Luc cells. One week after grafting, mice were treated 5x/week with one of the following: 1) Vehicle (50% v/v saline: DMSO mixture), 2) Histamine (0.5mg/kg), 3) Clemastine Fumarate (H_1_R antagonist, 25mg/kg), and 4) Cimetidine (H_2_R antagonist, 120 mg/kg). Weekly imaging using IVIS. Mice were euthanized 38 days post-graft. At the endpoint, lungs were excised from all mice and then briefly inflated with DPBS. Lungs were then split in half one of which was fixed in 10% formalin for histological staining, and one of which was further divided in half for RNA-Seq analysis and flow cytometric analysis as earlier described.

#### In Vivo Metastatic Colonization Placebo & Famotidine

One week after grafting with D2.0R-ff-Luc cells, mice were treated 5x/week with one of the following: 1) Vehicle (50% v/v saline: DMSO mixture), 2) Famotidine (H_2_R antagonist, 32mg/kg mg/kg). Weekly imaging using IVIS.

#### In Vivo Primary Tumor Growth- Met1 Placebo, Cimetidine, Famotidine

FVB/NJ mice were grafted with MET1 iRFP cells in the mammary fat pad. Tumors were measured via caliper measurements every other day. Mice were treated 5X per week with Saline, Cimetidine or Famotidine.

#### In Vivo Metastatic Colonization Mast Cell Depletion

BALB/c mice were grafted intravenously with 10^6^ D2.0R-ff-Luc cells. Baseline luminescence images were captured 72 hours after graft using IVIS. Mice were treated with anti-IgG or anti-FcER1alpha. Weekly imaging proceeded until tumor burden more than doubled from baseline.

#### In Vivo Metastatic Colonization- 4T1 timecourse

BALB/c mice were grafted intravenously with 4T1 cells. Mice were euthanized on days 0, 1, 5 and 10. Tissues were taken and analyzed with qPCR for gene expression.

#### In Vivo Metastatic Colonization Athymic Mice Placebo & Cimetidine

Athymic nude (nu/nu) mice were grafted intravenously with 10^6^ D2.0R-ff-Luc cells. Baseline luminescence images were captured 72 hours after graft using IVIS. Weekly imaging proceeded until tumor burden more than doubled from baseline. Day 3 post graft mice were treated 5 days/week with Cimetidine 120mg/kg or Saline control.

#### In Vivo Anti-Histamines on Basal NETosis

BALB/c, mice were dosed with IP 120mg/kg Cimetidine, 32mg/kg Famotidine, or Saline control daily, 5 days/week, for 49 days. At endpoint blood was isolated from cardiac puncture, and platelets and neutrophils were isolated as previously described. The isolated neutrophils and platelets were co-cultured to examine basal rates NETosis induced by 30nM PMA or 1uM PAR1-6 as described. Additionally, lungs were dissociated and used for flow cytometric analysis.

#### In Vivo Metastatic Colonization Anti-histamine, Heparin, DNAse I

BALB/c, mice were grafted intravenously D2.0R-ff-Luc cells. Three days post grafting mice were treated daily, 5 days/week, with the following: 1) Vehicle (Saline+10% DMSO), 2) Cimetidine (120mg/kg), 3) Heparin, 4) Cimetidine+Heparin, 5) DNAse I, 6) Cimetidine + DNAse I. Weekly imaging proceeded until tumor burden more than doubled from baseline.

#### In Vivo Met1 anti-Ly6G

FVB/NJ mice were grafted orthotopically with MET1 iRFP cells. Tumors were measured via caliper measurements initially every other day, and then every day once tumors had established. Mice were dosed daily with 1) Vehicle + αIgG, 2) Vehicle + αLy6G, 3) Cimetidine + αIgG, 4) Cimetidine + αLy6G.

### Statistics

Data expressed as mean ± standard error mean unless otherwise noted. Statistical analyses were performed using GraphPad Prism or by hand. Data was assessed for normality and either raw or ln-transformed was used for statistical analyses, either parametric or non-parametric as appropriate. Specific statistical tests are indicated in the figure legends. Initial informatics analyses of RNAseq were performed by HPCBio, University of Illinois. GO and GSEA analyses were performed in part with Shiny GO^147^.

## Supplementary Figure Legends

**Supplementary Figure 1: *Dietary cholesterol does not influence reemergence from dormancy, while its oxidized metabolite, 27-hydroxycholesterol (27HC) does.* (A)** Mice were placed on a control, normal cholesterol diet (NCD) or high cholesterol diet (HCD) for 8 weeks prior to graft with D2.0R cells to model dormant lesions. LPS was administered to spark reemergence. A doubling of metastatic burden (luciferase signal) was considered breaking of dormancy. No significant differences in recurrence were observed between groups (P value from Logrank test, N=40). **(B)** D2.0R cells were grafted into mice to model dormant lesions. 20 days after graft, mice were treated with either placebo or 20mg/kg 27HC, daily through 55 days post-graft (treatment window indicated by dashed lines), and then followed through time. A doubling of metastatic burden (luciferase signal) was considered breaking of dormancy (P value from Logrank test, N=25). **(C)** Py230 cells were grafted orthotopically. These grafts have a very long latency prior to detection of a palpable tumor, making them a model of dormancy. Treatment with placebo or 27HC commenced 3d after graft. Time to detection of a palpable tumor is shown (P value from Logrank test, N=12). **(D)** Met1 cells were grafted and tumors allowed to establish to ∼200mm^3^, at which point they were surgically removed and mice treated with three rounds of doxorubicin (dosed every 2d). Mice were then treated with placebo or 27HC for 21 days. Lungs were then imaged *ex vivo*; representative images to the left of quantified data. (unpaired t test, N=16).

**Supplementary Figure 2: *Fatty acid characterization of fat from cured, fried bacon (cfBF) or rendered lard.* (A)** Bacon content was characterized for water and lipid content after pan-frying. The left two pie charts are based on dried samples, whereas the right pie chart is based on wet-bacon (undried). **(B)** Total ion chromatogram of 36 mixed fatty acid methyl esters (FAMEs) analyzed by GC-MS on an HP-88 column (60 m × 0.25 mm, 0.20 µm). The oven temperature program was as follows: initial temperature of 60 °C for 1 min, ramped at 10 °C/min to 145 °C, then at 1 °C/min to 190 °C, and finally at 5 °C/min to 220 °C. Mass spectrometry was operated in scan mode over an m/z range of 50–500 amu with an EI voltage of 70 eV. Compound identification is indicated in **Supplementary Table 1A**. **(C)** GC-MS chromatograms of fatty acid methyl esters of lard and cfBF. Compound identification and quantification shown in **Supplementary Table 1B**. **(D)** PCA plots of cfBF vs lard. Individual samples were plotted based on the first two principle components (PC1 and PC2), which accounts for 60.7% and 33.2% of the total variance, respectively. The clustering suggests a clear separation from cfBF and rendered lard.

**Supplementary Figure 3: *Mice consuming diets enriched with lard or cured fried bacon fat (cfBF) do not have altered weight, but their tumors have decreased T cell infiltration.* (A)** Mean body weights through time of 4T1 tumor-bearing mice consuming control (modified AIN-93G to contain 10% fat from soybean oil), lard (5% lard, 5% soybean oil) or cfBF (5% cfBF, 5% soybean oil) diets. Data is presented as mean mass ± SEM (N=40). **(B)** Mean food consumed per mouse per day. The data from A & B correspond to the data in **Fig. 1A** of the main text. **(C)** Flow cytometric analysis of primary tumors found decreased T cell infiltration in mice fed cfBF (N=39, 1-Way ANOVA followed by multiple comparison test of geometric means with Šidák’s correction). **(D)** Left panel: Mean body weights through time of mice bearing 4T1 metastatic lesions, consuming control, lard or cfBF diets. Right Panel: Corresponding food consumption data. These data correspond to **Fig. 1B** of the main text. **(E)** Left panel: Mean body weights through time of mice bearing Met1 metastatic lesions, consuming control, lard or cfBF diets. Right Panel: Corresponding food consumption data. These data correspond to **Fig. 1C** of the main text. **(F)** Met1 metastatic lesions from mice consuming cfBF had decreased T cell infiltrate (N=34; 1-Way ANOVA followed by multiple comparison test of geometric means with Šidák’s correction). Different letters denote statistically significant differences.

**Supplementary Figure 4: *Mouse body weights and food consumption are not different when consuming different experimental diets***. Left panel: Mean body weights through time of mice grafted with a sub-optimal number of 4T1 cells, consuming control, lard or cfBF diets. Right Panel: Corresponding food consumption data. (N=25) These data correspond to **Fig. 1D** of the main text.

**Supplementary Figure 5: *Mice consuming a diet enriched in uncured fried bacon fat (ufBF) or cfBF did not show altered food consumption, but significantly increased metastatic burden was observed in cfBF fed mice.* (A)** Mean consumption of food per week comparing mice consuming ufBF or cfBF. **(B)** Fold change in metastatic burden of D2.0R bearing mice consuming ufBF or cfBF. Left: total metastatic burden. Middle: bone only. Right: Lung and liver only. These data correspond to **Fig. 1E** in the main text. Asterisks indicate statistical significance (Mann-Whitney U test or students T test, P<0.05).

**Supplementary Figure 6: *Flow cytometric analyses of lungs from naïve mice on different diets.*** Dendritic cells (CD11C+) and NK cells (NKp46+) were assessed by flow cytometry. These data correspond to **Fig. 1F** of the main text.

**Supplementary Figure 7: *Analysis of Nanostring results predicts decreases in dendritic cells.*** Transcriptomics was performed using Nanostring. Informatic analyses predicted the relative abundance of different immune cell types. These data correspond to **Figs. 1E** and **2A** in the main text.

**Supplementary Figure 8: *Expression of genes associated with MCs are a good prognostic across several tumor types.* (A)** *FCεR1a*, a gene selectively expressed on MCs, is decreased in human breast tumors with higher stage compared to lower. Data from the TCGA Invasive Breast Cancer Firehose Legacy archive, binned into tumors of stage 1, or 2&3 (unpaired t test, N=471). **(B)** Elevated expression of a *FCεR1a* in human breast tumors is associated with increased recurrence free survival (RFS). All breast cancer subtypes considered here. Data accessed from the Kaplan-Meier Plotter webtool based on data from GEO, EGA, and TCGA (P value indicated from Logrank test, N=4890). The Kaplan-Meier Plotter webtool uses aggregated data from GEO, EGA, and TCGA^148^. **(C)** Elevated expression of the ‘MC-gene signature’ in tumors from patients treated with immune checkpoint blockers (ICB) is associated with an increased progression free survival time. Tumor types included for this analysis were bladder, esophageal adenocarcinoma, glioblastoma, hepatocellular carcinoma, head and neck squamous cell carcinoma, melanoma, non-small cell lung cancer and urothelial cancer. (P value from Logrank test, N=463). **(D-E)** Elevated expression of the ‘MC-gene signature’ in lung cancer tumors is associated with longer recurrence free- and overall-survival (RFS and OS respectively, P value from Logrank test, RFS N=1218, OS N=2154). **(F-G)** Elevated expression of the ‘MC-gene signature’ in acute myeloid leukemia (AML) is associated with longer event free- and overall-survival (P value from Logrank test, EFS N=525, OS N=1603). **(H-I)** Elevated expression of the ‘MC-gene signature’ in multiple myeloma samples is associated with RFS and OS (P value from Logrank test, RFS N=797, OS N=1415).

**Supplementary Figure 9: *Loss of mast cells (MCs) results in recurrence from mammary cancer dormancy***.

**(A)** Overview of experimental design, corresponding to **Fig. 2H**. **(B)** Administration of an antibody against FCεR1 (αFCεR1) successfully immune-depletes peritoneal MCs. **(C)** αFCεR1 depletes MCs within D2.0R bearing lungs. Samples taken at the end of the study (Day 84, N=35). **(D-J)** Flow cytometric or qPCR analysis of immune cell populations within D2.0R bearing lungs. (D, G-J are flow cytometry, N=32-35; E-F are qPCR, N=33-35 (unpaired T test).

**Supplementary Figure 10: *HDC expression in human tumors is associated with improved survival***. Histidine decarboxylase (*HDC)* expression in **(A)** lung, **(B)** acute myeloid leukemia, or **(C)** myeloma is associated with improved overall survival. **(D)** Elevated *HDC* expression in tumors is associated with improved response to ICB. Data accessed from the Kaplan-Meier Plotter webtool based on data from GEO, EGA, and TCGA (P values indicated from Logrank test, A: N=2154, B: N=1603, C: N=1416, D: N=463).

**Supplementary Figure 11: *Expression of enzymes involved in histamine catabolism are associated with poor survival in different tumor types.*** Tumor types are by row, denoted on the left, and gene is by column, denoted on the top. Data accessed from the Kaplan-Meier Plotter webtool based on data from GEO, EGA, and TCGA (P values indicated from Logrank test, with the exception of AOC2-Breast and DAO-AML which used the Gehan Breslow Wilcoxon test). Sample numbers (N) are indicated on each graph.

**Supplementary Figure 12: *Expression of H_2_R in 4T1 metastatic lesions decreases through time.*** 4T1 cells were grafted intravenously. Mice were euthanized at indicated timepoints, and lungs assessed for mRNA content by qPCR. Data was fit to a one-phase decay regression model. Pearson’s one-way correlation was assessed with r and P values indicated. **(A)** H_1_R (*Hrh1*, N=17). **(B)** H_2_R (*Hrh2*, N=17). **(C)** H_3_R (*Hrh3*, N=17). **(D)** H_4_R (*Hrh4*, N=17).

**Supplementary Figure 13: *Antagonists to H_2_R do not have significant effects on proliferation of D2.0R or 4T1 cells***. **(A)** D2.0R cells were plated on day 0. On day 1 they were treated with vehicle (DMSO) or ligands at the indicated doses. On Day 4 the relative cell abundance was assessed by total DNA content (as described in ^131^). **(B)** 4T1 cells were treated as in (A) and assessed for total DNA content. Values are relative to vehicle (D2.0R N=58; 4T1 N=65).

**Supplementary Figure 14: *Single Cell RNA-sequencing of human and murine samples indicates elevated expression of H_2_R in myeloid immune cells.* (A-C)** Assessment of *HRH2* (H_2_R mRNA) expression by scRNA-seq circulating immune cells across three different datasets (data obtained from Human Protein Atlas; proteinatlas.org^149^). **(D-G)** scRNA-seq of breast/mammary tumors indicates *HRH2* expression across several different immune cells. Data was obtained from the indicated databases and accessed through TISCH2 ^150^. GSE176078: 26 tumors, 89,471 cells. GSE161529: 52 tumors, 332,168 cells. EMTAB8107: 14 tumors, 33,043 cells. GSE136206: 27,352 cells. (H) *Hrh2* expression in different isolated murine cell types and platelets, as determined by qPCR. Expression was normalized to cyclophilin.

**Supplementary Figure 15: *H_2_R antagonists do not have significant effects on NK cell viability or markers of activity.* (A)** Splenic NK cells were cultured in the presence of the indicated ligands at increasing concentrations (histamine: 1 or 2µM, cimetidine: 12 or 24µM, famotidine: 1 or 2µM, clemastine: 240nM) for 24h. Cell viability was assessed by reduction of resazurin to resorufin (as described^134,135^). **(B)** Markers of NK cell activation were assessed by qPCR after treatment with the indicated ligands for 24h. Statistical differences, if any, are denoted by different letters (N=21-23 per gene analysis, 1-Way ANOVA followed by multiple comparison test of geometric means with Šidák’s correction).

**Supplementary Figure 16: *H_2_R antagonists do not have significant effects on T cell viability, markers of activity or ability to expand.* (A)** Splenic T cells were cultured in the presence of indicated ligands at increasing concentrations (histamine: 1 or 2µM, cimetidine: 12 or 24µM, famotidine: 1 or 2µM, clemastine: 240nM) for 24h. Cell viability was assessed by reduction of resazurin to resorufin. **(B)** T cell expansion. CFSE-stained T cells were incubated with indicated ligands and activated with antibodies against CD3 and CD28. T cell proliferation was assessed with flow cytometry for dilution of CFSE signal. From left to right: pan T cells, CD4+ only, CD4+;FoxP3+ (Treg) only, and CD8+ only. **(C)** CFSE-stained T cells were cultured in media to support differentiation into Tregs (includes IL2, TGFβ and β-mercaptoethanol), and activated with antibodies against CD3 and CD8. T cell proliferation was assessed with flow cytometry for dilution of CFSE signal. From left to right: pan T cells, CD4+ only, CD4+;FoxP3+ (Treg) only, and CD8+ only.

**Supplementary Figure 17: *H_2_R antagonists do not have significant effects on macrophage markers of activity or ability to support T cell expansion* (A)** Bone marrow derived macrophages were cultured in the presence of indicated ligands at increasing concentrations (histamine: 1 or 2µM, cimetidine: 12 or 24µM, famotidine: 1 or 2µM, clemastine: 240nM) for 24h. Cell viability was assessed by reduction of resazurin to resorufin. **(B)** Markers of macrophage activation were assessed by qPCR after treatment of BMDMs with the indicated ligands for 24h. Statistical differences, if any, are denoted by different letters (N=16 per gene analysis, 1-Way ANOVA followed by multiple comparison test of geometric means with Šidák’s correction). **(C)** Bone marrow derived macrophages were pre-treated with indicated ligands, washed and then co-cultured with CFSE-stained and activated T cells. T cell proliferation was assessed with flow cytometry for dilution of CFSE signal. From left to right: pan T cells, CD4+ only, and CD8+ only (no statistically significant differences, N=25).

**Supplementary Figure 18: H_2_R antagonists do not have significant effects on dendritic cell (DC) viability, markers of activity, or ability to support T cell expansion. (A)** DCs were cultured in the presence of indicated ligands at increasing concentrations (histamine: 1 or 2µM, cimetidine: 12 or 24µM, famotidine: 1 or 2µM, clemastine: 240nM) for 24h. Cell viability was assessed by reduction of resazurin to resorufin. **(B)** Markers of DC activation were assessed by qPCR after treatment with the indicated ligands for 24h. **(C)** DCs were pre-treated with indicated ligands, washed and then co-cultured with CFSE-stained and activated T cells. T cell proliferation was assessed with flow cytometry for dilution of CFSE signal. From left to right: pan T cells, CD4+ only, CD4+;FoxP3+ (Treg) only, and CD8+ only. **(D)** Resulting expanded T cells were further assessed for markers including (from left to right): IL4, IL17, IFNγ in CD4+ cells, and IFNγ in CD8+ cells.

**Supplementary Figure 19: H_2_R antagonists do not have significant effects on mast cell (MC) viability, markers of activity, degranulation, or ability to present antigen and activate T cells. (A)** MCs were cultured in the presence of indicated ligands at increasing concentrations (histamine: 1 or 2µM, cimetidine: 12 or 24µM, famotidine: 1 or 2µM, clemastine: 240nM) for 24h. Cell viability was assessed by reduction of resazurin to resorufin. **(B)** Markers of MC activation were assessed by qPCR after treatment with the indicated ligands for 24h. **(C)** MCs were treated with indicated ligands in the absence or presence of the degranulation inducer, C48/80. Degranulation was assessed by enzymatic activity of β- hexosaminidase. **(D)** CFSE-stained OTII T cells were cultured in the presence of MCs previously pulsed with OVA. Subsequent T cell proliferation was assessed with flow cytometry for dilution of CFSE signal. OTII T cell expansion after OVA antigen presentation is predominantly CD4+. **(E)** CFSE-stained OTI T cells were cultured in the presence of MCs previously pulsed with OVA. Subsequent T cell proliferation was assessed with flow cytometry for dilution of CFSE signal. OTI T cell expansion after OVA antigen presentation is predominantly CD8+. **(F)** DCs and MCs were co-cultured. Pre-treated DCs and MCs were then co-cultured with CFSE-stained and activated T cells. T cell proliferation was assessed with flow cytometry for dilution of CFSE signal. From left to right: pan T cells, CD4+ only, and CD8+ only.

**Supplementary Figure 20: *H_2_R antagonists do not have significant effects on neutrophil viability or markers of activity.* (A)** Neutrophils were cultured in the presence of indicated ligands at increasing concentrations (histamine: 1 or 2µM, cimetidine: 12 or 24µM, famotidine: 1 or 2µM, clemastine: 240nM) for 24h. Cell viability was assessed by reduction of resazurin to resorufin. **(B)** Markers of neutrophil activity were assessed by qPCR after treatment with the indicated ligands for 24h.

**Supplementary Figure 21: *Increased presence of activated platelets and NETs after chronic treatment with H_2_R antagonists.* (A)** Flow cytometric analysis for CD61+ or **(B)** CD41a+ events, both markers of platelets. Different letters denote statistical significance (1-Way ANOVA followed by multiple comparison test with Šidák’s correction, N=43).

**Supplementary Figure 22: *Cimetidine or famotidine increase NETs as determined by flow cytometry for MPO+ events***. Flow cytometric analysis of MPO+ events after a co-culture of platelets and neutrophils from mice chronically treated with the indicated ligands. Platelets and neutrophils were derived from either **(A)** the bone marrow, or **(B)** the spleen. Different letters denote statistical significance (1-Way ANOVA followed by multiple comparison test with Šidák’s correction).

## Declarations

All procedures involving mice were approved by the Institutional Animal Care and Use Committee (IACUC) at the University of Illinois Urbana Champaign. Humane euthanasia was consistent with American Veterinary Medical Association (AVMA) guidelines. All authors have provided consent for the publication of this work. Data, reagents and/or protocol information will be provided upon reasonable request. RNA-seq data will be uploaded to GEO upon acceptance of this manuscript. There are no competing interests to declare.

## Funding

American Institute for Cancer Research (713063 and 579732)

Department of Defense Era of Hope Scholar Award BC200206/W81XWH-20-BCRP-EOHS (ERN)

Department of Defense BC241117/HT9425-25-1-0285 (ERN)

National Institutes of Health grant R01 CA288207 (ERN)

National Institutes of Health grant R01 CA234025 (ERN)

National Institutes of Health grant T32 GM136629 (HEVG)

National Institutes of Health grant T32 ES007326 (ATN)

National Institutes of Health grant T32 EB019944 (CPS)

The Endocrine Society Research Experiences for Graduate and Medical Students Award (SVB, DP)

The Cancer Center at Illinois Graduate Cancer Scholarship Program Award (SVB)

Postdoctoral Fellows Program at the Beckman Institute for Advanced Science and Technology (NK)

Vision 20/20 USDA Hatch Funds (ERN, WGH)

Keith W. and Sara M. Kelley Endowed Professorship in Immunophysiology (ERN)

Cancer Center at Illinois and University of Illinois (ERN)

-The funders did not have a role in study design, interpretation of results or writing of this manuscript.

## Acknowledgments

We would like to thank the patients whose tumors populated data in the TCGA, METABRIC and Human Protein Atlas initiatives. We would also like to thank the Cancer Research Advocacy Group at the University of Illinois and our breast cancer advocate team: Sarah Adams, Renaé Strawbridge, Jamie Holloway, Lea Ann Carson, Susan Stewart and Catherine Applegate. The Tumor Engineering and Phenotyping Core at the Cancer Center at Illinois provided mycoplasma testing, some histology and Nanostring analysis. The Roy J. Carver Biotechnology Center performed RNA-sequencing (DNA Services unit) and initial processing of RNA sequencing data (HPCBio unit). We extend special thanks to NR IMPACT for their early feedback on this work.

## References Cited

1 Rahib, L., Wehner, M. R., Matrisian, L. M. & Nead, K. T. Estimated Projection of US Cancer Incidence and Death to 2040. JAMA Netw Open 4, e214708 (2021). 10.1001/jamanetworkopen.2021.4708

2 Chaffer, C. L., San Juan, B. P., Lim, E. & Weinberg, R. A. EMT, cell plasticity and metastasis. Cancer Metastasis Rev 35, 645–654 (2016). 10.1007/s10555-016-9648-7

3 Welch, D. R. & Hurst, D. R. Defining the Hallmarks of Metastasis. Cancer Res 79, 3011–3027 (2019). 10.1158/0008-5472.CAN-19-0458

4 Morrissey, C., Vessella, R. L., Lange, P. H. & Lam, H. M. The biology and clinical implications of prostate cancer dormancy and metastasis. J Mol Med (Berl) 94, 259–265 (2016). 10.1007/s00109-015-1353-4

5 Ring, A., Spataro, M., Wicki, A. & Aceto, N. Clinical and Biological Aspects of Disseminated Tumor Cells and Dormancy in Breast Cancer. Front Cell Dev Biol 10, 929893 (2022). 10.3389/fcell.2022.929893

6 Braun, S., et al. A pooled analysis of bone marrow micrometastasis in breast cancer. The New England journal of medicine 353, 793–802 (2005). 10.1056/NEJMoa050434

7 Klein, C. A. Parallel progression of primary tumours and metastases. Nat Rev Cancer 9, 302–312 (2009). 10.1038/nrc2627

8 Heidrich, I., Deitert, B., Werner, S. & Pantel, K. Liquid biopsy for monitoring of tumor dormancy and early detection of disease recurrence in solid tumors. Cancer Metastasis Rev 42, 161–182 (2023). 10.1007/s10555-022-10075-x

9 Braun, S., et al. Cytokeratin-positive cells in the bone marrow and survival of patients with stage I, II, or III breast cancer. The New England journal of medicine 342, 525–533 (2000). 10.1056/NEJM200002243420801

10 Hartkopf, A. D., et al. Disseminated tumour cells from the bone marrow of early breast cancer patients: Results from an international pooled analysis. Eur J Cancer 154, 128–137 (2021). 10.1016/j.ejca.2021.06.028

11 Phan, T. G. & Croucher, P. I. The dormant cancer cell life cycle. Nat Rev Cancer 20, 398–411 (2020). 10.1038/s41568-020-0263-0

12 Bouchard, G., et al. Pre-irradiation of mouse mammary gland stimulates cancer cell migration and development of lung metastases. Br J Cancer 109, 1829–1838 (2013). 10.1038/bjc.2013.502

13 El Rayes, T., et al. Lung inflammation promotes metastasis through neutrophil protease-mediated degradation of Tsp-1. Proc Natl Acad Sci U S A 112, 16000–16005 (2015). 10.1073/pnas.1507294112

14 Cox, T. R., et al. LOX-mediated collagen crosslinking is responsible for fibrosis-enhanced metastasis. Cancer Res 73, 1721–1732 (2013). 10.1158/0008-5472.CAN-12-2233

15 Albrengues, J., et al. Neutrophil extracellular traps produced during inflammation awaken dormant cancer cells in mice. Science 361 (2018). 10.1126/science.aao4227

16 He, X. Y., et al. Chronic stress increases metastasis via neutrophil-mediated changes to the microenvironment. Cancer Cell 42, 474–486 e412 (2024). 10.1016/j.ccell.2024.01.013

17 Park, J., et al. Cancer cells induce metastasis-supporting neutrophil extracellular DNA traps. Sci Transl Med 8, 361ra138 (2016). 10.1126/scitranslmed.aag1711

18 Greten, F. R. & Grivennikov, S. I. Inflammation and Cancer: Triggers, Mechanisms, and Consequences. Immunity 51, 27–41 (2019). 10.1016/j.immuni.2019.06.025

19 Dolberg, D. S., Hollingsworth, R., Hertle, M. & Bissell, M. J. Wounding and its role in RSV-mediated tumor formation. Science 230, 676–678 (1985). 10.1126/science.2996144

20 De Cock, J. M., et al. Inflammation Triggers Zeb1-Dependent Escape from Tumor Latency. Cancer Res 76, 6778–6784 (2016). 10.1158/0008-5472.CAN-16-0608

21 Wang, Y., Bendre, S. V., Krauklis, S. A., Steelman, A. J. & Nelson, E. R. Role of Protein Regulators of Cholesterol Homeostasis in Immune Modulation and Cancer Pathophysiology. Endocrinology 166 (2025). 10.1210/endocr/bqaf031

22 Baek, A. E. & Nelson, E. R. The Contribution of Cholesterol and Its Metabolites to the Pathophysiology of Breast Cancer. Horm Cancer 7, 219–228 (2016). 10.1007/s12672-016-0262-5

23 Ahern, T. P., et al. Statin prescriptions and breast cancer recurrence risk: a Danish nationwide prospective cohort study. Journal of the National Cancer Institute 103, 1461–1468 (2011). 10.1093/jnci/djr291

24 Kwan, M. L., Habel, L. A., Flick, E. D., Quesenberry, C. P. & Caan, B. Post-diagnosis statin use and breast cancer recurrence in a prospective cohort study of early stage breast cancer survivors. Breast cancer research and treatment 109, 573–579 (2008). 10.1007/s10549-007-9683-8

25 Nielsen, S. F., Nordestgaard, B. G. & Bojesen, S. E. Statin use and reduced cancer-related mortality. The New England journal of medicine 367, 1792–1802 (2012). 10.1056/NEJMoa1201735

26 Nowakowska, M. K., et al. Association of statin use with clinical outcomes in patients with triple-negative breast cancer. Cancer 127, 4142–4150 (2021). 10.1002/cncr.33797

27 Mei, W., et al. A commonly inherited human PCSK9 germline variant drives breast cancer metastasis via LRP1 receptor. Cell 188, 371–389 e328 (2025). 10.1016/j.cell.2024.11.009

28 Baek, A. E., et al. The cholesterol metabolite 27 hydroxycholesterol facilitates breast cancer metastasis through its actions on immune cells. Nature Communications 8, 864 (2017). 10.1038/s41467-017-00910-z

29 He, S., et al. Host CYP27A1 expression is essential for ovarian cancer progression. Endocr Relat Cancer 26, 659–675 (2019). 10.1530/ERC-18-0572

30 Ma, L., et al. The Liver X Receptor Is Selectively Modulated to Differentially Alter Female Mammary Metastasis-associated Myeloid Cells. Endocrinology 163 (2022). 10.1210/endocr/bqac072

31 Ma, L., et al. 27-Hydroxycholesterol acts on myeloid immune cells to induce T cell dysfunction, promoting breast cancer progression. Cancer letters 493, 266–283 (2020). 10.1016/j.canlet.2020.08.020

32 Krawczynska, N., et al. Neutrophils exposed to a cholesterol metabolite secrete extracellular vesicles that promote epithelial-mesenchymal transition and stemness in breast cancer cells. Cancer letters 636, 218105 (2026). 10.1016/j.canlet.2025.218105

33 Das Gupta, A., et al. 27-Hydroxycholesterol Enhances Secretion of Extracellular Vesicles by ROS-Induced Dysregulation of Lysosomes. Endocrinology (2024). 10.1210/endocr/bqae127

34 McDonnell, D. P., et al. Obesity, cholesterol metabolism, and breast cancer pathogenesis. Cancer Res 74, 4976–4982 (2014). 10.1158/0008-5472.CAN-14-1756

35 Nelson, E. R., Chang, C. Y. & McDonnell, D. P. Cholesterol and breast cancer pathophysiology. Trends in endocrinology and metabolism: TEM 25, 649–655 (2014). 10.1016/j.tem.2014.10.001

36 Li, X., et al. Modifiable Risk Factors for Breast Cancer: Insights From Systematic Reviews. Public Health Nurs 42, 1060–1071 (2025). 10.1111/phn.13504

37 Uhomoibhi, T. O., et al. High-Fat Diet as a Risk Factor for Breast Cancer: A Meta-Analysis. Cureus 14, e32309 (2022). 10.7759/cureus.32309

38 Mahjourian, M., Anjom-Shoae, J., Mohammadi, M. A., Feinle-Bisset, C. & Sadeghi, O. Associations of dietary fat types (MUFA, PUFA, SFA) and sources (animal, plant) with colorectal cancer risk: A comprehensive systematic review and dose-response meta-analysis of prospective cohort studies. Cancer Epidemiol 95, 102768 (2025). 10.1016/j.canep.2025.102768

39 Martin-Sierra, C., Laranjeira, P., Domingues, M. R. & Paiva, A. Lipoxidation and cancer immunity. Redox Biol 23, 101103 (2019). 10.1016/j.redox.2019.101103

40 Rupert, J., Cao, P. H. A., Frigo, D. E. & Kolonin, M. G. Lipids grease the chain of cancer progression. Trends Cancer (2026). 10.1016/j.trecan.2025.11.012

41 Ma, C., et al. Lipotoxicity, lipid peroxidation and ferroptosis: a dilemma in cancer therapy. Cell Biol Toxicol 41, 75 (2025). 10.1007/s10565-025-10025-7

42 Hicks, K. C., Tyurina, Y. Y., Kagan, V. E. & Gabrilovich, D. I. Myeloid Cell-Derived Oxidized Lipids and Regulation of the Tumor Microenvironment. Cancer Res 82, 187–194 (2022). 10.1158/0008-5472.CAN-21-3054

43 Castillo, J. G., et al. Selective disruption of lipid peroxide homeostasis in intratumoral regulatory T cells by targeting FSP1 enhances cancer immunity. Sci Adv 12, eaea3703 (2026). 10.1126/sciadv.aea3703

44 Nelson, E. R., et al. 27-Hydroxycholesterol links hypercholesterolemia and breast cancer pathophysiology. Science 342, 1094–1098 (2013). 10.1126/science.1241908

45 Baek, A. E., et al. The cholesterol metabolite 27-hydroxycholesterol increases the secretion of extracellular vesicles which promote breast cancer progression. Endocrinology (2021). 10.1210/endocr/bqab095

46 Lappano, R., et al. The cholesterol metabolite 25-hydroxycholesterol activates estrogen receptor alpha-mediated signaling in cancer cells and in cardiomyocytes. PLoS One 6, e16631 (2011). 10.1371/journal.pone.0016631

47 BT Bacon Today.

48 Echarte, M., Ansorena, D. & Astiasaran, I. Fatty acid modifications and cholesterol oxidation in pork loin during frying at different temperatures. J Food Prot 64, 1062–1066 (2001).

49 Echarte, M., Zulet, M. A. & Astiasaran, I. Oxidation process affecting fatty acids and cholesterol in fried and roasted salmon. J Agric Food Chem 49, 5662–5667 (2001). 10.1021/jf010199e

50 Ferioli, F., Dutta, P. C. & Caboni, M. F. Cholesterol and lipid oxidation in raw and pan-fried minced beef stored under aerobic packaging. J Sci Food Agric 90, 1050–1055 (2010). 10.1002/jsfa.3918

51 Ferreira, F. S., et al. Impact of Air Frying on Cholesterol and Fatty Acids Oxidation in Sardines: Protective Effects of Aromatic Herbs. J Food Sci 82, 2823–2831 (2017). 10.1111/1750-3841.13967

52 Sabolova, M., et al. Formation of oxysterols during thermal processing and frozen storage of cooked minced meat. J Sci Food Agric 97, 5092–5099 (2017). 10.1002/jsfa.8386

53 Hu, X.-f., et al. Effect of frying on the lipid oxidation and volatile substances in grass carp (Ctenopharyngodon idellus) fillet. Journal of Food Processing and Preservation 46, e16342 (2022). 10.1111/jfpp.16342

54 Al-Saghir, S., et al. Effects of Different Cooking Procedures on Lipid Quality and Cholesterol Oxidation of Farmed Salmon Fish (Salmo salar). Journal of Agricultural and Food Chemistry 52, 5290–5296 (2004). 10.1021/jf0495946

55 Bejaoui, S., Chetoui, I., Ghribi, F., Soudani, N. & Cafsi, M. E. Different frying processes stimulate lipid peroxidation and promote changes in the composition of cholesterol, free fatty acids and triglycerides in the commercial clam’s tissues Venerupis decussata. International Journal of Food Engineering 18, 87–103 (2022). doi:10.1515/ijfe-2021-0224

56 Mazzoni, A., Siraganian, R. P., Leifer, C. A. & Segal, D. M. Dendritic cell modulation by mast cells controls the Th1/Th2 balance in responding T cells. J Immunol 177, 3577–3581 (2006). 10.4049/jimmunol.177.6.3577

57 Dudeck, A., Suender, C. A., Kostka, S. L., von Stebut, E. & Maurer, M. Mast cells promote Th1 and Th17 responses by modulating dendritic cell maturation and function. Eur J Immunol 41, 1883–1893 (2011). 10.1002/eji.201040994

58 Kitawaki, T., et al. IgE-activated mast cells in combination with pro-inflammatory factors induce Th2-promoting dendritic cells. Int Immunol 18, 1789–1799 (2006). 10.1093/intimm/dxl113

59 Reuter, S., et al. Mast cells induce migration of dendritic cells in a murine model of acute allergic airway disease. Int Arch Allergy Immunol 151, 214–222 (2010). 10.1159/000242359

60 Shelburne, C. P., et al. Mast cells augment adaptive immunity by orchestrating dendritic cell trafficking through infected tissues. Cell Host Microbe 6, 331–342 (2009). 10.1016/j.chom.2009.09.004

61 Dawicki, W., Jawdat, D. W., Xu, N. & Marshall, J. S. Mast cells, histamine, and IL-6 regulate the selective influx of dendritic cell subsets into an inflamed lymph node. J Immunol 184, 2116–2123 (2010). 10.4049/jimmunol.0803894

62 de Vries, V. C., et al. Mast cells condition dendritic cells to mediate allograft tolerance. Immunity 35, 550–561 (2011). 10.1016/j.immuni.2011.09.012

63 Carroll-Portillo, A., et al. Mast cells and dendritic cells form synapses that facilitate antigen transfer for T cell activation. J Cell Biol 210, 851–864 (2015). 10.1083/jcb.201412074

64 Mazzoni, A., Young, H. A., Spitzer, J. H., Visintin, A. & Segal, D. M. Histamine regulates cytokine production in maturing dendritic cells, resulting in altered T cell polarization. J Clin Invest 108, 1865–1873 (2001). 10.1172/JCI13930

65 Pavlinkova, G., Yanagawa, Y., Kikuchi, K., Iwabuchi, K. & Onoe, K. Effects of histamine on functional maturation of dendritic cells. Immunobiology 207, 315–325 (2003). 10.1078/0171-2985-00247

66 Schenk, H., Neumann, D. & Kloth, C. Histamine regulates murine primary dendritic cell functions. Immunopharmacol Immunotoxicol 38, 379–384 (2016). 10.1080/08923973.2016.1214144

67 Aponte-Lopez, A. & Munoz-Cruz, S. Mast Cells in the Tumor Microenvironment. Adv Exp Med Biol 1273, 159–173 (2020). 10.1007/978-3-030-49270-0_9

68 Lichterman, J. N. & Reddy, S. M. Mast Cells: A New Frontier for Cancer Immunotherapy. Cells 10 (2021). 10.3390/cells10061270

69 Kwon, J. Y., Moskwa, N., Kang, W., Fan, T. M. & Lee, C. Canine as a Comparative and Translational Model for Human Mammary Tumor. J Breast Cancer 26, 1–13 (2023). 10.4048/jbc.2023.26.e4

70 Panula, P. Histamine receptors, agonists, and antagonists in health and disease. Handb Clin Neurol 180, 377–387 (2021). 10.1016/B978-0-12-820107-7.00023-9

71 Zampeli, E. & Tiligada, E. The role of histamine H4 receptor in immune and inflammatory disorders. Br J Pharmacol 157, 24–33 (2009). 10.1111/j.1476-5381.2009.00151.x

72 Sarasola, M. P., Taquez Delgado, M. A., Nicoud, M. B. & Medina, V. A. Histamine in cancer immunology and immunotherapy. Current status and new perspectives. Pharmacol Res Perspect 9, e00778 (2021). 10.1002/prp2.778

73 Trybus, E. & Trybus, W. H1 Antihistamines-Promising Candidates for Repurposing in the Context of the Development of New Therapeutic Approaches to Cancer Treatment. Cancers (Basel) 16 (2024). 10.3390/cancers16244253

74 Karlsson, M., et al. A single-cell type transcriptomics map of human tissues. Sci Adv 7 (2021). 10.1126/sciadv.abh2169

75 Banovac, K., Neylan, D., Leone, J., Ghandur-Mnaymneh, L. & Rabinovitch, A. Are the mast cells antigen presenting cells? Immunol Invest 18, 901–906 (1989). 10.3109/08820138909050768

76 Conti, P., et al. Mast Cells Accumulate in the Stroma of Breast Adenocarcinoma and Secrete Pro-Inflammatory Cytokines and Tumor-Damaging Mediators: Could IL-37 and IL-38 Play an Anti-Tumor Role? Int J Mol Sci 27 (2026). 10.3390/ijms27020824

77 Zhou, L., et al. The critical role of platelet in cancer progression and metastasis. Eur J Med Res 28, 385 (2023). 10.1186/s40001-023-01342-w

78 Ward, M. P., et al. Circulating tumour cells are a prognostic indicator in advanced high-grade serous ovarian cancer and are associated with platelets and immune cells following dissemination. Br J Cancer 134, 22–32 (2026). 10.1038/s41416-025-03227-7

79 Guan, C., Hao, L., Gong, B. & Zheng, L. CTC cluster in breast cancer: synergetic metastasis promotion mechanism and transformation path from platelet cloaking to leukocyte escort. Clin Transl Oncol (2025). 10.1007/s12094-025-04150-2

80 Wang, X., Zhao, S., Wang, Z. & Gao, T. Platelets involved tumor cell EMT during circulation: communications and interventions. Cell Commun Signal 20, 82 (2022). 10.1186/s12964-022-00887-3

81 Ward, M. P., et al. Platelets, immune cells and the coagulation cascade; friend or foe of the circulating tumour cell? Mol Cancer 20, 59 (2021). 10.1186/s12943-021-01347-1

82 Xu, X. R., Yousef, G. M. & Ni, H. Cancer and platelet crosstalk: opportunities and challenges for aspirin and other antiplatelet agents. Blood 131, 1777–1789 (2018). 10.1182/blood-2017-05-743187

83 Labelle, M., Begum, S. & Hynes, R. O. Direct signaling between platelets and cancer cells induces an epithelial-mesenchymal-like transition and promotes metastasis. Cancer Cell 20, 576–590 (2011). 10.1016/j.ccr.2011.09.009

84 Filippelli, A., et al. Scoping Review on Platelets and Tumor Angiogenesis: Do We Need More Evidence or Better Analysis? Int J Mol Sci 23 (2022). 10.3390/ijms232113401

85 Liao, K., et al. The role of platelets in the regulation of tumor growth and metastasis: the mechanisms and targeted therapy. MedComm (2020) 4, e350 (2023). 10.1002/mco2.350

86 Perdomo, J., et al. Neutrophil activation and NETosis are the major drivers of thrombosis in heparin-induced thrombocytopenia. Nat Commun 10, 1322 (2019). 10.1038/s41467-019-09160-7

87 Taifour, T., et al. The tumor-derived cytokine Chi3l1 induces neutrophil extracellular traps that promote T cell exclusion in triple-negative breast cancer. Immunity (2023). 10.1016/j.immuni.2023.11.002

88 Yeh, A. C. & Ramaswamy, S. Mechanisms of Cancer Cell Dormancy—Another Hallmark of Cancer? Cancer Research 75, 5014–5022 (2015). 10.1158/0008-5472.Can-15-1370

89 Coffelt, S. B., et al. IL-17-producing gammadelta T cells and neutrophils conspire to promote breast cancer metastasis. Nature 522, 345–348 (2015). 10.1038/nature14282

90 Sharma, A., et al. Cholesterol-targeting Wnt–β-catenin signaling inhibitors for colorectal cancer. Nature Chemical Biology (2025). 10.1038/s41589-025-01870-y

91 Ono, M., Toyomoto, M., Yamauchi, M. & Hagiwara, M. Platelets accelerate lipid peroxidation and induce pathogenic neutrophil extracellular trap release. Cell Chemical Biology 31, 2085–2095.e2084 (2024). 10.1016/j.chembiol.2024.11.003

92 NaveenKumar, S. K., et al. Platelet activation and ferroptosis mediated NETosis drives heme induced pulmonary thrombosis. Biochimica et Biophysica Acta (BBA) - Molecular Basis of Disease 1869, 166688 (2023). 10.1016/j.bbadis.2023.166688

93 Constantinescu-Bercu, A., et al. Activated αIIbβ3 on platelets mediates flow-dependent NETosis via SLC44A2. eLife 9, e53353 (2020). 10.7554/eLife.53353

94 Kött, J., et al. Synergistic effects of anticoagulants and platelet aggregation inhibitors with immune checkpoint inhibitors in cancer therapy: a comprehensive review of preclinical and clinical evidence. Journal for ImmunoTherapy of Cancer 14, e013879 (2026). 10.1136/jitc-2025-013879

95 Majorini, M. T., Colombo, M. P. & Lecis, D. Few, but Efficient: The Role of Mast Cells in Breast Cancer and Other Solid Tumors. Cancer Res 82, 1439–1447 (2022). 10.1158/0008-5472.CAN-21-3424

96 Rodewald, H. R. & Feyerabend, T. B. Widespread immunological functions of mast cells: fact or fiction? Immunity 37, 13–24 (2012). 10.1016/j.immuni.2012.07.007

97 Suurmond, J., et al. Communication between human mast cells and CD4(+) T cells through antigen-dependent interactions. Eur J Immunol 43, 1758–1768 (2013). 10.1002/eji.201243058

98 Gaudenzio, N., Laurent, C., Valitutti, S. & Espinosa, E. Human mast cells drive memory CD4+ T cells toward an inflammatory IL-22+ phenotype. J Allergy Clin Immunol 131, 1400–1407 e1411 (2013). 10.1016/j.jaci.2013.01.029

99 Lotfi-Emran, S., et al. Human mast cells present antigen to autologous CD4(+) T cells. J Allergy Clin Immunol 141, 311–321 e310 (2018). 10.1016/j.jaci.2017.02.048

100 Wu, S. Y., et al. Mobilizing antigen-presenting mast cells in anti-PD-1-refractory triple-negative breast cancer: a phase 2 trial. Nat Med 31, 2405–2415 (2025). 10.1038/s41591-025-03776-7

101 Ribatti, D., Annese, T. & Tamma, R. Controversial role of mast cells in breast cancer tumor progression and angiogenesis. Clin Breast Cancer 21, 486–491 (2021). 10.1016/j.clbc.2021.08.010

102 Aponte-Lopez, A., Fuentes-Panana, E. M., Cortes-Munoz, D. & Munoz-Cruz, S. Mast Cell, the Neglected Member of the Tumor Microenvironment: Role in Breast Cancer. J Immunol Res 2018, 2584243 (2018). 10.1155/2018/2584243

103 Keser, S. H., et al. Relationship of mast cell density with lymphangiogenesis and prognostic parameters in breast carcinoma. Kaohsiung J Med Sci 33, 171–180 (2017). 10.1016/j.kjms.2017.01.005

104 Rajput, A. B., et al. Stromal mast cells in invasive breast cancer are a marker of favourable prognosis: a study of 4,444 cases. Breast cancer research and treatment 107, 249–257 (2008). 10.1007/s10549-007-9546-3

105 Dabiri, S., et al. The presence of stromal mast cells identifies a subset of invasive breast cancers with a favorable prognosis. Mod Pathol 17, 690–695 (2004). 10.1038/modpathol.3800094

106 Bowers, H. M., Jr., Mahapatro, R. C. & Kennedy, J. W. Numbers of mast cells in the axillary lymph nodes of breast cancer patients. Cancer 43, 568–573 (1979). 10.1002/1097-0142(197902)43:2<568::aid-cncr2820430225>3.0.co;2-#

107 Bendre, S. V. et al. Cholesterol efflux protein, ABCA1, supports anticancer functions of myeloid immune cells. Sci Adv 12, eadx5490 (2026). 10.1126/sciadv.adx5490

108 Greene, J. T., Brian, B. F. t., Senevirathne, S. E. & Freedman, T. S. Regulation of myeloid-cell activation. Curr Opin Immunol 73, 34–42 (2021). 10.1016/j.coi.2021.09.004

109 Coats, B. R., et al. Metabolically Activated Adipose Tissue Macrophages Perform Detrimental and Beneficial Functions during Diet-Induced Obesity. Cell reports 20, 3149–3161 (2017). 10.1016/j.celrep.2017.08.096

110 Naik, R., Baliga, P., Bansal, R. & Pai, M. Distribution of mast cells in the axillary lymph nodes of breast cancer patients. J Indian Med Assoc 95, 606–607 (1997).

111 Ranieri, G., et al. Tryptase-positive mast cells correlate with angiogenesis in early breast cancer patients. Int J Oncol 35, 115–120 (2009). 10.3892/ijo_00000319

112 Somasundaram, R., et al. Tumor-infiltrating mast cells are associated with resistance to anti-PD-1 therapy. Nat Commun 12, 346 (2021). 10.1038/s41467-020-20600-7

113 Korashy, H. M., et al. Sunitinib Inhibits Breast Cancer Cell Proliferation by Inducing Apoptosis, Cell-cycle Arrest and DNA Repair While Inhibiting NF-kappaB Signaling Pathways. Anticancer research 37, 4899–4909 (2017). 10.21873/anticanres.11899

114 Samei, L., Yaling, P., Lihua, Y., Yan, Z. & Shuyan, J. Effects and Mechanism of Imatinib in Inhibiting Colon Cancer Cell Proliferation. Med Sci Monit 22, 4126–4131 (2016). 10.12659/msm.898152

115 Nguyen, P. L. & Cho, J. Pathophysiological Roles of Histamine Receptors in Cancer Progression: Implications and Perspectives as Potential Molecular Targets. Biomolecules 11 (2021). 10.3390/biom11081232

116 Azimi, H., Jafari, A., Maralani, M. & Davoodi, H. The role of histamine and its receptors in breast cancer: from pathology to therapeutic targets. Med Oncol 41, 190 (2024). 10.1007/s12032-024-02437-y

117 Yang, X. D., et al. Histamine deficiency promotes inflammation-associated carcinogenesis through reduced myeloid maturation and accumulation of CD11b+Ly6G+ immature myeloid cells. Nat Med 17, 87–95 (2011). 10.1038/nm.2278

118 Merrill, R. M., Isakson, R. T. & Beck, R. E. The association between allergies and cancer: what is currently known? Ann Allergy Asthma Immunol 99, 102–116; quiz 117-109, 150 (2007). 10.1016/S1081-1206(10)60632-1

119 Hedderson, M. M., Malone, K. E., Daling, J. R. & White, E. Allergy and risk of breast cancer among young women (United States). Cancer Causes Control 14, 619–626 (2003). 10.1023/a:1025686913626

120 Kallen, B., Gunnarskog, J. & Conradson, T. B. Cancer risk in asthmatic subjects selected from hospital discharge registry. Eur Respir J 6, 694–697 (1993).

121 Ji, J., et al. Cancer risk in hospitalised asthma patients. British Journal of Cancer 100, 829–833 (2009). 10.1038/sj.bjc.6604890

122 Li, H., et al. The allergy mediator histamine confers resistance to immunotherapy in cancer patients via activation of the macrophage histamine receptor H1. Cancer Cell 40, 36–52 e39 (2022). 10.1016/j.ccell.2021.11.002

123 Coogan, P. F., Zhang, Y., Palmer, J. R., Strom, B. L. & Rosenberg, L. Cimetidine and other histamine2-receptor antagonist use in relation to risk of breast cancer. Cancer Epidemiol Biomarkers Prev 14, 1012–1015 (2005). 10.1158/1055-9965.EPI-04-0547

124 Habel, L. A., Levin, T. R. & Friedman, G. D. Cimetidine use and risk of breast, prostate, and other cancers. Pharmacoepidemiol Drug Saf 9, 149–155 (2000). 10.1002/(SICI)1099-1557(200003/04)9:2<149::AID-PDS481>3.0.CO;2-1

125 Rossing, M. A., Scholes, D., Cushing-Haugen, K. L. & Voigt, L. F. Cimetidine use and risk of prostate and breast cancer. Cancer Epidemiol Biomarkers Prev 9, 319–323 (2000).

126 Meghnem, D., Oldford, S. A., Haidl, I. D., Barrett, L. & Marshall, J. S. Histamine receptor 2 blockade selectively impacts B and T cells in healthy subjects. Sci Rep 11, 9405 (2021). 10.1038/s41598-021-88829-w

127 Nicoud, M. B., et al. Study of the antitumour effects and the modulation of immune response by histamine in breast cancer. Br J Cancer 122, 348–360 (2020). 10.1038/s41416-019-0636-x

128 Crouchet, E., et al. A human liver cell-based system modeling a clinical prognostic liver signature for therapeutic discovery. Nat Commun 12, 5525 (2021). 10.1038/s41467-021-25468-9

129 Győrffy, B. Survival analysis across the entire transcriptome identifies biomarkers with the highest prognostic power in breast cancer. Comput Struct Biotechnol J 19, 4101–4109 (2021). 10.1016/j.csbj.2021.07.014

130 Tang, Z., Kang, B., Li, C., Chen, T. & Zhang, Z. GEPIA2: an enhanced web server for large-scale expression profiling and interactive analysis. Nucleic Acids Res 47, W556–W560 (2019). 10.1093/nar/gkz430

131 Nelczyk, A. T., et al. The nuclear receptor TLX (NR2E1) inhibits growth and progression of triple- negative breast cancer. Biochim Biophys Acta Mol Basis Dis 1868, 166515 (2022). 10.1016/j.bbadis.2022.166515

132 Nelson, E. R. et al. The oxysterol, 27-hydroxycholesterol, links cholesterol metabolism to bone homeostasis through its actions on the estrogen and liver X receptors. Endocrinology 152, 4691–4705 (2011). 10.1210/en.2011-1298

133 Tsvilovskyy, V., Solis-Lopez, A., Ohlenschlager, K. & Freichel, M. Isolation of Peritoneum-derived Mast Cells and Their Functional Characterization with Ca2+-imaging and Degranulation Assays. J Vis Exp (2018). 10.3791/57222

134 Vidana Gamage, H. E., et al. Development of NR0B2 as a therapeutic target for the re-education of tumor associated myeloid cells. Cancer letters 597, 217086 (2024). 10.1016/j.canlet.2024.217086

135 Vidana Gamage, H. E., et al. NR0B2 re-educates myeloid immune cells to reduce regulatory T cell expansion and progression of breast and other solid tumors. Cancer letters 597, 217042 (2024). 10.1016/j.canlet.2024.217042

136 Lopez-Sanz, C., Sanchez-Martinez, E. & Jimenez-Saiz, R. Protocol to desensitize human and murine mast cells after polyclonal IgE sensitization. STAR Protoc 3, 101755 (2022). 10.1016/j.xpro.2022.101755

137 Rothschild, A. M. Mechanisms of histamine release by compound 48-80. Br J Pharmacol 38, 253–262 (1970). 10.1111/j.1476-5381.1970.tb10354.x

138 Adamczyk-Engelmann, P. & Gietzen, K. Induction of histamine release and calmodulin antagonism are two distinct properties of compound 48/80. Cell Calcium 10, 93–99 (1989). 10.1016/0143-4160(89)90049-3

139 Kashem, S. W., et al. G protein coupled receptor specificity for C3a and compound 48/80-induced degranulation in human mast cells: roles of Mas-related genes MrgX1 and MrgX2. Eur J Pharmacol 668, 299–304 (2011). 10.1016/j.ejphar.2011.06.027

140 Nelson, E. R. & Habibi, H. R. Functional significance of a truncated thyroid receptor subtype lacking a hormone-binding domain in goldfish. Endocrinology 149, 4702–4709 (2008). 10.1210/en.2008-0107

141 Jeffries, K. M., Nelson, E. R., Jackson, L. J. & Habibi, H. R. Basin-wide impacts of compounds with estrogen-like activity on longnose dace (Rhinichthys cataractae) in two prairie rivers of Alberta, Canada. Environ Toxicol Chem 27, 2042–2052 (2008). 10.1897/07-529.1

142 Geiss, G. K., et al. Direct multiplexed measurement of gene expression with color-coded probe pairs. Nat Biotechnol 26, 317–325 (2008). 10.1038/nbt1385

143 Peterson, B. C. Effects of brine temperature on ham and bacon processing characteristics. Thesis Disertation Animal Sciences, University of Illinois at Urbana-Champaign (2016).

144 Reeves, P. G., Nielsen, F. H. & Fahey, G. C., Jr. AIN-93 purified diets for laboratory rodents: final report of the American Institute of Nutrition ad hoc writing committee on the reformulation of the AIN-76A rodent diet. J Nutr 123, 1939–1951 (1993). 10.1093/jn/123.11.1939

145 Cam, A., et al. Thermally Abused Frying Oil Potentiates Metastasis to Lung in a Murine Model of Late-Stage Breast Cancer. Cancer Prev Res (Phila) 12, 201–210 (2019). 10.1158/1940-6207.CAPR-18-0220

146 Quail, D. F., et al. Obesity alters the lung myeloid cell landscape to enhance breast cancer metastasis through IL5 and GM-CSF. Nat Cell Biol 19, 974–987 (2017). 10.1038/ncb3578

147 Ge, S. X., Jung, D. & Yao, R. ShinyGO: a graphical gene-set enrichment tool for animals and plants. Bioinformatics 36, 2628–2629 (2020). 10.1093/bioinformatics/btz931

148 Lanczky, A. & Gyorffy, B. Web-Based Survival Analysis Tool Tailored for Medical Research (KMplot): Development and Implementation. J Med Internet Res 23, e27633 (2021). 10.2196/27633

149 Uhlen, M., et al. A pathology atlas of the human cancer transcriptome. Science 357 (2017). 10.1126/science.aan2507

150 Han, Y., et al. TISCH2: expanded datasets and new tools for single-cell transcriptome analyses of the tumor microenvironment. Nucleic Acids Res 51, D1425–D1431 (2023). 10.1093/nar/gkac959

