## Supplementary Material for "Loss of Mast cells and histaminergic signaling link diet to platelet-mediated NETosis and mammary cancer recurrence"

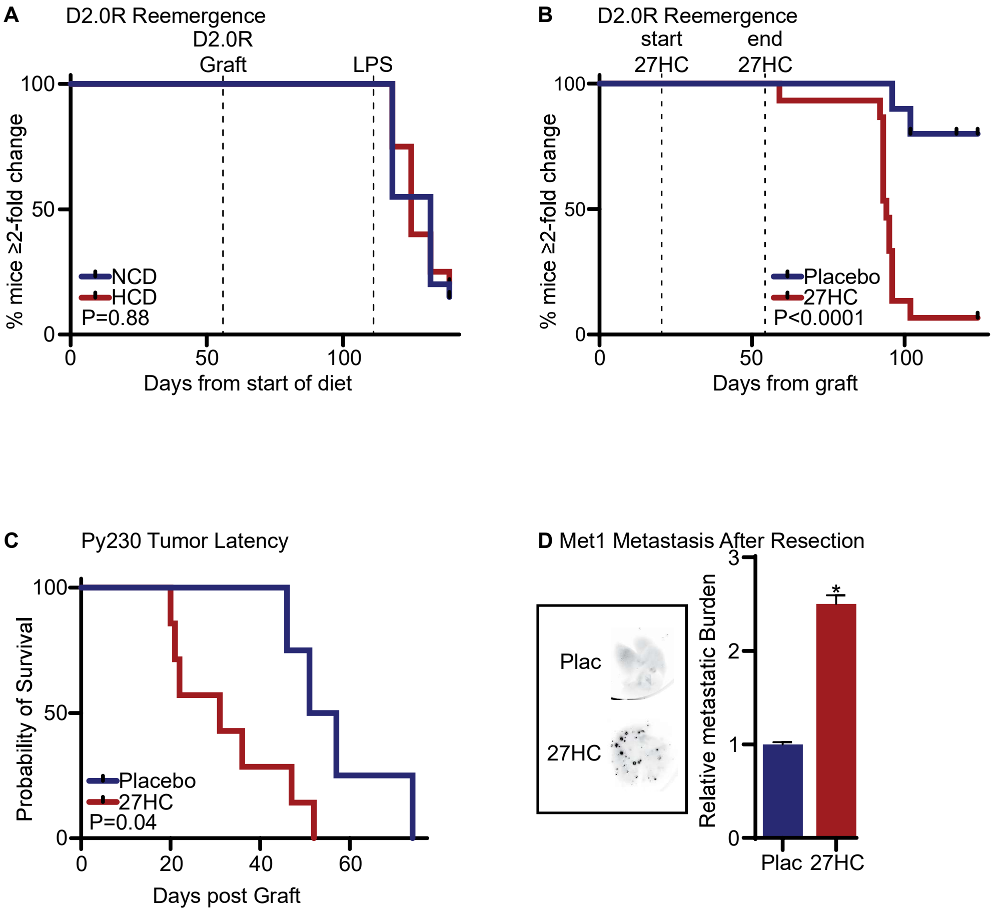
A
D2.0R Reemergence
D2.0R Graft
LPS
% mice ≥2-fold change
100
50
0
NCD
HCD
P=0.88
0
50
100
Days from start of diet
B
D2.0R Reemergence
start 27HC
end 27HC
% mice ≥2-fold change
100
50
0
Placebo
27HC
P<0.0001
0
50
100
Days from graft
C
Py230 Tumor Latency
Probability of Survival
100
50
0
Placebo
27HC
P=0.04
0
20
40
60
Days post Graft
D Met1 Metastasis After Resection
Plac
27HC
Relative metastatic Burden
3
2
1
0
*
Plac
27HC

SFig 1

**Supplementary Figure 1:** ***Dietary cholesterol does not influence reemergence from dormancy, while its oxidized metabolite, 27-hydroxycholesterol (27HC) does.***

**(A)** Mice were placed on a control, normal cholesterol diet (NCD) or high cholesterol diet (HCD) for 8 weeks prior to graft with D2.0R cells to model dormant lesions. LPS was administered to spark reemergence. A doubling of metastatic burden (luciferase signal) was considered breaking of dormancy. No significant differences in recurrence were observed between groups (P value from Logrank test, N=40). **(B)** D2.0R cells were grafted into mice to model dormant lesions. 20 days after graft, mice were treated with either placebo or 20mg/kg 27HC, daily through 55 days post-graft (treatment window indicated by dashed lines), and then followed through time. A doubling of metastatic burden (luciferase signal) was considered breaking of dormancy (P value from Logrank test, N=25). **(C)** Py230 cells were grafted orthotopically. These grafts have a very long latency prior to detection of a palpable tumor, making them a model of dormancy. Treatment with placebo or 27HC commenced 3d after graft. Time to detection of a palpable tumor is shown (P value from Logrank test, N=12). **(D)** Met1 cells were grafted and tumors allowed to establish to ~200mm$^3$, at which point they were surgically removed and mice treated with three rounds of doxorubicin (dosed every 2d). Mice were then treated with placebo or 27HC for 21 days. Lungs were then imaged *ex vivo*; representative images to the left of quantified data. (unpaired t test, N=16).

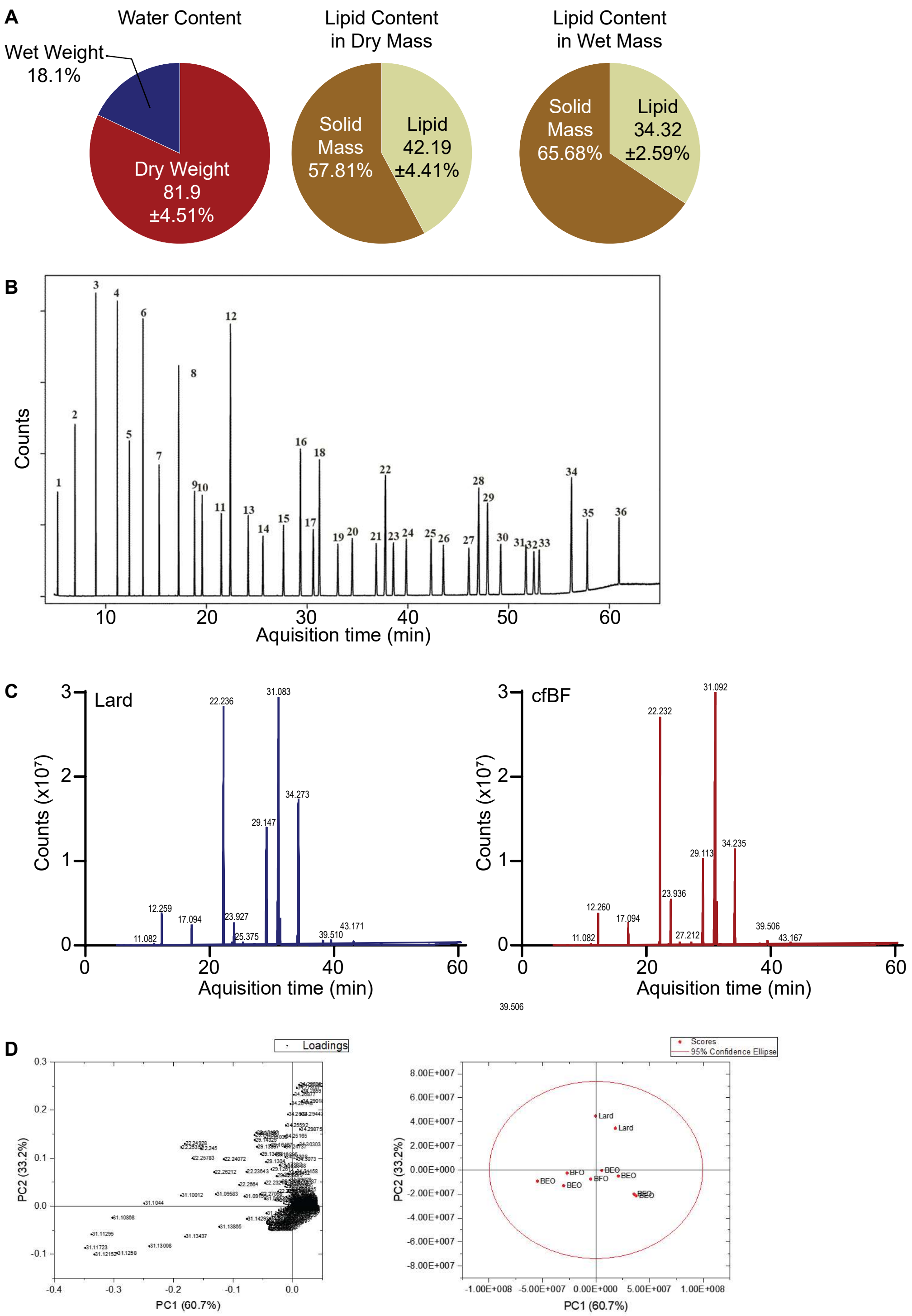
A
Water Content
Wet Weight
18.1%
Dry Weight
81.9
±4.51%
Lipid Content
in Dry Mass
Solid
Mass
57.81%
Lipid
42.19
±4.41%
Lipid Content
in Wet Mass
Solid
Mass
65.68%
Lipid
34.32
±2.59%
B
Counts
Aquisition time (min)
C
Lard
cfBF
Counts (x10⁷)
Aquisition time (min)
22.236
31.083
34.273
29.147
12.259
17.094
23.927
25.375
11.082
39.510
43.171
31.092
22.232
34.235
29.113
23.936
12.260
17.094
27.212
11.082
39.506
43.167
39.506
D
Loadings
PC1 (60.7%)
PC2 (33.2%)
Scores
95% Confidence Ellipse
Lard
BEO
BFO

SFig 2

**Supplementary Figure 2:** ***Fatty acid characterization of fat from cured, fried bacon (cfBF) or rendered lard.***

**(A)** Bacon content was characterized for water and lipid content after pan-frying. The left two pie charts are based on dried samples, whereas the right pie chart is based on wet-bacon (undried). **(B)** Total ion chromatogram of 36 mixed fatty acid methyl esters (FAMEs) analyzed by GC-MS on an HP-88 column (60 m × 0.25 mm, 0.20 μm). The oven temperature program was as follows: initial temperature of 60 °C for 1 min, ramped at 10 °C/min to 145 °C, then at 1 °C/min to 190 °C, and finally at 5 °C/min to 220 °C. Mass spectrometry was operated in scan mode over an m/z range of 50–500 amu with an EI voltage of 70 eV. Compound identification is indicated in **Supplementary Table 1A**. **(C)** GC-MS chromatograms of fatty acid methyl esters of lard and cfBF. Compound identification and quantification shown in **Supplementary Table 1B**. **(D)** PCA plots of cfBF vs lard. Individual samples were plotted based on the first two principle components (PC1 and PC2), which accounts for 60.7% and 33.2% of the total variance, respectively. The clustering suggests a clear separation from cfBF and rendered lard.

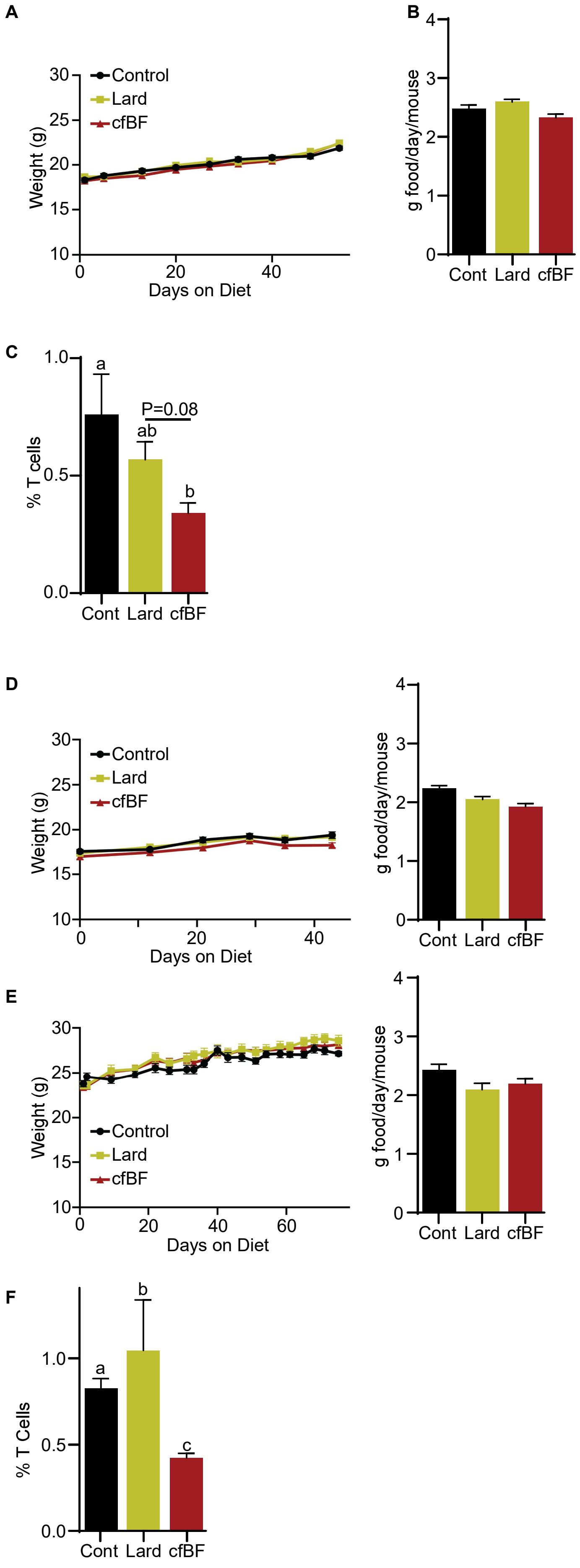
A
B
Control
Lard
cfBF
Weight (g)
Days on Diet
g food/day/mouse
Cont
Lard
cfBF
C
% T cells
a
ab
b
P=0.08
D
E
F
% T Cells
a
b
c

SFig 3

**Supplementary Figure 3:** ***Mice consuming diets enriched with lard or cured fried bacon fat (cfBF) do not have altered weight, but their tumors have decreased T cell infiltration*.**
**(A)** Mean body weights through time of 4T1 tumor-bearing mice consuming control (modified AIN-93G to contain 10% fat from soybean oil), lard (5% lard, 5% soybean oil) or cfBF (5% cfBF, 5% soybean oil) diets. Data is presented as mean mass ± SEM (N=40). **(B)** Mean food consumed per mouse per day. The data from A & B correspond to the data in **Fig. 1A** of the main text. **(C)** Flow cytometric analysis of primary tumors found decreased T cell infiltration in mice fed cfBF (N=39, 1-Way ANOVA followed by multiple comparison test of geometric means with Šidák's correction). **(D)** Left panel: Mean body weights through time of mice bearing 4T1 metastatic lesions, consuming control, lard or cfBF diets. Right Panel: Corresponding food consumption data. These data correspond to **Fig. 1B** of the main text. **(E)** Left panel: Mean body weights through time of mice bearing Met1 metastatic lesions, consuming control, lard or cfBF diets. Right Panel: Corresponding food consumption data. These data correspond to **Fig. 1C** of the main text. **(F)** Met1 metastatic lesions from mice consuming cfBF had decreased T cell infiltrate (N=34; 1-Way ANOVA followed by multiple comparison test of geometric means with Šidák's correction). Different letters denote statistically significant differences.

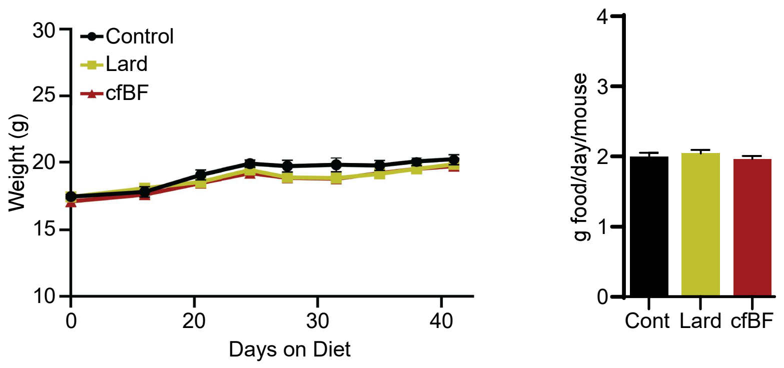
Control
Lard
cfBF
Weight (g)
Days on Diet
g food/day/mouse
Cont
Lard
cfBF

SFig 4

**Supplementary Figure 4:** ***Mouse body weights and food consumption are not different when consuming different experimental diets***. Left panel: Mean body weights through time of mice grafted with a sub-optimal number of 4T1 cells, consuming control, lard or cfBF diets. Right Panel: Corresponding food consumption data. (N=25) These data correspond to **Fig. 1D** of the main text.

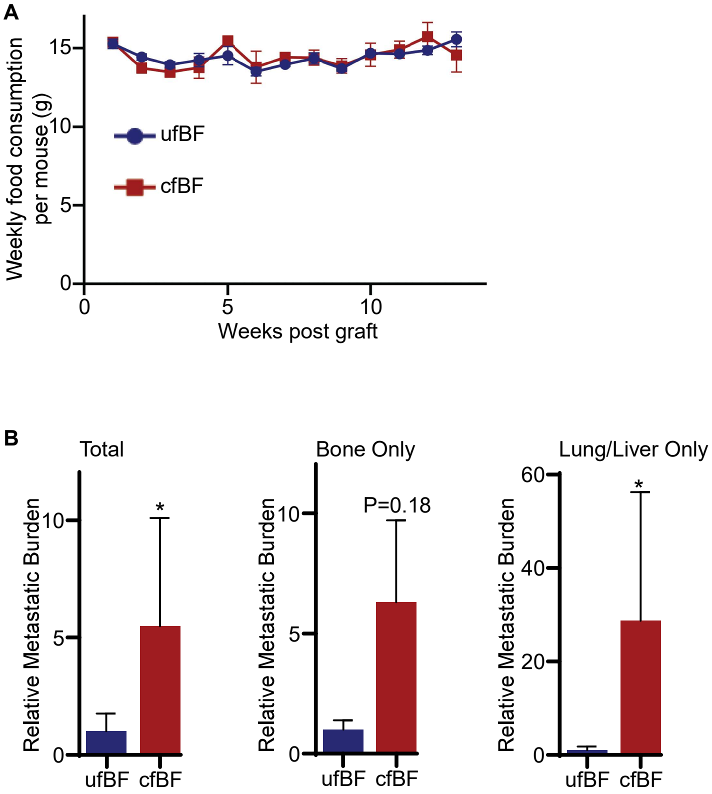
A
Weekly food consumption per mouse (g)
15
10
5
0
ufBF
cfBF
0
5
10
Weeks post graft
B
Total
Relative Metastatic Burden
10
5
0
*
ufBF
cfBF
Bone Only
Relative Metastatic Burden
10
5
0
P=0.18
ufBF
cfBF
Lung/Liver Only
Relative Metastatic Burden
60
40
20
0
*
ufBF
cfBF

SFig 5

**Supplementary Figure 5:** ***Mice consuming a diet enriched in uncured fried bacon fat (ufBF) or cfBF did not show altered food consumption, but significantly increased metastatic burden was observed in cfBF fed mice***. **(A)** Mean consumption of food per week comparing mice consuming ufBF or cfBF. **(B)** Fold change in metastatic burden of D2.0R bearing mice consuming ufBF or cfBF. Left: total metastatic burden. Middle: bone only. Right: Lung and liver only. These data correspond to **Fig. 1E** in the main text. Asterisks indicate statistical significance (Mann-Whitney U test or students T test, P<0.05).

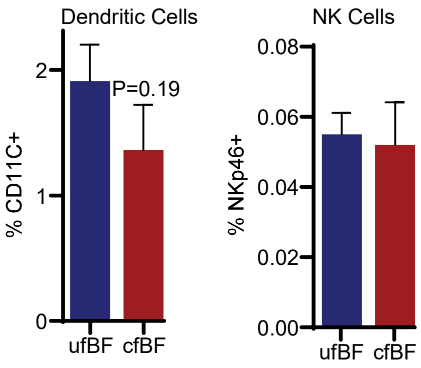
Dendritic Cells
% CD11C+
2
1
0
P=0.19
ufBF
cfBF
NK Cells
% NKp46+
0.08
0.06
0.04
0.02
0.00
ufBF
cfBF

SFig 6

**Supplementary Figure 6:** ***Flow cytometric analyses of lungs from naïve mice on different diets.*** Dendritic cells (CD11C+) and NK cells (NKp46+) were assessed by flow cytometry. These data correspond to **Fig. 1F** of the main text.

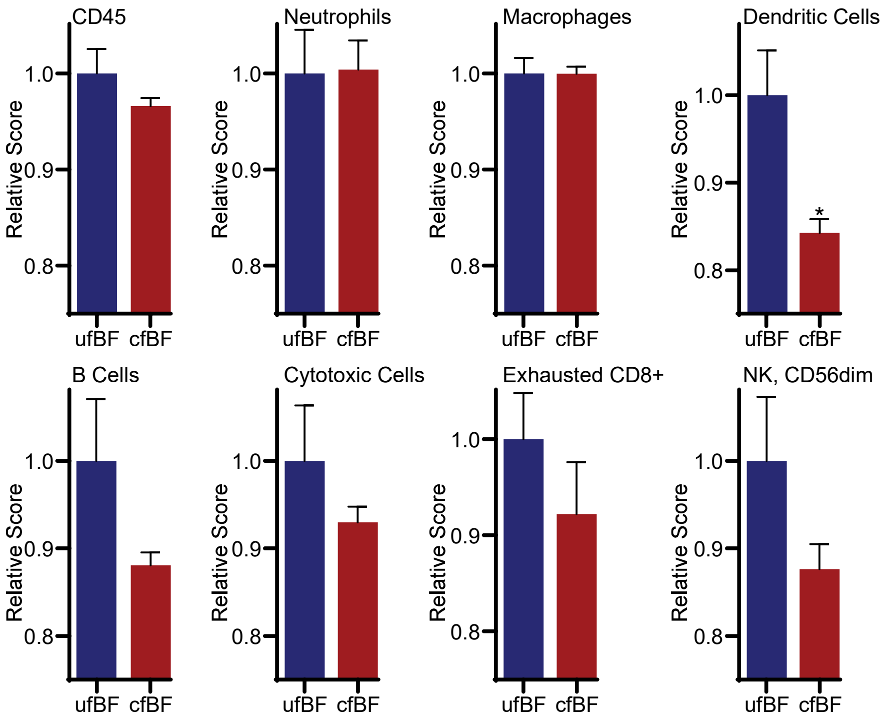
CD45
Neutrophils
Macrophages
Dendritic Cells
B Cells
Cytotoxic Cells
Exhausted CD8+
NK, CD56dim
Relative Score
1.0
0.9
0.8
*
ufBF
cfBF

SFig 7

**Supplementary Figure 7:** ***Analysis of Nanostring results predicts decreases in dendritic cells.*** Transcriptomics was performed using Nanostring. Informatic analyses predicted the relative abundance of different immune cell types. These data correspond to **Figs. 1E** and **2A** in the main text.

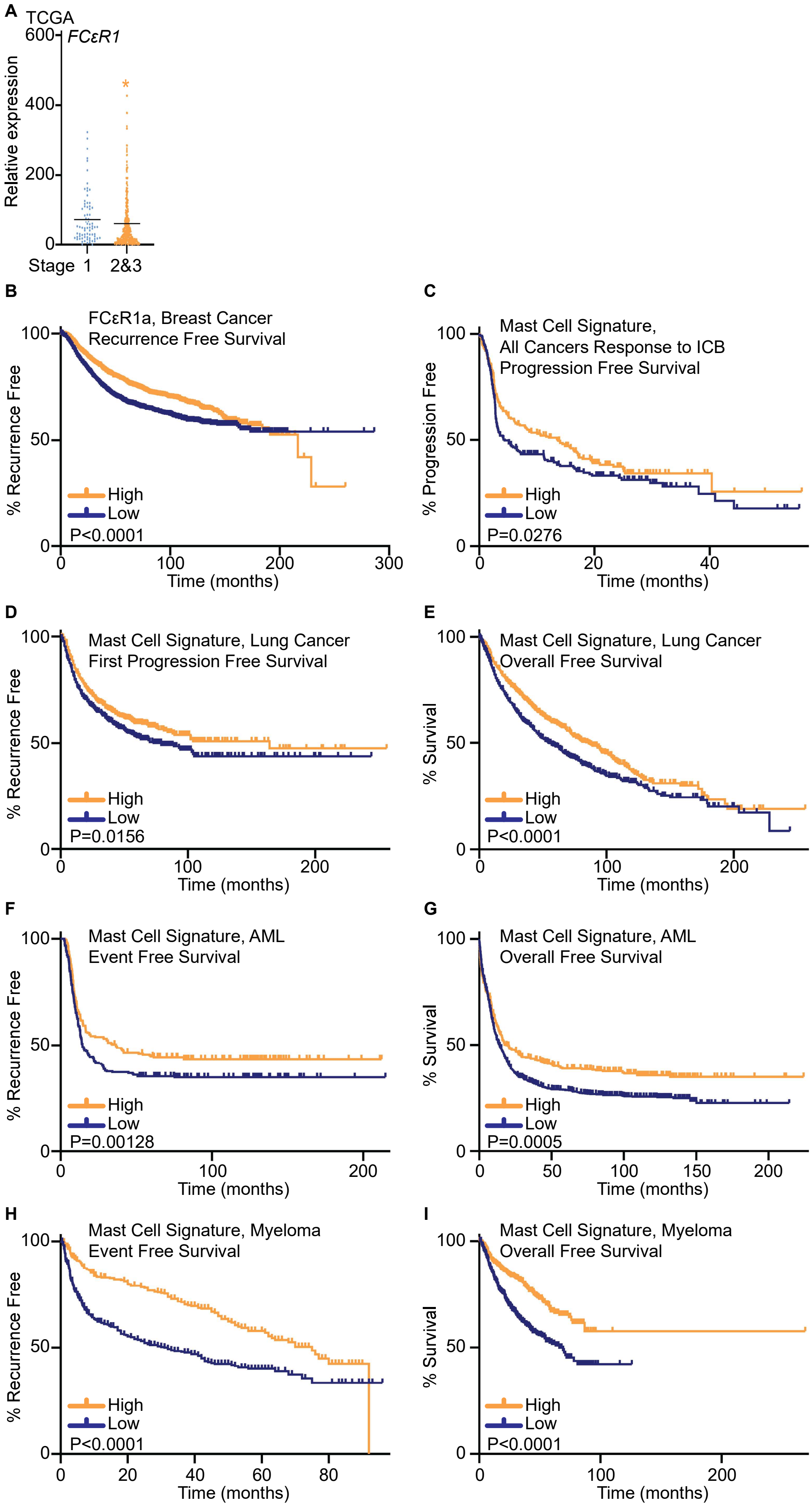

A
TCGA
FCεR1
Relative expression
600
400
200
0
Stage 1
2&3
*
B
FCεR1a, Breast Cancer
Recurrence Free Survival
% Recurrence Free
High
Low
P<0.0001
Time (months)
C
Mast Cell Signature,
All Cancers Response to ICB
Progression Free Survival
% Progression Free
High
Low
P=0.0276
Time (months)
D
Mast Cell Signature, Lung Cancer
First Progression Free Survival
% Recurrence Free
High
Low
P=0.0156
Time (months)
E
Mast Cell Signature, Lung Cancer
Overall Free Survival
% Survival
High
Low
P<0.0001
Time (months)
F
Mast Cell Signature, AML
Event Free Survival
% Recurrence Free
High
Low
P=0.00128
Time (months)
G
Mast Cell Signature, AML
Overall Free Survival
% Survival
High
Low
P=0.0005
Time (months)
H
Mast Cell Signature, Myeloma
Event Free Survival
% Recurrence Free
High
Low
P<0.0001
Time (months)
I
Mast Cell Signature, Myeloma
Overall Free Survival
% Survival
High
Low
P<0.0001
Time (months)


SFig 8

**Supplementary Figure 8:** ***Expression of genes associated with MCs are a good prognostic across several tumor types***.

**(A)** *FCεR1a*, a gene selectively expressed on MCs, is decreased in human breast tumors with higher stage compared to lower. Data from the TCGA Invasive Breast Cancer Firehose Legacy archive, binned into tumors of stage 1, or 2&3 (unpaired t test, N=471). **(B)** Elevated expression of a *FCεR1a* in human breast tumors is associated with increased recurrence free survival (RFS). All breast cancer subtypes considered here. Data accessed from the Kaplan-Meier Plotter webtool based on data from GEO, EGA, and TCGA (P value indicated from Logrank test, N=4890). The Kaplan-Meier Plotter webtool uses aggregated data from GEO, EGA, and TCGA[147]. **(C)** Elevated expression of the 'MC-gene signature' in tumors from patients treated with immune checkpoint blockers (ICB) is associated with an increased progression free survival time. Tumor types included for this analysis were bladder, esophageal adenocarcinoma, glioblastoma, hepatocellular carcinoma, head and neck squamous cell carcinoma, melanoma, non-small cell lung cancer and urothelial cancer. (P value from Logrank test, N=463). **(D-E)** Elevated expression of the 'MC-gene signature' in lung cancer tumors is associated with longer recurrence free- and overall- survival (RFS and OS respectively, P value from Logrank test, RFS N=1218, OS N=2154). **(F-G)** Elevated expression of the 'MC-gene signature' in acute myeloid leukemia (AML) is associated with longer event free- and overall- survival (P value from Logrank test, EFS N=525, OS N=1603). **(H-I)** Elevated expression of the 'MC-gene signature' in multiple myeloma samples is associated with RFS and OS (P value from Logrank test, RFS N=797, OS N=1415).

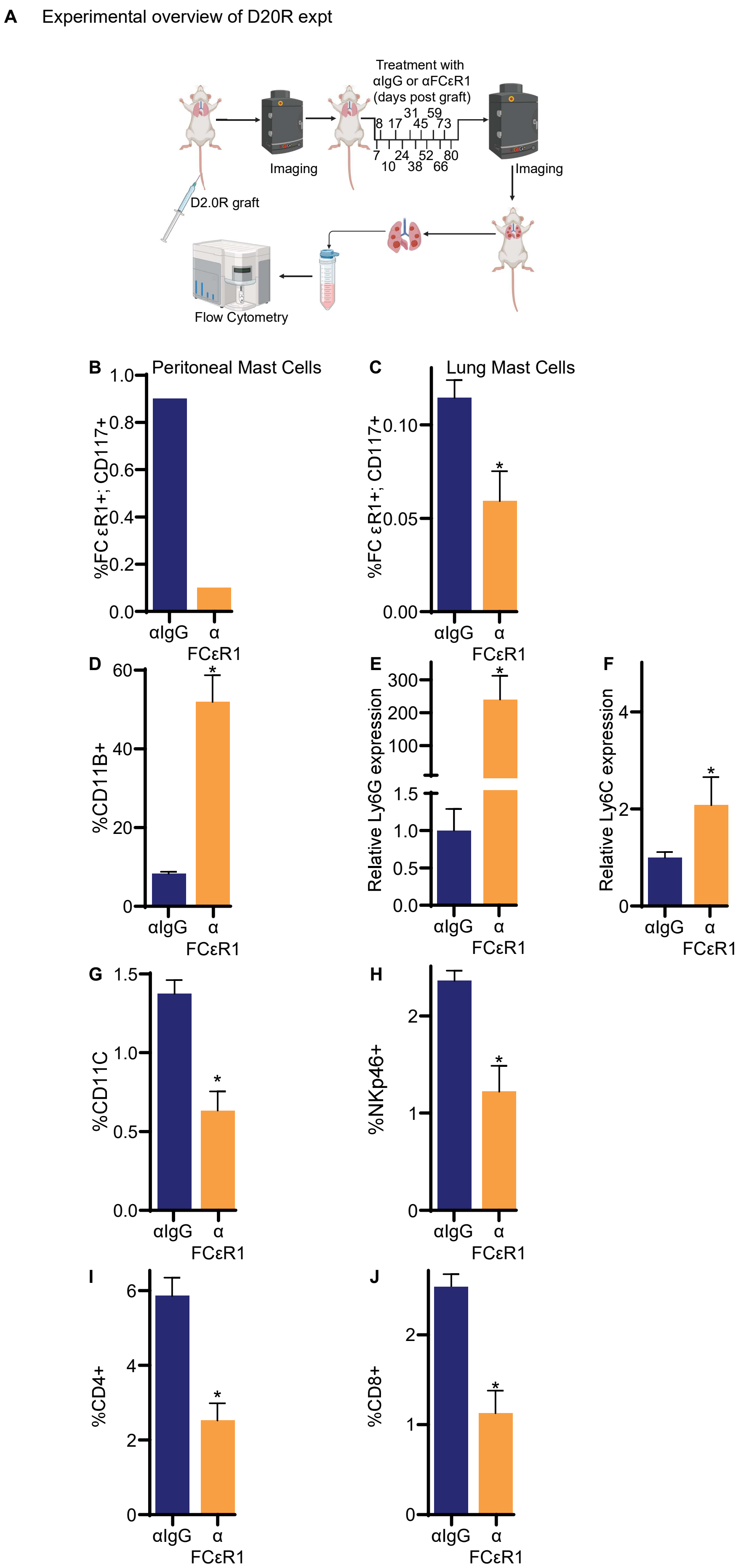

A
Experimental overview of D20R expt
Treatment with
αIgG or αFCεR1
(days post graft)
8 17 31 45 59 73
7 10 24 38 52 66 80
Imaging
Imaging
D2.0R graft
Flow Cytometry
B
Peritoneal Mast Cells
%FC εR1+; CD117+
1.0
0.8
0.6
0.4
0.2
0.0
αIgG
α FCεR1
C
Lung Mast Cells
%FC εR1+; CD117+
0.10
0.05
0.00
*
αIgG
α FCεR1
D
%CD11B+
60
40
20
0
*
αIgG
α FCεR1
E
Relative Ly6G expression
300
200
100
1.5
1.0
0.5
0.0
*
αIgG
α FCεR1
F
Relative Ly6C expression
4
2
0
*
αIgG
α FCεR1
G
%CD11C
1.5
1.0
0.5
0.0
*
αIgG
α FCεR1
H
%NKp46+
2
1
0
*
αIgG
α FCεR1
I
%CD4+
6
4
2
0
*
αIgG
α FCεR1
J
%CD8+
2
1
0
*
αIgG
α FCεR1


SFig 9

**Supplementary Figure 9:** ***Loss of mast cells (MCs) results in recurrence from mammary cancer dormancy***.

**(A)** Overview of experimental design, corresponding to **Fig. 2H**. **(B)** Administration of an antibody against FCεR1 (αFCεR1) successfully immune-depletes peritoneal MCs. **(C)** αFCεR1 depletes MCs within D2.0R bearing lungs. Samples taken at the end of the study (Day 84, N=35). **(D-J)** Flow cytometric or qPCR analysis of immune cell populations within D2.0R bearing lungs. (D, G-J are flow cytometry, N=32-35; E-F are qPCR, N=33-35 (unpaired T test).

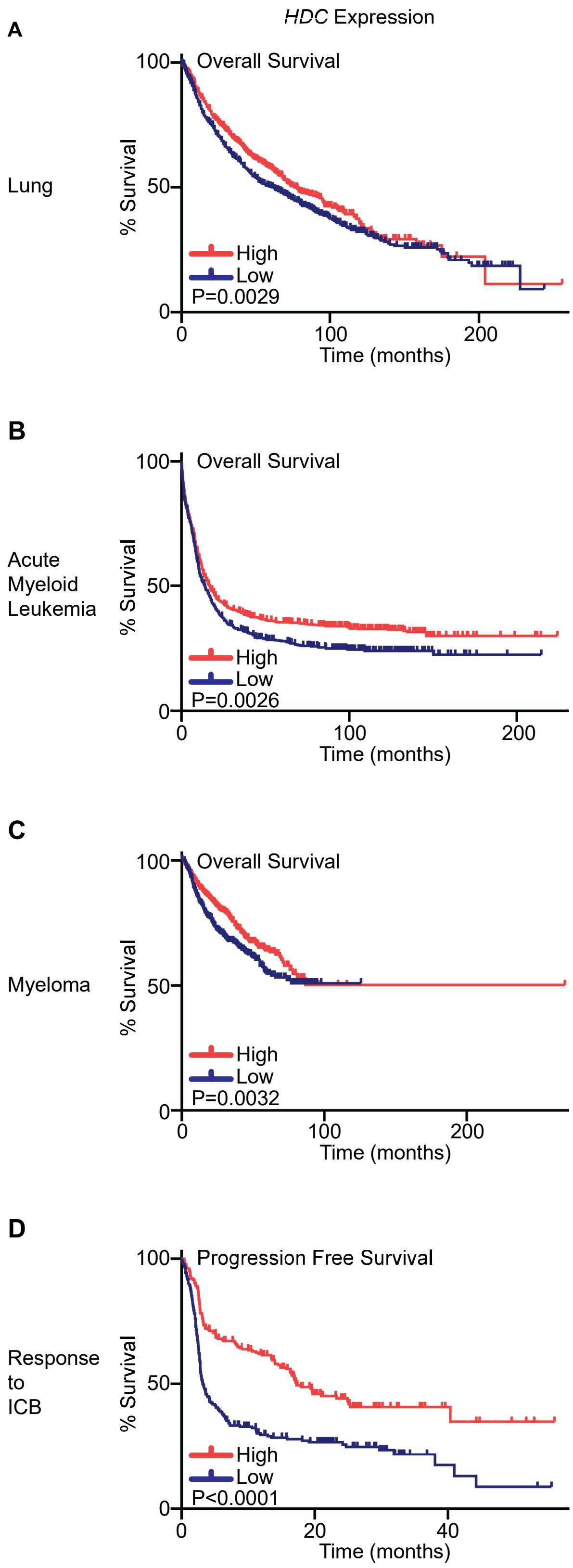
HDC Expression
A
Lung
Overall Survival
100
50
0
% Survival
High
Low
P=0.0029
0
100
200
Time (months)
B
Acute
Myeloid
Leukemia
Overall Survival
100
50
0
% Survival
High
Low
P=0.0026
0
100
200
Time (months)
C
Myeloma
Overall Survival
100
50
0
% Survival
High
Low
P=0.0032
0
100
200
Time (months)
D
Response
to
ICB
Progression Free Survival
100
50
0
% Survival
High
Low
P<0.0001
0
20
40
Time (months)

SFig 10

**Supplementary Figure 10:** ***HDC expression in human tumors is associated with improved survival***. Histidine decarboxylase (*HDC)* expression in **(A)** lung, **(B)** acute myeloid leukemia, or **(C)** myeloma is associated with improved overall survival. **(D)** Elevated *HDC* expression in tumors is associated with improved response to ICB. Data accessed from the Kaplan-Meier Plotter webtool based on data from GEO, EGA, and TCGA (P values indicated from Logrank test, A: N=2154 , B: N=1603 , C: N=1416 , D: N=463).

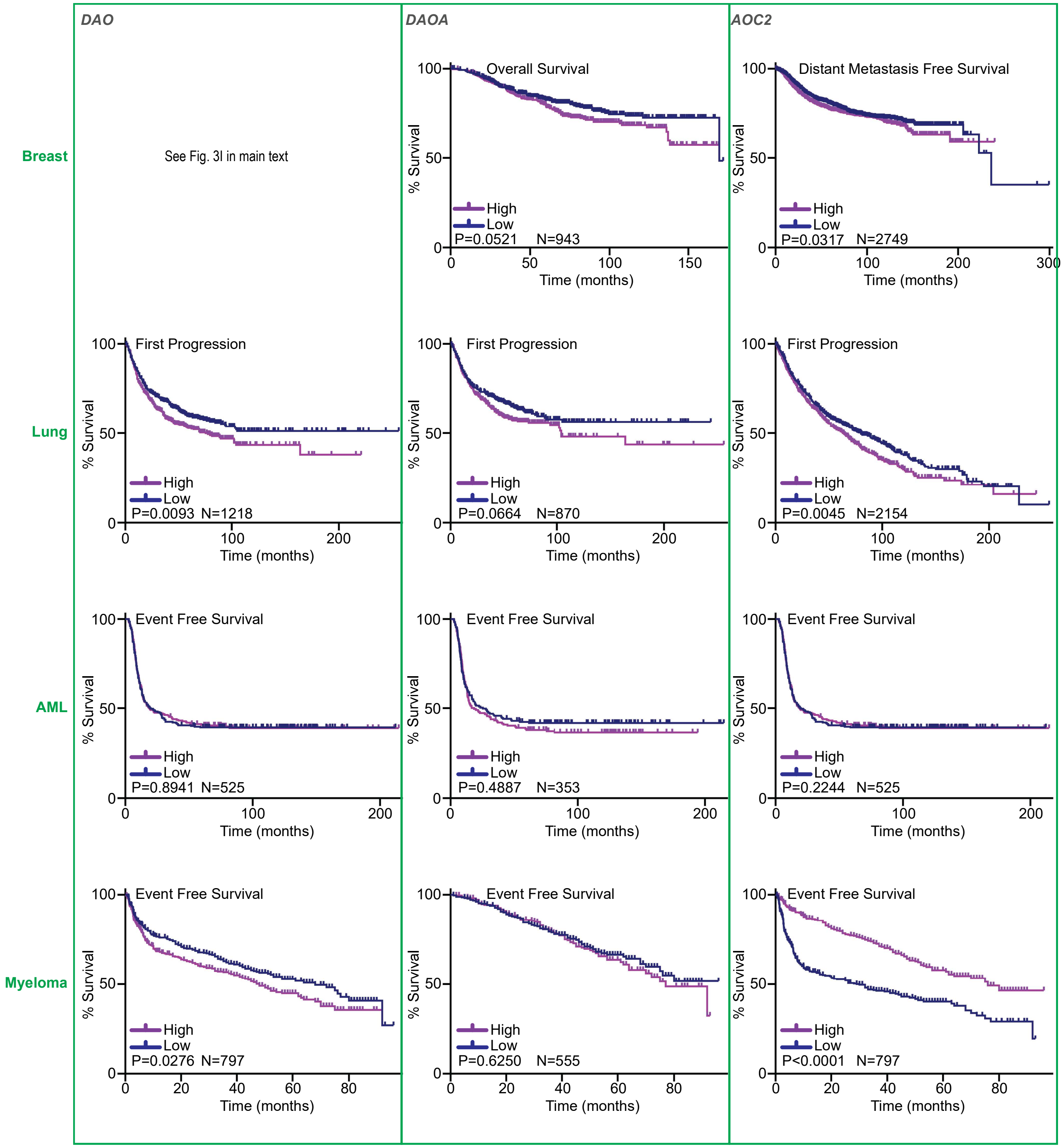
DAO
DAOA
AOC2
Breast
Lung
AML
Myeloma
See Fig. 3I in main text
Overall Survival
High
Low
P=0.0521 N=943
% Survival
Time (months)
Distant Metastasis Free Survival
P=0.0317 N=2749
First Progression
P=0.0093 N=1218
P=0.0664 N=870
P=0.0045 N=2154
Event Free Survival
P=0.8941 N=525
P=0.4887 N=353
P=0.2244 N=525
P=0.0276 N=797
P=0.6250 N=555
P<0.0001 N=797

SFig 11

**Supplementary Figure 11:** ***Expression of enzymes involved in histamine catabolism are associated with poor survival in different tumor types.*** Tumor types are by row, denoted on the left, and gene is by column, denoted on the top. Data accessed from the Kaplan-Meier Plotter webtool based on data from GEO, EGA, and TCGA (P values indicated from Logrank test, with the exception of AOC2-Breast and DAO-AML which used the Gehan Breslow Wilcoxon test). Sample numbers (N) are indicated on each graph.

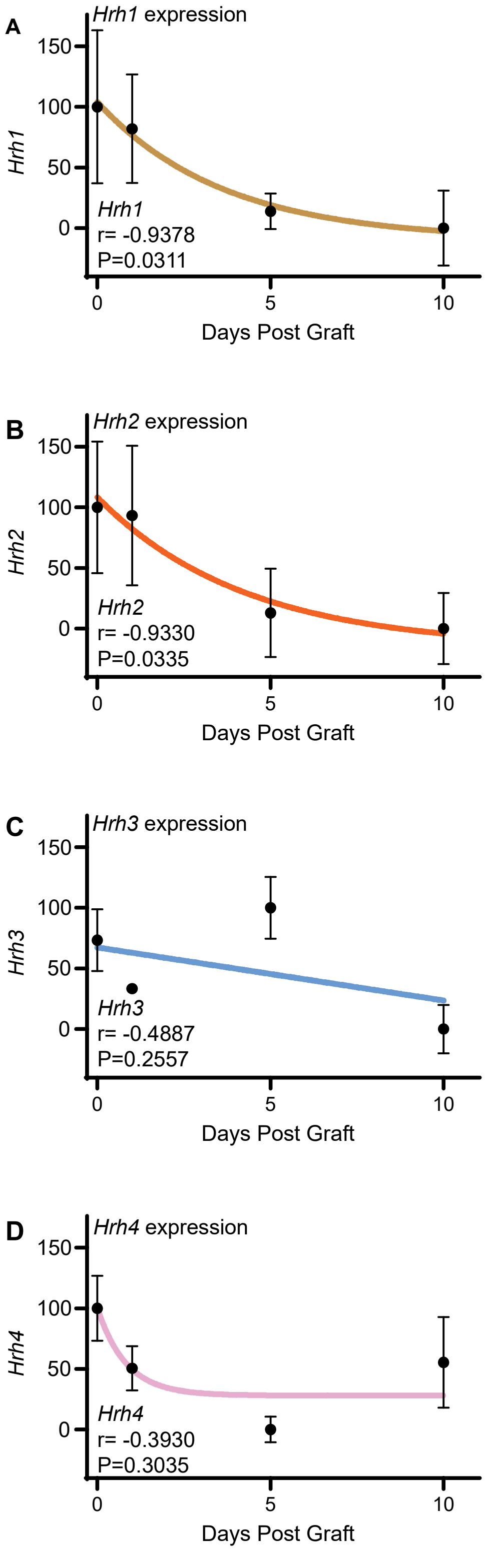

A
Hrh1 expression
150
100
50
0
Hrh1
Hrh1
r= -0.9378
P=0.0311
0
5
10
Days Post Graft
B
Hrh2 expression
150
100
50
0
Hrh2
Hrh2
r= -0.9330
P=0.0335
0
5
10
Days Post Graft
C
Hrh3 expression
150
100
50
0
Hrh3
Hrh3
r= -0.4887
P=0.2557
0
5
10
Days Post Graft
D
Hrh4 expression
150
100
50
0
Hrh4
Hrh4
r= -0.3930
P=0.3035
0
5
10
Days Post Graft


SFig 12

**Supplementary Figure 12:** ***Expression of $H_2R$ in 4T1 metastatic lesions decreases through time.*** 4T1 cells were grafted intravenously. Mice were euthanized at indicated timepoints, and lungs assessed for mRNA content by qPCR. Data was fit to a one-phase decay regression model. Pearson's one-way correlation was assessed with r and P values indicated. **(A)** $H_1R$ (*Hrh1*, N=17). **(B)** $H_2R$ (*Hrh2*, N=17). **(C)** $H_3R$ (*Hrh3*, N=17). **(D)** $H_4R$ (*Hrh4*, N=17).

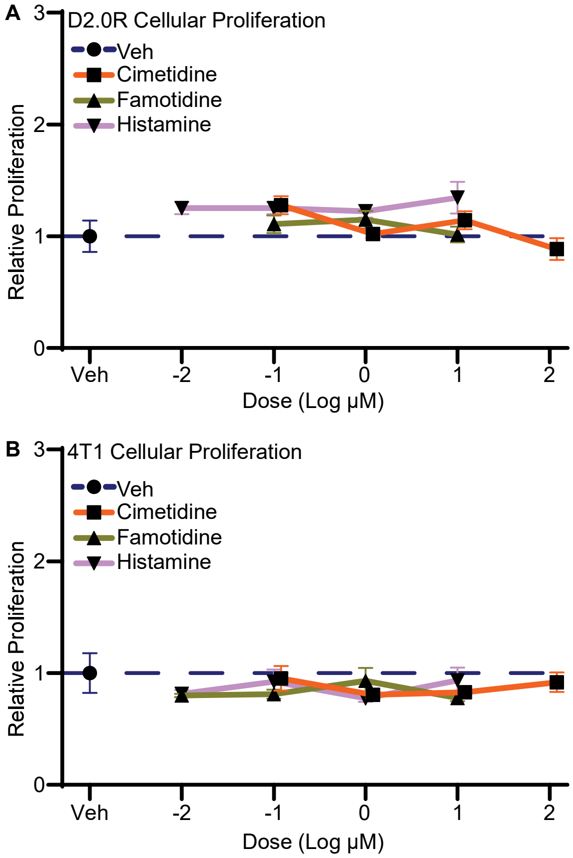
A
D2.0R Cellular Proliferation
Veh
Cimetidine
Famotidine
Histamine
Relative Proliferation
0
1
2
3
Veh
-2
-1
0
1
2
Dose (Log µM)
B
4T1 Cellular Proliferation
Veh
Cimetidine
Famotidine
Histamine
Relative Proliferation
0
1
2
3
Veh
-2
-1
0
1
2
Dose (Log µM)

SFig 13

**Supplementary Figure 13:** ***Antagonists to $H_2R$ do not have significant effects on proliferation of D2.0R or 4T1 cells***. **(A)** D2.0R cells were plated on day 0. On day 1 they were treated with vehicle (DMSO) or ligands at the indicated doses. On Day 4 the relative cell abundance was assessed by total DNA content (as described in [131]). **(B)** 4T1 cells were treated as in (A) and assessed for total DNA content. Values are relative to vehicle (D2.0R N=58; 4T1 N=65).

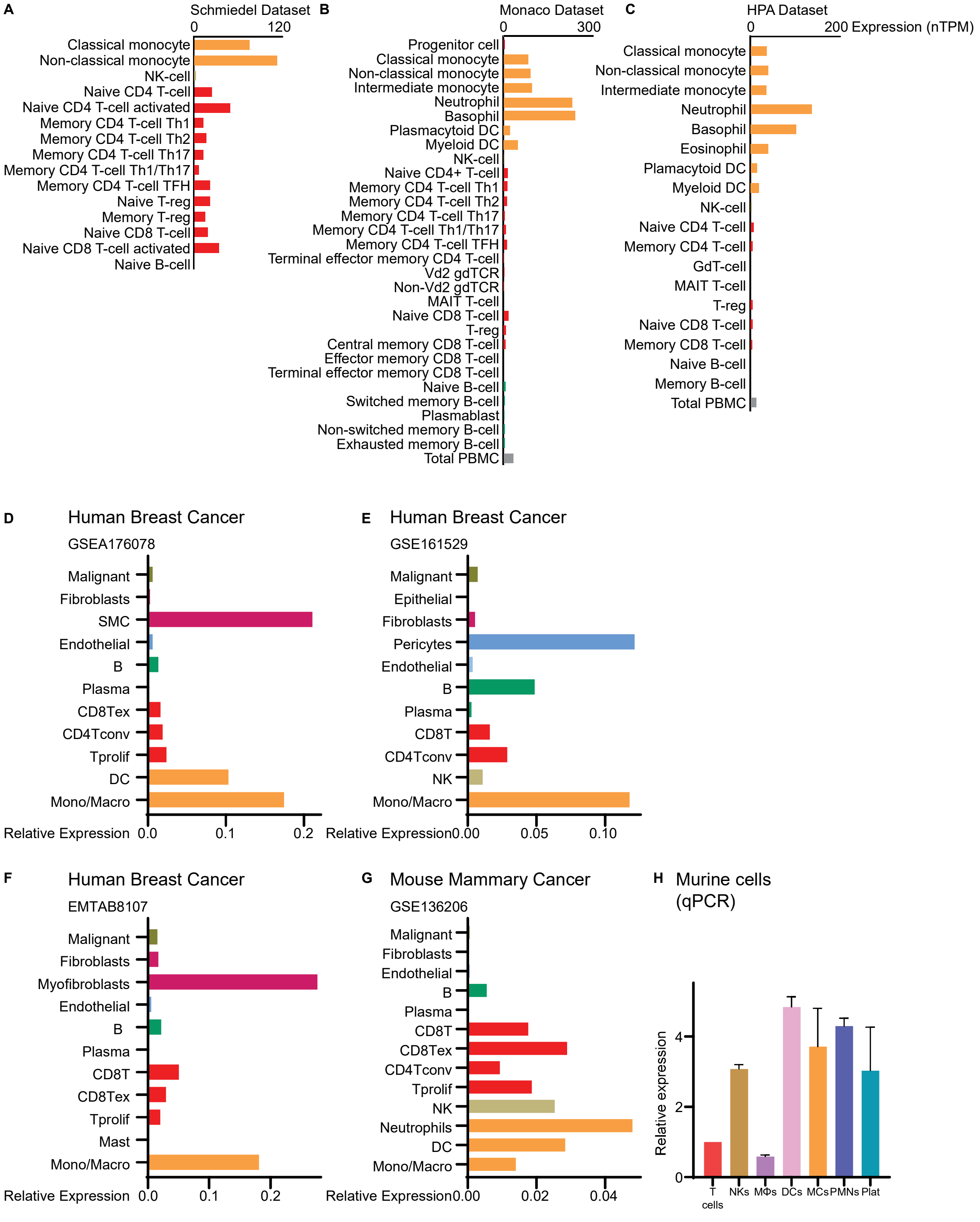
A
Schmiedel Dataset
0
120
Classical monocyte
Non-classical monocyte
NK-cell
Naive CD4 T-cell
Naive CD4 T-cell activated
Memory CD4 T-cell Th1
Memory CD4 T-cell Th2
Memory CD4 T-cell Th17
Memory CD4 T-cell Th1/Th17
Memory CD4 T-cell TFH
Naive T-reg
Memory T-reg
Naive CD8 T-cell
Naive CD8 T-cell activated
Naive B-cell
B
Monaco Dataset
0
300
Progenitor cell
Classical monocyte
Non-classical monocyte
Intermediate monocyte
Neutrophil
Basophil
Plasmacytoid DC
Myeloid DC
NK-cell
Naive CD4+ T-cell
Memory CD4 T-cell Th1
Memory CD4 T-cell Th2
Memory CD4 T-cell Th17
Memory CD4 T-cell Th1/Th17
Memory CD4 T-cell TFH
Terminal effector memory CD4 T-cell
Vd2 gdTCR
Non-Vd2 gdTCR
MAIT T-cell
Naive CD8 T-cell
T-reg
Central memory CD8 T-cell
Effector memory CD8 T-cell
Terminal effector memory CD8 T-cell
Naive B-cell
Switched memory B-cell
Plasmablast
Non-switched memory B-cell
Exhausted memory B-cell
Total PBMC
C
HPA Dataset
0
200 Expression (nTPM)
Classical monocyte
Non-classical monocyte
Intermediate monocyte
Neutrophil
Basophil
Eosinophil
Plamacytoid DC
Myeloid DC
NK-cell
Naive CD4 T-cell
Memory CD4 T-cell
GdT-cell
MAIT T-cell
T-reg
Naive CD8 T-cell
Memory CD8 T-cell
Naive B-cell
Memory B-cell
Total PBMC
D
Human Breast Cancer
GSEA176078
Malignant
Fibroblasts
SMC
Endothelial
B
Plasma
CD8Tex
CD4Tconv
Tprolif
DC
Mono/Macro
Relative Expression
0.0
0.1
0.2
E
Human Breast Cancer
GSE161529
Malignant
Epithelial
Fibroblasts
Pericytes
Endothelial
B
Plasma
CD8T
CD4Tconv
NK
Mono/Macro
Relative Expression
0.00
0.05
0.10
F
Human Breast Cancer
EMTAB8107
Malignant
Fibroblasts
Myofibroblasts
Endothelial
B
Plasma
CD8T
CD8Tex
Tprolif
Mast
Mono/Macro
Relative Expression
0.0
0.1
0.2
G
Mouse Mammary Cancer
GSE136206
Malignant
Fibroblasts
Endothelial
B
Plasma
CD8T
CD8Tex
CD4Tconv
Tprolif
NK
Neutrophils
DC
Mono/Macro
Relative Expression
0.00
0.02
0.04
H
Murine cells
(qPCR)
Relative expression
0
2
4
T cells
NKs
MΦs
DCs
MCs
PMNs
Plat

SFig 14

**Supplementary Figure 14:** ***Single Cell RNA-sequencing of human and murine samples indicates elevated expression of $H_2R$ in myeloid immune cells.*** **(A-C)** Assessment of *HRH2* ($H_2R$ mRNA) expression by scRNA-seq circulating immune cells across three different datasets (data obtained from Human Protein Atlas; proteinatlas.org[148]). **(D-G)** scRNA-seq of breast/mammary tumors indicates *HRH2* expression across several different immune cells. Data was obtained from the indicated databases and accessed through TISCH2 [149]. GSE176078: 26 tumors, 89,471 cells. GSE161529: 52 tumors, 332,168 cells. EMTAB8107: 14 tumors, 33,043 cells. GSE136206: 27,352 cells. **(H)** *Hrh2* expression in different isolated murine cell types and platelets, as determined by qPCR. Expression was normalized to cyclophilin.

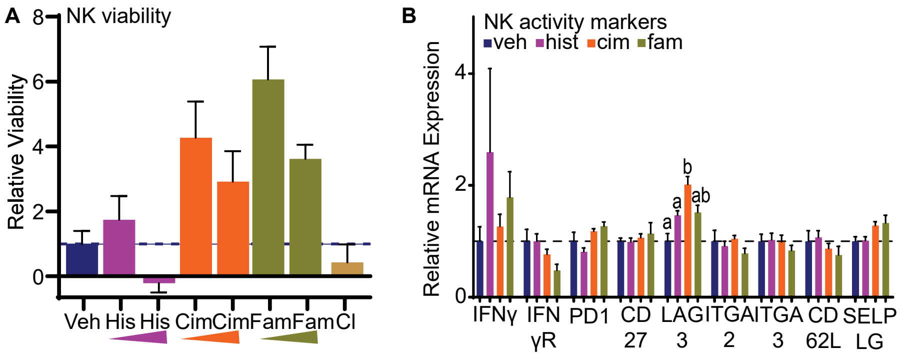

A
NK viability
Relative Viability
8
6
4
2
0
Veh
His
His
Cim
Cim
Fam
Fam
Cl
B
NK activity markers
veh
hist
cim
fam
Relative mRNA Expression
4
2
0
a
a
b
ab
IFNγ
IFN γR
PD1
CD 27
LAG 3
ITGA 2
ITGA 3
CD 62L
SELP LG


SFig 15

**Supplementary Figure 15:** ***$H_2R$ antagonists do not have significant effects on NK cell viability or markers of activity.*** **(A)** Splenic NK cells were cultured in the presence of the indicated ligands at increasing concentrations (histamine: 1 or 2μM, cimetidine: 12 or 24μM, famotidine: 1 or 2μM, clemastine: 240nM) for 24h. Cell viability was assessed by reduction of resazurin to resorufin (as described[133,134]). **(B)** Markers of NK cell activation were assessed by qPCR after treatment with the indicated ligands for 24h. Statistical differences, if any, are denoted by different letters (N=21-23 per gene analysis, 1-Way ANOVA followed by multiple comparison test of geometric means with Šidák's correction).

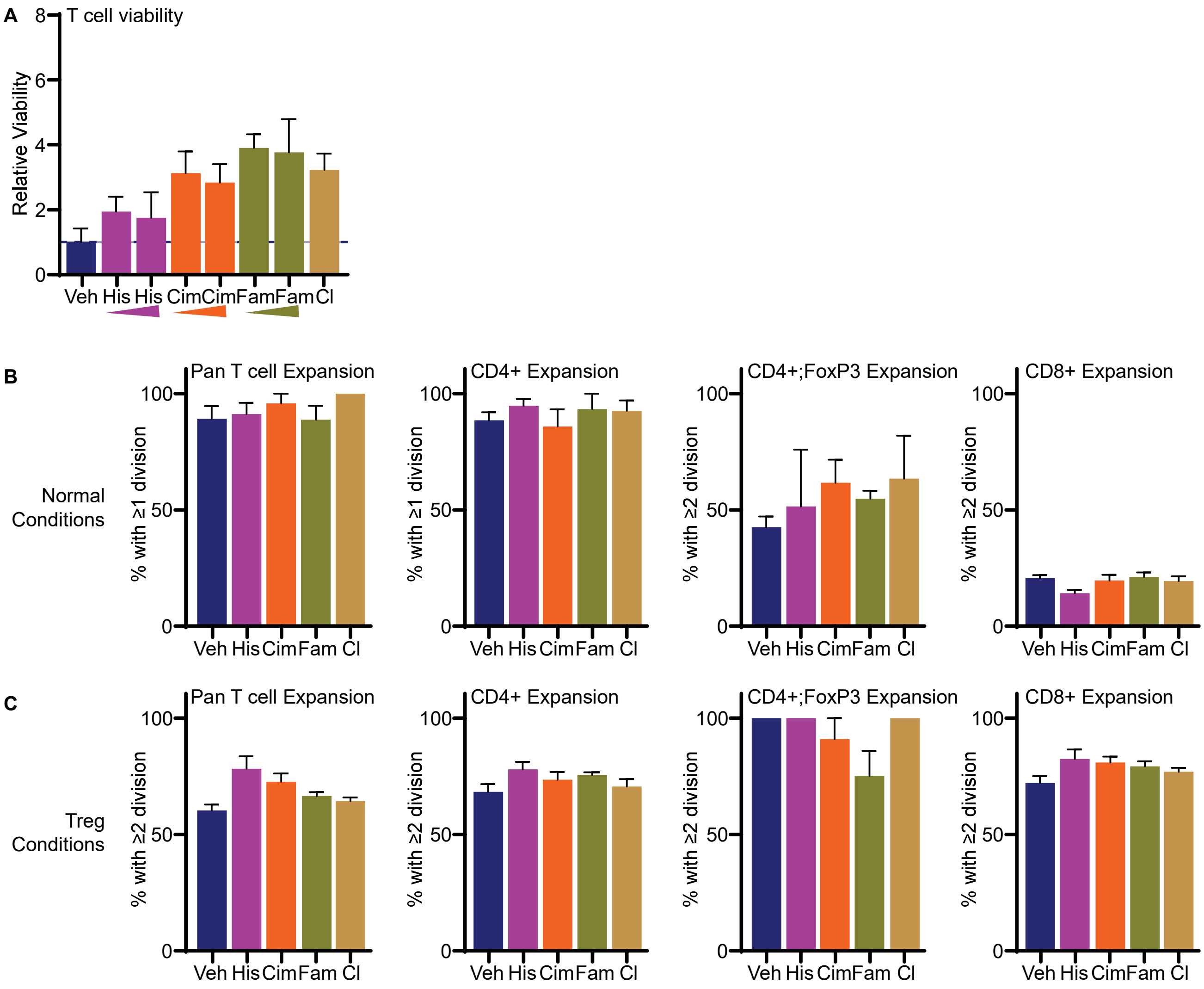

A
T cell viability
Relative Viability
Veh His His Cim Cim Fam Fam Cl
B
Normal Conditions
Pan T cell Expansion
CD4+ Expansion
CD4+;FoxP3 Expansion
CD8+ Expansion
% with ≥1 division
% with ≥2 division
Veh His Cim Fam Cl
C
Treg Conditions
Pan T cell Expansion
CD4+ Expansion
CD4+;FoxP3 Expansion
CD8+ Expansion
% with ≥2 division
Veh His Cim Fam Cl


SFig 16

**Supplementary Figure 16:** ***$H_2R$ antagonists do not have significant effects on T cell viability, markers of activity or ability to expand.*** **(A)** Splenic T cells were cultured in the presence of indicated ligands at increasing concentrations (histamine: 1 or 2µM, cimetidine: 12 or 24µM, famotidine: 1 or 2µM, clemastine: 240nM) for 24h. Cell viability was assessed by reduction of resazurin to resorufin. **(B)** T cell expansion. CFSE-stained T cells were incubated with indicated ligands and activated with antibodies against CD3 and CD28. T cell proliferation was assessed with flow cytometry for dilution of CFSE signal. From left to right: pan T cells, CD4+ only, CD4+;FoxP3+ (Treg) only, and CD8+ only. **(C)** CFSE-stained T cells were cultured in media to support differentiation into Tregs (includes IL2, TGFβ and β-mercaptoethanol), and activated with antibodies against CD3 and CD8. T cell proliferation was assessed with flow cytometry for dilution of CFSE signal. From left to right: pan T cells, CD4+ only, CD4+;FoxP3+ (Treg) only, and CD8+ only.

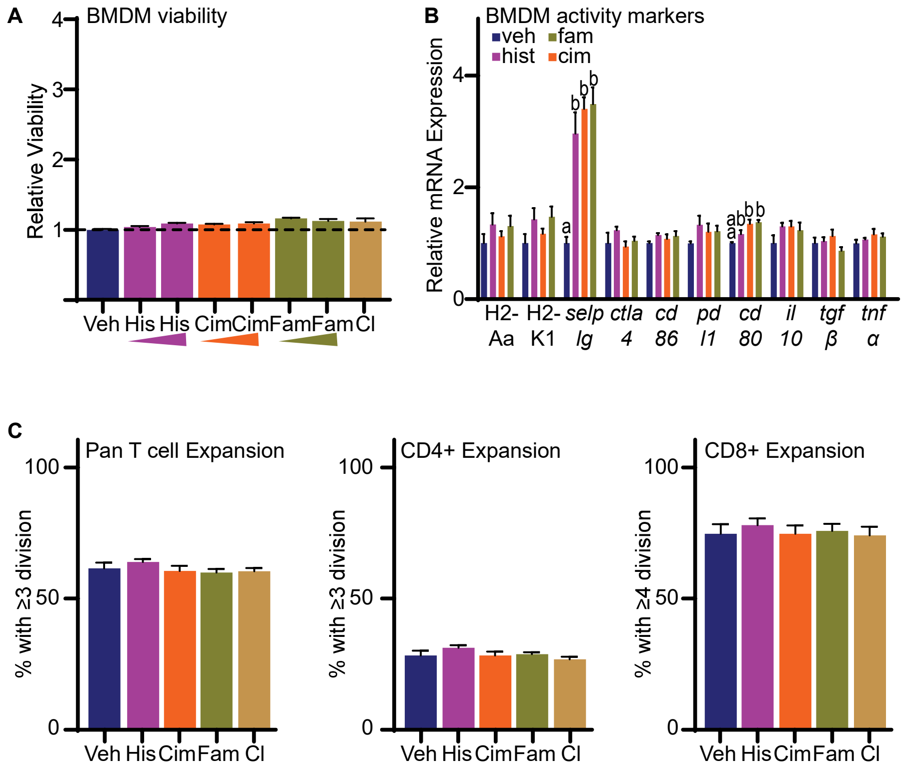
A
BMDM viability
Relative Viability
4
3
2
1
Veh His His Cim Cim Fam Fam Cl
B
BMDM activity markers
veh
fam
hist
cim
Relative mRNA Expression
4
2
0
a
b
b
b
a
ab
b
b
H2- Aa
H2- K1
selp lg
ctla 4
cd 86
pd l1
cd 80
il 10
tgf β
tnf α
C
Pan T cell Expansion
% with ≥3 division
100
50
0
Veh His Cim Fam Cl
CD4+ Expansion
% with ≥3 division
100
50
0
Veh His Cim Fam Cl
CD8+ Expansion
% with ≥4 division
100
50
0
Veh His Cim Fam Cl

SFig 17

**Supplementary Figure 17:** ***$H_2R$ antagonists do not have significant effects on macrophage markers of activity or ability to support T cell expansion*** **(A)** Bone marrow derived macrophages were cultured in the presence of indicated ligands at increasing concentrations (histamine: 1 or 2μM, cimetidine: 12 or 24μM, famotidine: 1 or 2μM, clemastine: 240nM) for 24h. Cell viability was assessed by reduction of resazurin to resorufin. **(B)** Markers of macrophage activation were assessed by qPCR after treatment of BMDMs with the indicated ligands for 24h. Statistical differences, if any, are denoted by different letters (N=16 per gene analysis, 1-Way ANOVA followed by multiple comparison test of geometric means with Šidák's correction). **(C)** Bone marrow derived macrophages were pre-treated with indicated ligands, washed and then co-cultured with CFSE-stained and activated T cells. T cell proliferation was assessed with flow cytometry for dilution of CFSE signal. From left to right: pan T cells, CD4+ only, and CD8+ only (no statistically significant differences, N=25).

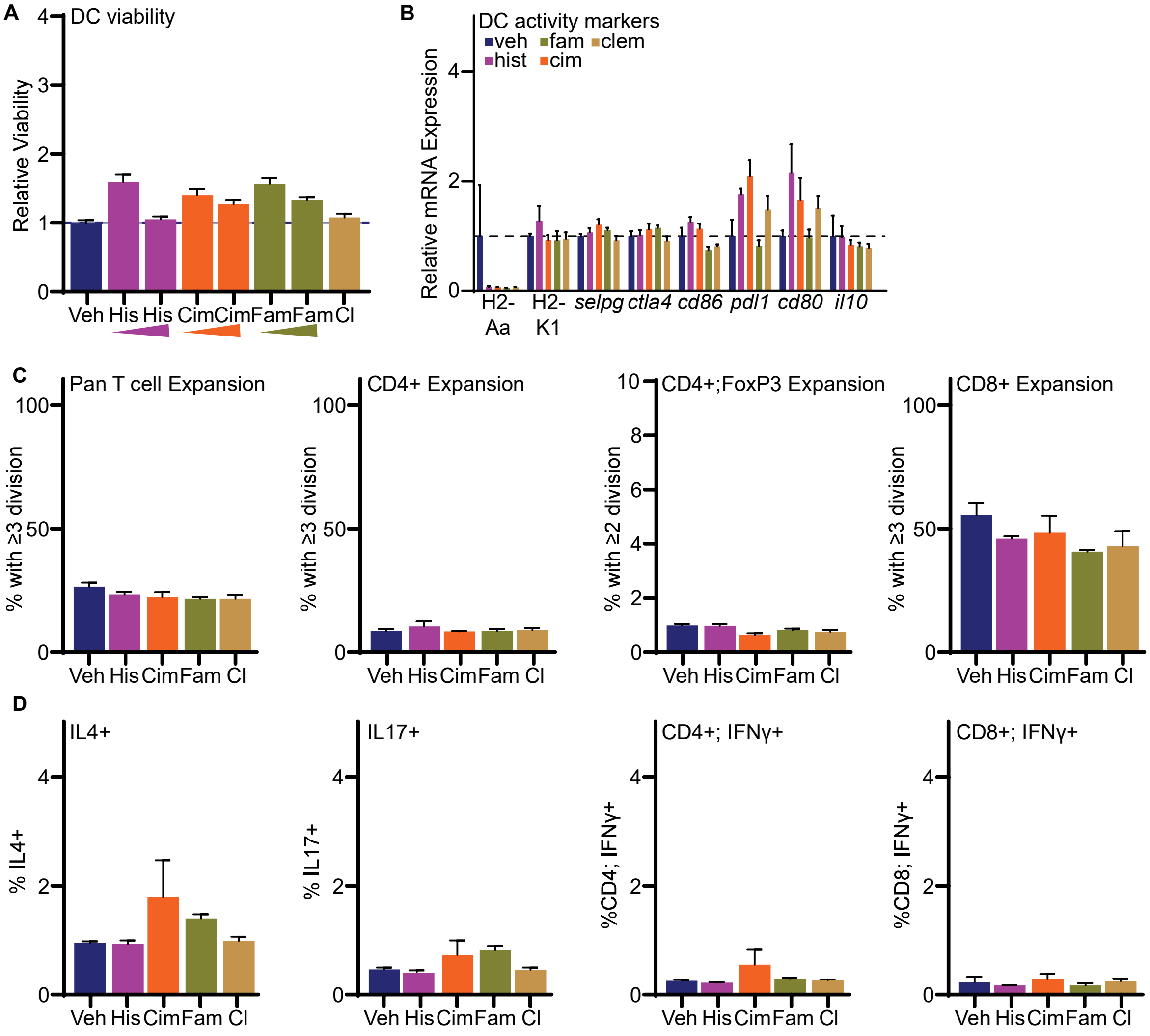
A
DC viability
Relative Viability
Veh His His Cim Cim Fam Fam Cl
B
DC activity markers
veh fam clem
hist cim
Relative mRNA Expression
H2-Aa
H2-K1
selpg
ctla4
cd86
pdl1
cd80
il10
C
Pan T cell Expansion
CD4+ Expansion
CD4+;FoxP3 Expansion
CD8+ Expansion
% with ≥3 division
% with ≥2 division
Veh His Cim Fam Cl
D
IL4+
IL17+
CD4+; IFNγ+
CD8+; IFNγ+
% IL4+
% IL17+
%CD4; IFNγ+
%CD8; IFNγ+
Veh His Cim Fam Cl

SFig 18

**Supplementary Figure 18:** ***$H_2R$ antagonists do not have significant effects on dendritic cell (DC) viability, markers of activity, or ability to support T cell expansion.*** **(A)** DCs were cultured in the presence of indicated ligands at increasing concentrations (histamine: 1 or 2μM, cimetidine: 12 or 24μM, famotidine: 1 or 2μM, clemastine: 240nM) for 24h. Cell viability was assessed by reduction of resazurin to resorufin. **(B)** Markers of DC activation were assessed by qPCR after treatment with the indicated ligands for 24h. **(C)** DCs were pre-treated with indicated ligands, washed and then co-cultured with CFSE-stained and activated T cells. T cell proliferation was assessed with flow cytometry for dilution of CFSE signal. From left to right: pan T cells, CD4+ only, CD4+;FoxP3+ (Treg) only, and CD8+ only. **(D)** Resulting expanded T cells were further assessed for markers including (from left to right): IL4, IL17, IFNγ in CD4+ cells, and IFNγ in CD8+ cells.

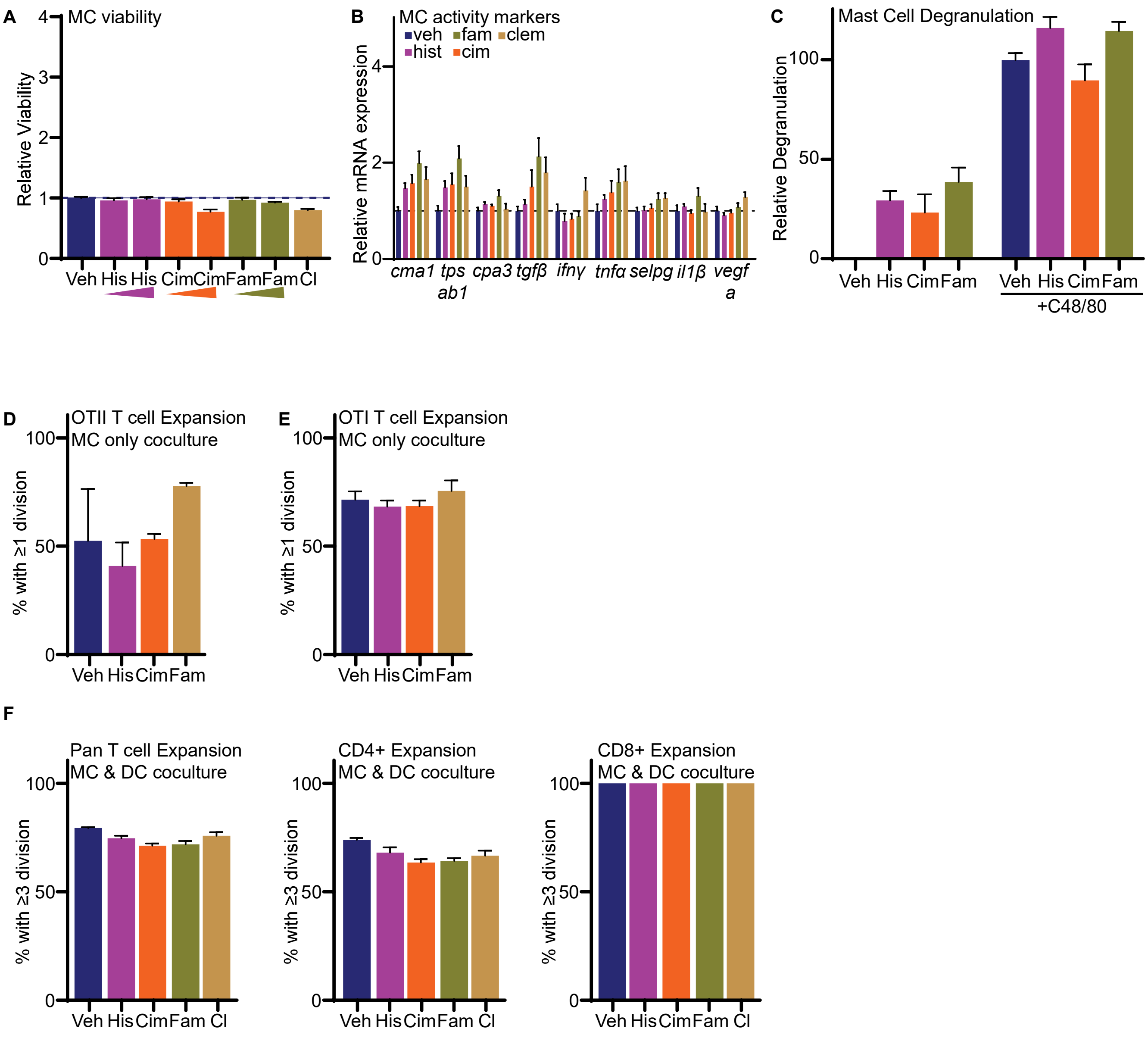
A
MC viability
Relative Viability
Veh His His Cim Cim Fam Fam Cl
B
MC activity markers
veh fam clem
hist cim
Relative mRNA expression
cma1 tpsab1 cpa3 tgfβ ifnγ tnfα selpg il1β vegfa
C
Mast Cell Degranulation
Relative Degranulation
Veh His Cim Fam Veh His Cim Fam
+C48/80
D
OTII T cell Expansion
MC only coculture
% with ≥1 division
Veh His Cim Fam
E
OTI T cell Expansion
MC only coculture
% with ≥1 division
Veh His Cim Fam
F
Pan T cell Expansion
MC & DC coculture
% with ≥3 division
Veh His Cim Fam Cl
CD4+ Expansion
MC & DC coculture
% with ≥3 division
Veh His Cim Fam Cl
CD8+ Expansion
MC & DC coculture
% with ≥3 division
Veh His Cim Fam Cl

SFig 19

**Supplementary Figure 19:** ***$H_2R$ antagonists do not have significant effects on mast cell (MC) viability, markers of activity, degranulation, or ability to present antigen and activate T cells.*** **(A)** MCs were cultured in the presence of indicated ligands at increasing concentrations (histamine: 1 or 2μM, cimetidine: 12 or 24μM, famotidine: 1 or 2μM, clemastine: 240nM) for 24h. Cell viability was assessed by reduction of resazurin to resorufin. **(B)** Markers of MC activation were assessed by qPCR after treatment with the indicated ligands for 24h. **(C)** MCs were treated with indicated ligands in the absence or presence of the degranulation inducer, C48/80. Degranulation was assessed by enzymatic activity of β- hexosaminidase. **(D)** CFSE-stained OTII T cells were cultured in the presence of MCs previously pulsed with OVA. Subsequent T cell proliferation was assessed with flow cytometry for dilution of CFSE signal. OTII T cell expansion after OVA antigen presentation is predominantly CD4+. **(E)** CFSE-stained OTI T cells were cultured in the presence of MCs previously pulsed with OVA. Subsequent T cell proliferation was assessed with flow cytometry for dilution of CFSE signal. OTI T cell expansion after OVA antigen presentation is predominantly CD8+. **(F)** DCs and MCs were co-cultured. Pre-treated DCs and MCs were then co-cultured with CFSE-stained and activated T cells. T cell proliferation was assessed with flow cytometry for dilution of CFSE signal. From left to right: pan T cells, CD4+ only, and CD8+ only.

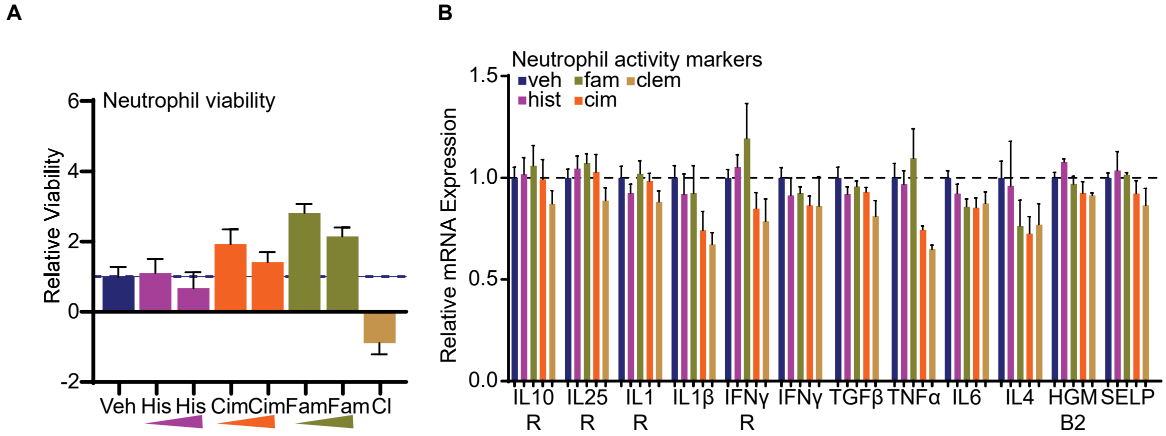
A
B
Neutrophil viability
Relative Viability
6
4
2
0
-2
Veh
His
His
Cim
Cim
Fam
Fam
Cl
Neutrophil activity markers
veh
fam
clem
hist
cim
Relative mRNA Expression
1.5
1.0
0.5
0.0
IL10 R
IL25 R
IL1 R
IL1β
IFNγ R
IFNγ
TGFβ
TNFα
IL6
IL4
HGM B2
SELP

SFig 20

**Supplementary Figure 20:** ***$H_2R$ antagonists do not have significant effects on neutrophil viability or markers of activity.*** **(A)** Neutrophils were cultured in the presence of indicated ligands at increasing concentrations (histamine: 1 or 2μM, cimetidine: 12 or 24μM, famotidine: 1 or 2μM, clemastine: 240nM) for 24h. Cell viability was assessed by reduction of resazurin to resorufin. **(B)** Markers of neutrophil activity were assessed by qPCR after treatment with the indicated ligands for 24h.

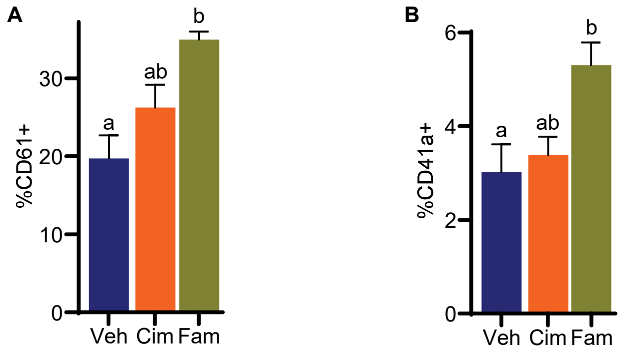


SFig 21

**Supplementary Figure 21:** ***Increased presence of activated platelets and NETs after chronic treatment with $H_2R$ antagonists.*** **(A)** Flow cytometric analysis for CD61+ or **(B)** CD41a+ events, both markers of platelets. Different letters denote statistical significance (1-Way ANOVA followed by multiple comparison test with Šidák's correction, N=43).

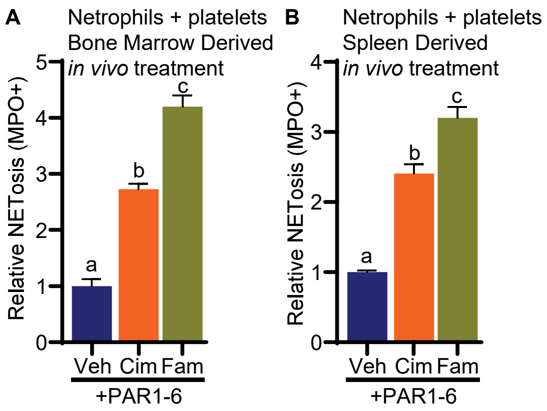
A
Netrophils + platelets
Bone Marrow Derived
*in vivo* treatment
Relative NETosis (MPO+)
0
1
2
3
4
5
a
b
c
Veh
Cim
Fam
+PAR1-6
B
Netrophils + platelets
Spleen Derived
*in vivo* treatment
Relative NETosis (MPO+)
0
1
2
3
4
a
b
c
Veh
Cim
Fam
+PAR1-6

SFig 22

**Supplementary Figure 22:** ***Cimetidine or famotidine increase NETs as determined by flow cytometry for MPO+ events***. Flow cytometric analysis of MPO+ events after a co-culture of platelets and neutrophils from mice chronically treated with the indicated ligands. Platelets and neutrophils were derived from either **(A)** the bone marrow, or **(B)** the spleen. Different letters denote statistical significance (1-Way ANOVA followed by multiple comparison test with Šidák's correction).

**Supplementary Table 1A:** ***Peak identification of the 36 mix standard fatty acid methyl ester from GC-MS analysis***. 36 mixed fatty acid methyl esters (FAMEs) analyzed by GC-MS on an HP-88 column (60 m × 0.25 mm, 0.20 µm). The oven temperature program was as follows: initial temperature of 60 °C for 1 min, ramped at 10 °C/min to 145 °C, then at 1 °C/min to 190 °C, and finally at 5 °C/min to 220 °C. Mass spectrometry was operated in scan mode over an m/z range of 50–500 amu with an EI voltage of 70 eV. Compound identification is based on peaks shown in **SFig2B**.

| **Peak NO.** | **Compound** | **Abbreviation** | **$R_t$ (min)** | **MW** |
|---|---|---|---|---|
| 1 | Butyric | C4:0 | 5.194 | 102 |
| 2 | Caproic | C6:0 | 6.946 | 130 |
| 3 | Caprylic | C8:0 | 9.027 | 158 |
| 4 | Capric | C10:0 | 11.172 | 186 |
| 5 | Undecanoic | C11:0 | 12.371 | 200 |
| 6 | Lauric | C12:0 | 13.745 | 214 |
| 7 | Tridecanoic | C13:0 | 15.368 | 228 |
| 8 | Myristic | C14:0 | 17.334 | 242 |
| 9 | Myristoleic | C14:1 | 18.939 | 240 |
| 10 | Pentadecanoic | C15:0 | 19.693 | 256 |
| 11 | cis-10-Petadecanoic | C15:1 | 21.624 | 254 |
| 12 | Palmitic | C16:0 | 22.523 | 270 |
| 13 | Palmitoleic | C16:1 | 24.322 | 268 |
| 14 | Heptadecanoic | C17:0 | 25.791 | 284 |
| 15 | cis-10-Heptadecenoic | C17:1 | 27.854 | 282 |
| 16 | Stearic | C18:0 | 29.524 | 298 |
| 17 | Elaidic | C18:1 n9t | 30.820 | 296 |
| 18 | Oleic | C18:1 n9c | 31.438 | 296 |
| 19 | Linolelaidic | C18:2 n6t | 33.297 | 294 |
| 20 | Linoleic | C18:2 n6c | 34.736 | 294 |
| 21 | γ-Linolenic | C18:3 n6 | 37.143 | 292 |
| 22 | Arachidic | C20:0 | 37.990 | 326 |
| 23 | Linolenic | C18:3 n3 | 38.855 | 292 |
| 24 | *cis*-11-Eicosenoic | C20:1 | 40.101 | 324 |
| 25 | Heneicosanoic | C21:0 | 42.542 | 340 |
| 26 | *cis*-11,14-Eicosadienoic | C20:2 | 43.803 | 322 |
| 27 | *cis*-8,11,14-Eicosatrienoic | C20:3 n6 | 46.353 | 320 |

| | | | | |
|---|---|---|---|---|
| 28 | Behenic | C22:0 | 47.252 | 354 |
| 29 | *cis*-11,14,17-Eicosatrienoic | C20:3 n3 | 48.224 | 320 |
| 30 | Erucic | C22:1 n9 | 49.478 | 352 |
| 31 | Tricosanoic | C23:0 | 51.958 | 368 |
| 32 | Methyl-cis-5,8,11,14-Eicosatetraenoic | C20:4 n6 | 52.859 | 318 |
| 33 | *cis*-13,16-Docosadienoic | C22:2 | 53.332 | 350 |
| 34 | Lignoceric | C24:0 | 56.415 | 382 |
| 35 | Nervonic | C24:1 | 57.978 | 380 |
| 36 | *cis*-4,7,19,13,16,19-Docosahexaenoic | C22:6 n3 | 61.125 | 342 |

**Supplementary Table 1B.** Total ion chromatogram of cfBF analyzed by GC-MS on an HP-88 column (60 m × 0.25 mm, 0.20 µm). Compound identification was based on the NIST Mass Spectral Library and Wiley Registry of Mass Spectral Data libraries, 36 mixed FAME reference standards, and other commercially available standards, including vaccenic acid and dihomo-γ-linolenic acid. Compound identification is indicated in in table below

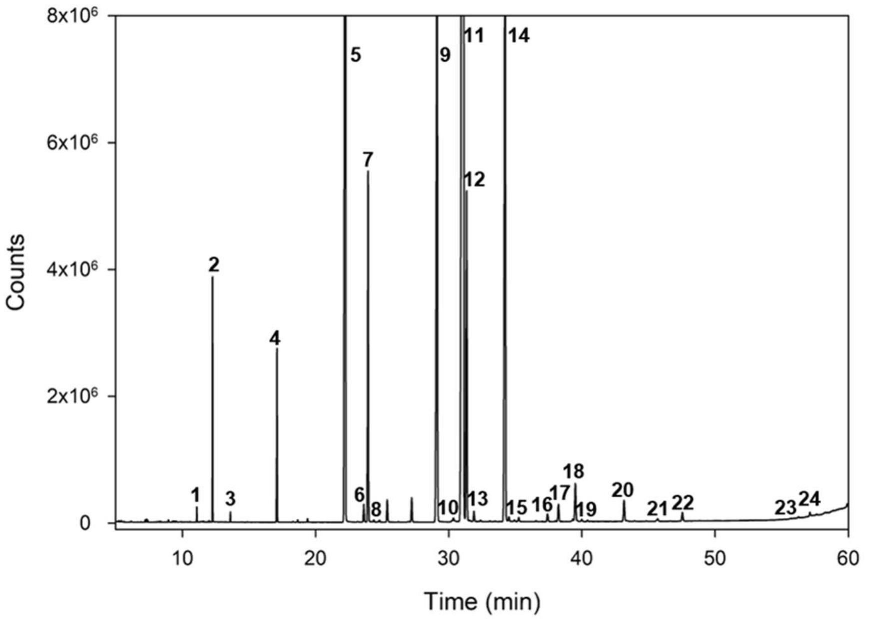


| Peak | Identification | RT (min) |
|---|---|---|
| 1 | Decanoic C10:0 | 11.082 |
| 2 | C11:0 (IS)* | 12.259 |
| 3 | Tridecylic C12:0 | 13.6 |
| 4 | Myristic C14:0 | 17.094 |
| 5 | Palmitic C16:0 | 22.232 |
| 6 | C16:1-7c | 23.615 |
| 7 | Palmitoleic C16:1-9c | 23.936 |
| 8 | C16:1-11c | 24.365 |
| 9 | Stearic C18:0 | 29.113 |
| 10 | Elaidic C18:1 n9t | 30.347 |
| 11 | Oleic C18:1 n9c | 31.092 |
| 12 | Vaccenic C18:1 n11 | 31.336 |
| 13 | C18:1 n13 | 31.888 |
| 14 | Linoleic C18:2 n6c | 34.235 |
| 15 | Linolenic C18:3 n6 | 34.522 |
| 16 | Arachidic C20:0 | 37.42 |
| 17 | Linolenic C18:3 n3 | 38.243 |
| 18 | Gondoic C20:1 11c | 39.317 |
| 19 | Paullinic C20:1 13c | 39.506 |
| 20 | C20:2 | 43.167 |
| 21 | Dihomo-γ-linolenic C20:3 n6 | 45.685 |
| 22 | Mead C20:3;<br>Arachidonic C20:4 | 47.543 |
| 23 | Eicosapentaenoic C20:5 n3 | 52.634 |
| 24 | Nervonic C24:1 | 57.117 |

* IS: internal standard

**Supplementary Table 2:** ***Fatty acid composition of lard, from cured fried bacon (cfBF) and fried bacon.*** Cured bacon was pan-fried as described in the main text. Soxhlet extraction was performed on rendered lard, the dried bacon post-frying and the oil resulting from the pan-frying of bacon (cfBF). [FAME: fatty acid methyl ester]

| **Lipid Species** | **Lard FAME%** | **cfBF FAME%** | **Fried Bacon Fat FAME%** |
|---|---|---|---|
| C10:0 | 0.05 ± 0.01 | 0.07 ± 0.01 | 0.07 ± 0.01 |
| C12:0 | 0.05 ± 0.01 | 0.06 ± 0.01 | 0.06 ± 0.01 |
| C14:0 | 1.09 ± 0.01 | 1.27 ± 0.01 | 1.25 ± 0.06 |
| C16:0 | 22.79 ± 0.07 | 23.27 ± 0.01 | 23.17 ± 0.27 |
| C16:1-7c | 0.25 ± 0.01 | 0.19 ± 0.01 | 0.19 ± 0.04 |
| C16:1-9c | 1.65 ± 0.01 | 3.78 ± 0.01 | 3.95 ± 0.59 |
| C16:1-11c | 0.03 ± 0.01 | 0.02 ± 0.01 | 0.02 ± 0.01 |
| C18:0 | 13.77 ± 0.37 | 10.55 ± 0.23 | 10.09 ± 0.68 |
| C18:1 ω9t | 0.04 ± 0.02 | 0.04 ± 0.01 | 0.04 ± 0.01 |
| C18:1 ω9c | 38.13 ± 0.26 | 43.60 ± 0.20 | 44.85 ± 0.33 |
| C18:1 ω11 | 2.35 ± 0.03 | 4.14 ± 0.04 | 4.46 ± 0.18 |
| C18:1 ω13 | 0.07 ± 0.01 | 0.13 ± 0.01 | 0.14 ± 0.01 |
| C18:2 ω6c | 17.49 ± 0.10 | 11.21 ± 0.02 | 10.10 ± 0.87 |
| C18:3 ω6 | 0.03 ± 0.01 | 0.06 ± 0.01 | 0.06 ± 0.01 |
| C20:0 | 0.16 ± 0.01 | 0.13 ± 0.01 | 0.13 ± 0.01 |
| C18:3 ω3 | 0.53 ± 0.01 | 0.26 ± 0.01 | 0.21 ± 0.01 |
| C20:1 11c | 0.05 ± 0.01 | 0.03 ± 0.01 | 0.03 ± 0.01 |
| C20:1 13c | 0.61 ± 0.03 | 0.61 ± 0.01 | 0.64 ± 0.03 |
| C20:2 | 0.56 ± 0.01 | 0.33 ± 0.01 | 0.33 ± 0.02 |
| C20:3 ω6 | 0.13 ± 0.10 | 0.15 ± 0.01 | 0.14 ± 0.01 |
| Total saturated | 37.91 ± 0.44 | 35.34 ± 0.24 | 34.77 ± 0.36 |
| Total monounsaturated | 43.23 ± 0.32 | 52.59 ± 0.23 | 54.37 ± 0.45 |
| Total polyunsaturated | 18.85 ± 0.11 | 12.07 ± 0.01 | 10.86 ± 0.81 |
| Total trans fat | 0.04 ± 0.02 | 0.04 ± 0.01 | 0.04 ± 0.01 |
| Total ω6 | 17.56 ± 0.11 | 11.32 ± 0.01 | 10.21 ± 0.83 |
| Total ω3 | 0.54 ± 0.01 | 0.26 ± 0.01 | 0.22 ± 0.01 |
| Ratio ω6-ω3 | 32.8 | 42.7 | 47.5 |

**Supplementary Table 3:** ***Cholesterol and oxysterol content of lard and fat from cured fried bacon (cfBF).*** Results are expressed in mg/g of sample for the cholesterol and μg/g of sample for oxysterols. Mean ± SD (N = 4 for lard, N = 5 for cfBF). An asterisk (*) denotes significant differences among samples ($P < 0.05$). Note that the LC-MS/MS technique employed did not have the resolution to distinguish 7α- and 7β- HC.

| **Sterol Species** | **Lard** | **cfBF** |
|---|---|---|
| 22RHC | 0.016 ± 0.004 | 0.023 ± 0.007 |
| 25HC | 0.175 ± 0.028 | 0.250 ± 0.043 |
| 27HC | 0.313 ± 0.041 | 0.234 ± 0.035 |
| 4βHC | 0.055 ± 0.010 | 0.103 ± 0.012 |
| 7α/β HC | 0.196 ± 0.003 | 0.771 ± 0.074 |
| **Cholesterol** | **0.826 ± 0.120** | **0.985 ± 0.137** |
| **Total oxysterols** | **0.700 ± 0.065** | **1.123 ± 0.275 *** |
| **Oxysterols% of total cholesterol** | **0.08%** | **0.11%** |
